# Automated *in situ* profiling of chromatin modifications resolves cell types and gene regulatory programs

**DOI:** 10.1101/418681

**Authors:** Derek H. Janssens, Steven J. Wu, Jay F. Sarthy, Michael P. Meers, Carrie H. Myers, James M. Olson, Kami Ahmad, Steven Henikoff

## Abstract

Our understanding of eukaryotic gene regulation is limited by the complexity of protein-DNA interactions that comprise the chromatin landscape and by inefficient methods for characterizing these interactions. We recently introduced CUT&RUN, an antibody-targeted nuclease-cleavage method that profiles DNA-binding proteins, histones and chromatin modifying proteins *in situ* with exceptional sensitivity and resolution. Here we describe an automated CUT&RUN platform and apply it to characterize the chromatin landscapes of human cell lines. We find that CUT&RUN profiles of histone modifications crisply demarcate active and repressed chromatin regions, and we develop a continuous metric to identify cell-type specific promoter and enhancer activities. We test the ability of automated CUT&RUN to profile frozen tumor samples, and find that our method readily distinguishes two diffuse midline gliomas by their subtype-specific gene expression programs. The easy, cost-effective workflow makes automated CUT&RUN an attractive tool for high-throughput characterization of cell types and patient samples.

---

Cells establish their distinct identities by altering activity of the cis-regulatory DNA elements that control gene expression^1,2^. Promoter elements lie near the 5’ transcriptional start sites (TSSs) of all genes, whereas distal *cis*-regulatory elements such as enhancers often bridge long stretches in the DNA to interact with select promoters and direct cell-type-specific gene expression^1,2^. Defects in the nuclear proteins that recognize these cis-regulatory elements underlie many human diseases that often manifest in specific tissues and cell-types^3-7^. To provide a reference for molecular diagnosis of patient samples, efforts are underway to generate a comprehensive atlas of cells in the human body^8,9^. Characterizing cell-type specific chromatin landscapes is essential for this atlas; however, technical limitations have prevented implementation of traditional approaches for genome-wide profiling of chromatin proteins on the scales necessary for this project.

Despite the growing awareness that epigenetic derangements underlie many human diseases^10^, very few methods for high-throughput profiling of epigenomic information are available. Realizing the clinical potential of epigenomic technologies requires robust, scalable approaches that can profile large numbers of patient samples in parallel. Chromatin immunoprecipitation with antigen-specific antibodies combined with massively parallel sequencing (ChIP-seq) has been used extensively for epigenome profiling, but this method is labor-intensive, prone to artifacts^11^, and requires high sequencing depth to distinguish weak signals from genomic background noise. The combination of these factors has prevented implementation of ChIP-seq in clinical laboratory settings. Recently, we introduced CUT&RUN as an alternative chromatin profiling technique that uses factor-specific antibodies to tether micrococcal nuclease (MNase) to genomic binding sites^12,13^. The targeted nuclease cleaves chromatin around the binding sites, and the released DNA is sequenced using standard library preparation techniques, resulting in efficient mapping of protein-DNA interactions. CUT&RUN has very low backgrounds, which greatly reduces sample amounts and sequencing costs required to obtain high-quality genome-wide profiles^12,14^.

Here we modify the CUT&RUN protocol to profile chromatin proteins and modifications in a 96-well format on a liquid handling robot. By applying this method to the H1 human embryonic stem cell (hESC) line and K562 leukemia cell line, we demonstrate AutoCUT&RUN can be used to identify cell-type specific promoter and enhancer activities, providing a means to quantitatively distinguish cell-types based on their unique gene regulatory programs. In addition, we show that this method is able to define chromatin features from frozen solid tumor samples, setting the stage to analyze typical clinical specimens. AutoCUT&RUN is ideal for high-throughput studies of chromatin-based gene regulation, allowing for examination of chromatin landscapes in patient samples and expanding the toolbox for epigenetic medicine.

## Results

### An automated platform for genome-wide profiling of chromatin proteins

To adapt CUT&RUN to an automated format we equipped a Beckman Biomek FX liquid handling robot for magnetic separation and temperature control (Fig. 1a). First, cells are bound to Concanavalin A-coated magnetic beads, allowing all subsequent washes to be performed by magnetic separation. Bead-bound samples are then incubated with antibodies, and up to 96 samples are arrayed in a plate (Fig. 1a). Successive washes, tethering of a proteinA-MNase fusion protein, cleavage of DNA, and release of cleaved chromatin fragments into the sample supernatant are performed on the Biomek (Supplementary Fig. 1a). A major stumbling block to automating epigenomics protocols is they typically require purification of small amounts of DNA prior to library preparation. To overcome this hurdle, we developed a method to polish the DNA ends in chromatin fragments for direct ligation of Illumina library adapters (Supplementary Fig. 1a). End-polishing and adapter ligation are performed on a separate thermocycler and deproteinated CUT&RUN libraries are purified on the Biomek using Ampure XP magnetic beads both before and after PCR enrichment. This AutoCUT&RUN protocol allows a single operator to generate up to 96 libraries in 2 days that are ready to be pooled and sequenced (Fig. 1a) (Appendix 1).

**Figure 1:**
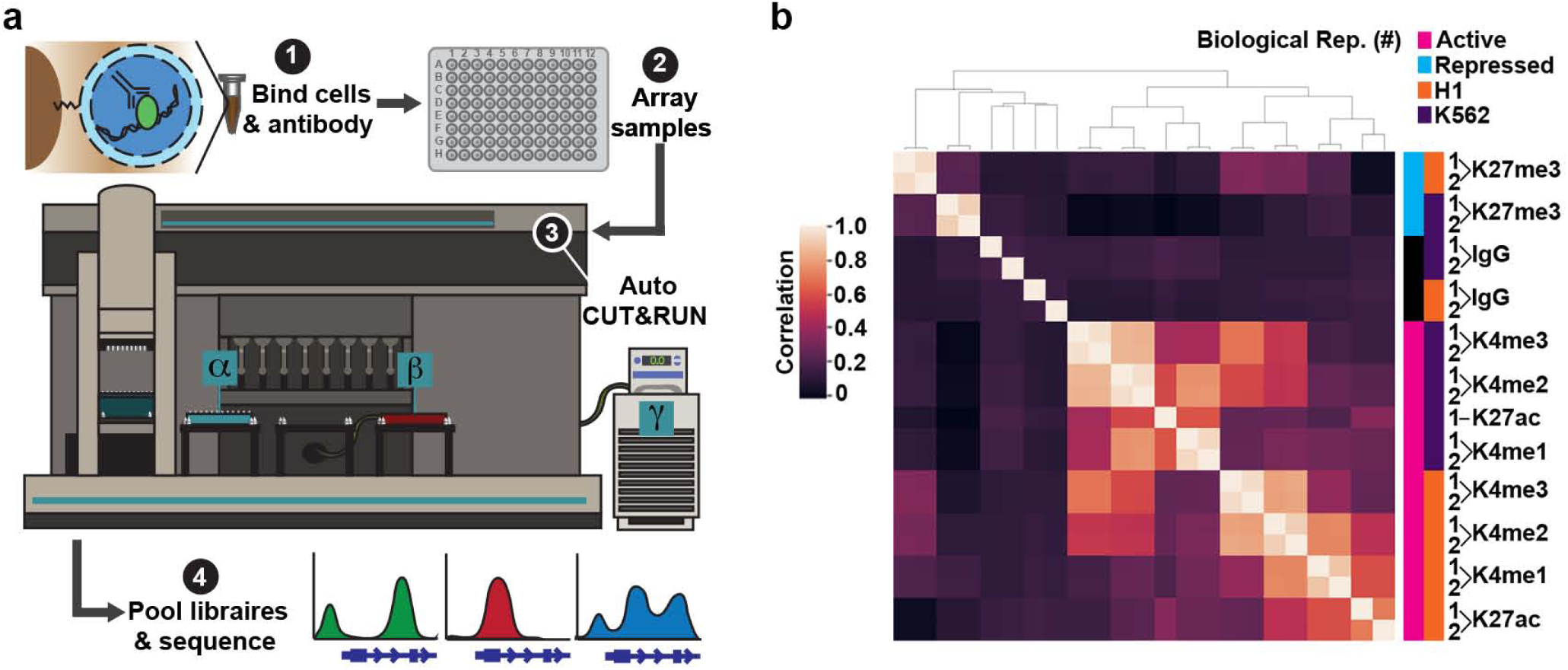
An automated platform for high-throughput *in situ* profiling of chromatin proteins. (**a**) AutoCUT&RUN workflow. (1) Cells or tissue are bound to Concanavalin A-coated beads, permeabilized with digitonin, and incubated with an antibody targeting a chromatin protein. (2) Samples are arrayed in a 96-well plate and (3) processed on a Biomek robot fitted with a 96-well magnetic plate for magnetic separation during washes (α), and an aluminum chiller block (β) routed to a circulating water bath (γ) for temperature control. (4) AutoCUT&RUN produces in 2 days up to 96 libraries that are ready to be pooled and sequenced. (**b**) Hierarchically clustered correlation matrix of AutoCUT&RUN profiles of histone-H3 modifications that mark active (pink) and repressed (blue) chromatin in H1 (orange) and K562 (purple) cells. Pearson correlations were calculated using the log_2_ transformed values of read counts split into 500 bp bins across the genome.

To test the consistency of AutoCUT&RUN, we simultaneously profiled two biological replicates of H1 hESCs and K562 cells using antibodies targeting four histone modifications that mark active chromatin states (H3K4me1, H3K4me2, H3K4me3, and H3K27ac) and one repressive modification (H3K27me3). Comparing the global distribution of reads for each histone mark, we found that samples highly correlate with their biological replicate, and cluster together in an unbiased hierarchical matrix (Fig. 1b). Additionally, the genome-wide profiles of the four active histone marks clustered together within a given cell type, and separated away from the repressive histone mark H3K27me3 (Fig. 1b). These profiles represent antibody-specific signals, as all five are poorly correlated with an IgG negative control. Together, these results indicate that AutoCUT&RUN chromatin profiling reproducibly captures the cell-type specific distributions of histone marks.

Histones are tightly associated with DNA in chromatin, so we also examined whether AutoCUT&RUN can be applied to mapping DNA-binding transcription factors that have lower residence times. We tested the performance of AutoCUT&RUN with two transcription factors, the histone locus-specific gene regulator NPAT, and the insulator protein CTCF^15,16^. AutoCUT&RUN profiles of both NPAT and CTCF are highly specific for their expected targets in both H1 and K562 cells (Supplementary Fig. 1b, c), and the signal sensitivity of CTCF in K562 cells was comparable to previous results^13^. Thus, AutoCUT&RUN is suitable for high-throughput, genome-wide profiling of diverse DNA binding proteins.

To maintain their developmental plasticity, hESCs have a generally open, hyper-acetylated chromatin landscape interspersed with repressed domains of “bivalent” chromatin, marked by overlapping H3K27me3 and H3K4 methylation^17-20^. AutoCUT&RUN recapitulates these features of hESCs; we observed that H1 cells have increased H3K27ac as compared to the lineage-restricted K562 cell line, whereas domains of the repressive histone mark H3K27me3 are rare in H1 cells, but prevalent in K562 cells (Fig. 2a). We also observed extensive overlap between H3K27me3 and H3K4me2 signals in H1 cells, but not K562 cells (Fig. 2a, b). Thus, Auto CUT&RUN profiles are consistent with the specialized chromatin features found in hESCs.

**Figure 2:**
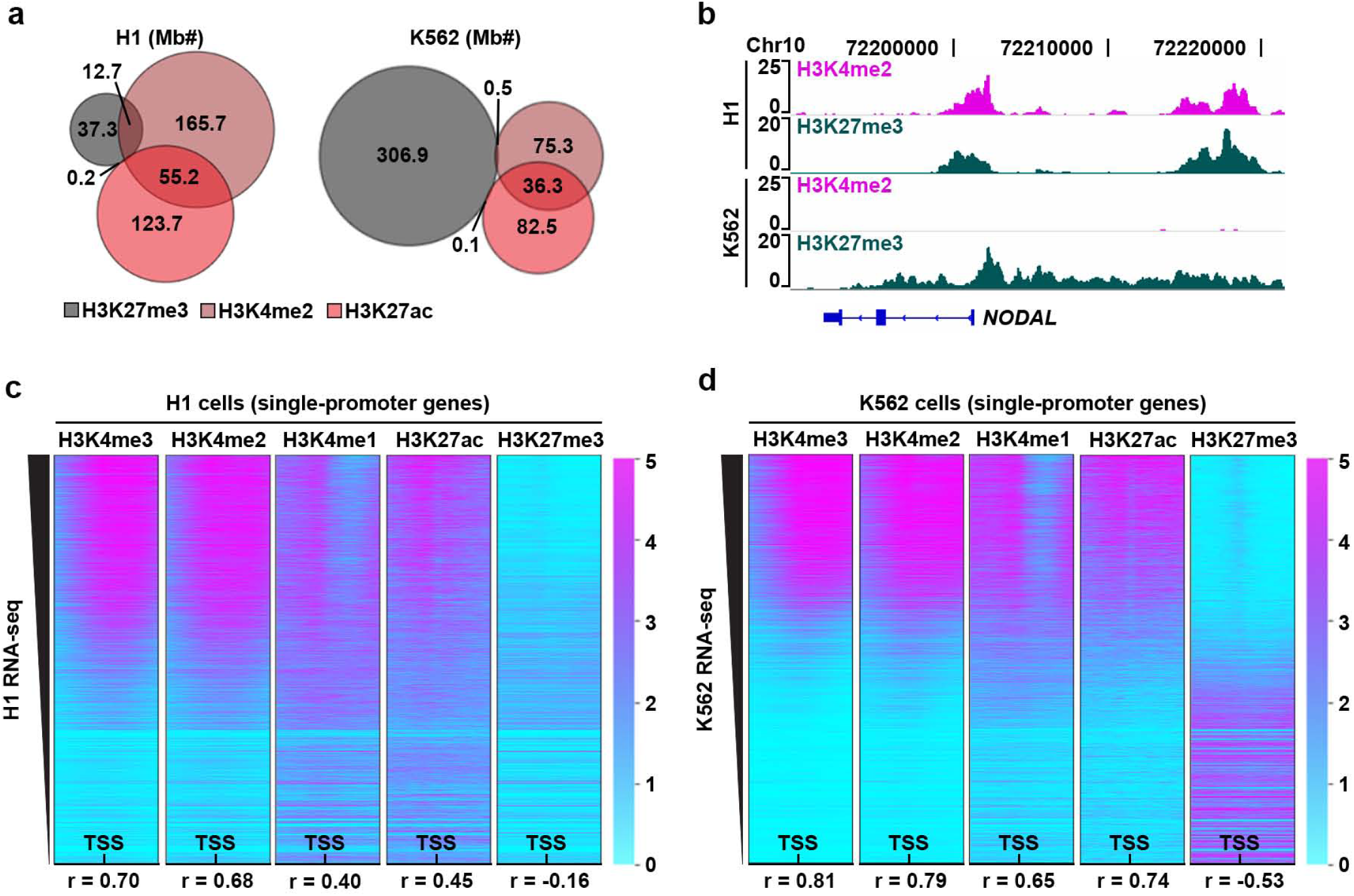
AutoCUT&RUN reproduces the expected chromatin landscape of H1 and K562 cells. (**a**) Scaled Venn diagrams showing the relative amount of the genome that falls within H3K27me3 (grey), H3K4me2 (brown), and H3K27ac (red) domains in H1 cells and K562 cells. Numbers indicate megabases (Mb). (**b**) Genome browser tracks showing the overlap of H3K4me2 and H3K27me3 in H1 cells, as well as the expansion of H3K27me3 domains and loss of overlap with H3K4me2 in K562 cells at a representative locus (*NODAL*). (**c**) Heat maps showing the distribution of AutoCUT&RUN profiles of histone modifications in H1 cells centered on the TSSs of genes with a single promoter, oriented left-to-right according to the 5’-to-3’ direction of transcription and rank-ordering according to RNA-seq values (FPKM). (**d**) Heat maps showing the distribution of AutoCUT&RUN histone modification profiles on transcriptionally active and repressed promoters in K562 cells. Pearson correlations (r-value) between AutoCUT&RUN profiles of individual histone marks around these TSSs and their corresponding RNA-seq values are indicated.

Post-translational modifications to the H3 histone tail closely correlate with transcriptional activity^21^. To determine whether our AutoCUT&RUN profiles of histone modifications are indicative of transcriptional activity, we examined the distribution of the five histone marks around the transcriptional start sites (TSSs) of genes, rank-ordered according to RNA-seq expression data (Fig. 2c, d)^22^. We find the active mark H3K4me3 is the most highly correlated with expression in both cell types (r = 0.70 and 0.81 for H1 and K562 respectively), followed by H3K4me2 and H3K27ac (Fig. 2c, d). The repressive histone mark H3K27me3 is anti-correlated with expression (r = −0.16 and −0.53 in H1 and K562 respectively) (Fig. 2c, d). We conclude AutoCUT&RUN for these marks recapitulates transcriptional activity, providing a strategy to identify cell-type specific gene regulatory programs.

### Modeling cell-type specific gene expression from AutoCUT&RUN profiles

To use AutoCUT&RUN data to compare cell-types and distinguish their gene regulatory programs we wanted to develop a continuous metric that incorporates both active and repressive chromatin marks. RNA-seq has been widely used to identify cell-type specific gene expression programs^22^, so we used RNA-seq data as a reference for training a weighted linear regression model that incorporates normalized H3K4me2, H3K27ac, and H3K27me3 read counts to assign promoters a relative activity score. We initially focused our analysis on genes with a single TSS that could be unambiguously assigned RNA-seq values. H3K4me2 was selected over H3K4me3 and H3K4me1 because H3K4me2 is uniquely applicable for modeling the activity of both proximal and distal cis-regulatory elements (see below). When applied to K562 cells, promoter chromatin scores correlate very well with RNA-seq values (r=0.83) (Fig. 3a), providing a comparable power for predicting gene expression as similar models that used up to 39 histone modifications mapped by ChIP-seq (r=0.81)^21^. In addition, our weighted model trained on K562 cells performs well when applied to H1 cells (Supplementary Fig. 2a, b), indicating that the linear model and data quality are sufficiently robust to assign promoter scores to diverse cell types.

**Figure 3:**
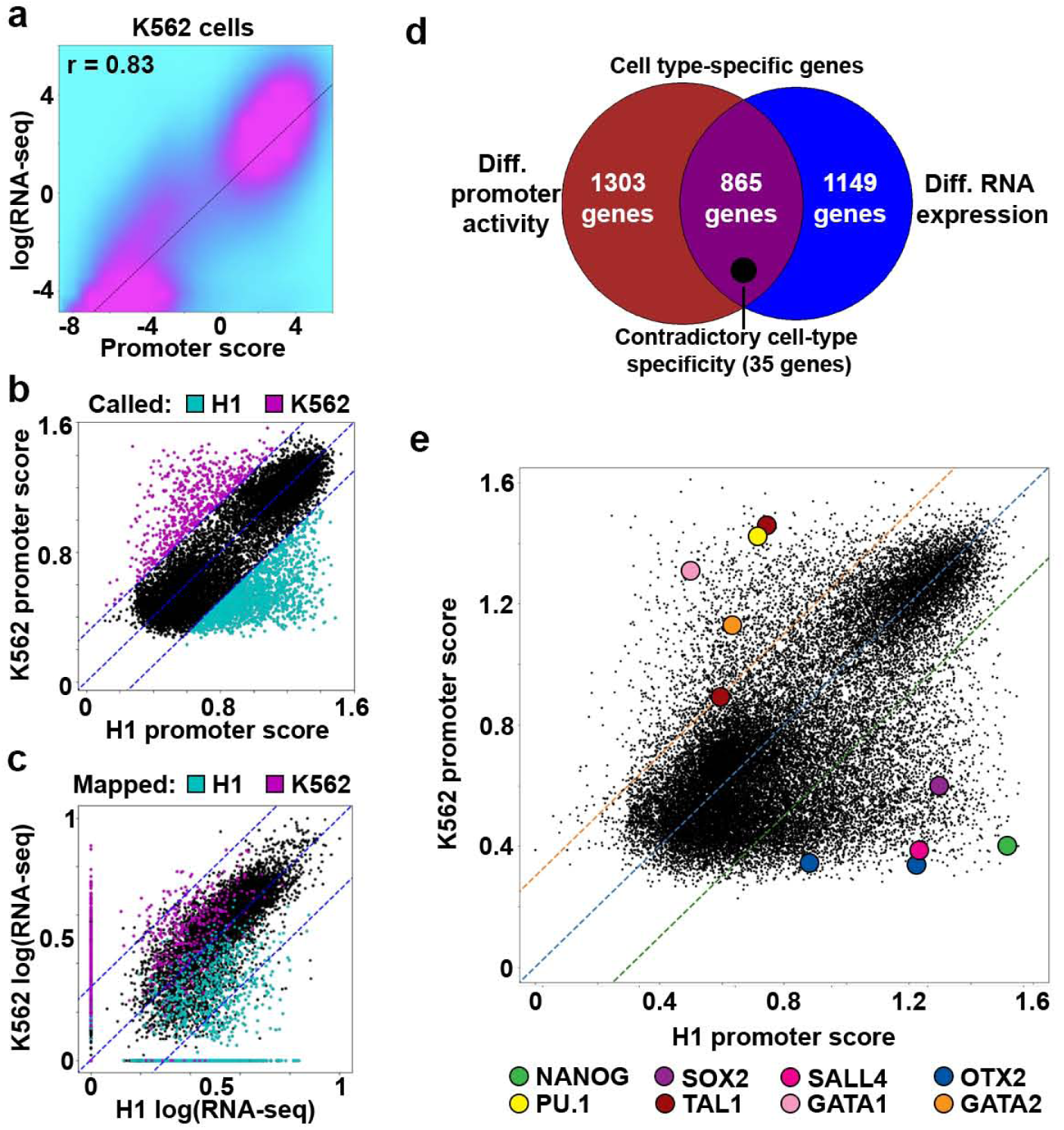
A linear regression model accurately predicts cell-type specific promoter activity. (**a**) Density scatterplot comparing RNA-seq values for single-promoter genes to K562 promoter scores predicted by the model trained on K562 data. (**b**) Scatterplot of promoter chromatin scores for single-promoter genes in H1 and K562 cells. Colored dots indicate that the promoter scores are ≥2-fold enriched in either H1 cells (cyan) or K562 cells (magenta). (**c**) Scatterplot of promoter scores that are ≥2-fold enriched in either H1 cells (cyan) or K562 cells (magenta) mapped onto their corresponding RNA-seq values. Blue dotted lines indicate the 2-fold difference cut-off. (**d**) Scaled Venn diagram showing the overlap between genes called as cell-type specific according to their promoter scores, or according to their RNA-expression values. Genes predicted to have contradictory cell-type specificities according to promoter activity modeling vs RNA-seq are indicated (scaled black circle). (**e**) Scatterplot comparing the H1 and K562 scores of all promoters separated by ≥2 kb. Master regulators of H1 and K562 cell identities are indicated as colored circles. Both OTX2 and TAL1 have two promoters that can be distinguished.

Next, we examined if AutoCUT&RUN accurately identifies promoters with cell-type specific activity. By calling promoter scores that were enriched more than two-fold in either H1 or K562 cells we identified 2,168 cell-type specific genes and approximately 40% of these genes (865) were also differentially-enriched between H1 and K562 cells according to RNA-seq (Fig. 3b-d). However, promoter activity modeling did not capture transcriptional differences for 1149 genes (Fig 3d, Supplementary Fig. 2c, d), implying that these genes are differentially expressed without changes in the chromatin features included in our model. This differential sensitivity between methods suggests the three histone marks included in our chromatin model may more accurately predict the cell type specific expression of certain classes of genes than others. Indeed, we find the 865 cell-type specific genes identified by both promoter activity modeling and RNA-seq are highly enriched for developmental regulators, whereas the genes called by either promoter scores or RNA-seq alone are not nearly as enriched for developmental GO terms (Fig. 3d, Supplementary Fig. 2e-g, Supplementary table 1). In addition, only 35 genes display contradictory cell-type specificities according to promoter chromatin scores and RNA-seq (Fig. 3d). This demonstrates AutoCUT&RUN profiling of these widely studied modification to the H3 histone tail can be applied to accurately identify cell-type specific developmental regulators.

To determine whether AutoCUT&RUN data recapitulates the expression of cell-type-specific transcription factors, we expanded our analysis to include all promoters. We find that components of the hESC pluripotency network (*NANOG, SOX2, SALL4*, and *OTX2*) have higher promoter chromatin scores in H1 cells, while regulators of hematopoetic progenitor cell fate (*PU.1, TAL1, GATA1*, and *GATA2*) are enriched in K562 cells (Fig. 3e, Supplementary Table 1)^23,24^. This method also identifies activities of alternative promoters (e.g. at the *OTX2* and *TAL1* genes), providing an indication of the specific gene isoforms that are expressed in a given cell-type (Fig. 3e). We conclude that AutoCUT&RUN allows identification of master regulators of cellular identity, providing a powerful tool to characterize cell-types in a high-throughput format.

### Profiling tumors by AutoCUT&RUN

Traditional methods to profile protein-DNA interactions (e.g. ChIP-seq) are generally unable to handle clinically-relevant samples, which often contain small amounts of starting material and have been flash-frozen. To test whether AutoCUT&RUN is suitable for profiling frozen tumor specimens, we obtained two diffuse midline glioma (DMG) patient-derived cell lines (VUMC-10 and SU-DIPG-XIII) that were autopsied from similar regions of the brainstem, but differ in their oncogenic backgrounds^25^. Both of these DMG cell lines readily form xenografts in murine models, and we applied AutoCUT&RUN to profile histone modifications in VUMC-10 and SU-DIPG-XIII xenografts that were seeded in the brains of mice, and then resected upon tumor formation and frozen under typical clinical conditions (Fig. 4a). For comparison, on the same AutoCUT&RUN plate we profiled the parental DMG cell lines grown in culture (Fig. 4a). Again, we found that replicates were highly concordant, so we combined them for further analysis. Importantly, cell culture samples were highly correlated with the same mark profiled in the corresponding frozen xenografts, and AutoCUT&RUN on xenograft tissues and cell culture samples produced similar data quality (Fig. 4b, Supplementary Fig. 3). Thus, AutoCUT&RUN reliably generates genome-wide chromatin profiles from frozen tissue samples.

**Figure 4:**
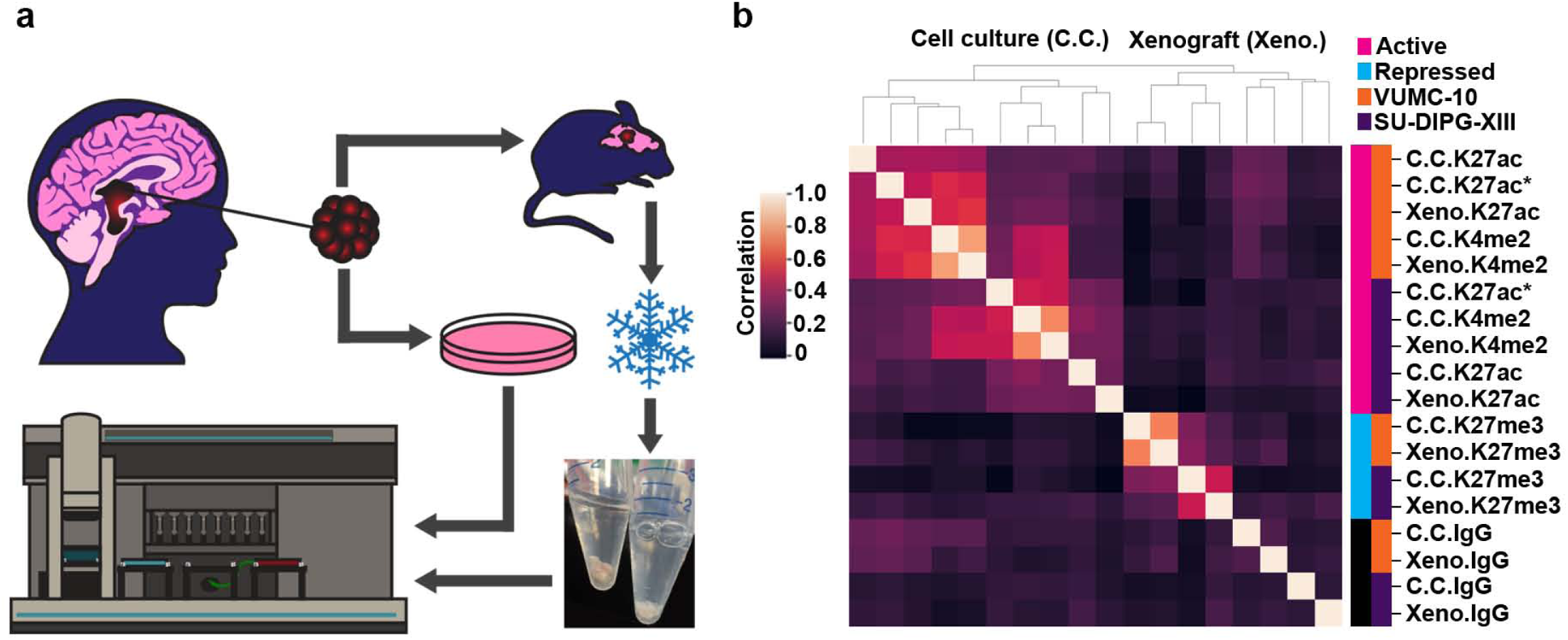
AutoCUT&RUN is suitable for profiling the chromatin landscape of frozen tumor samples. (**a**) DMG experimental set up. Two DMG cell lines derived from a similar region of the brainstem were grown as xenografts in the brains of immuno-compromised mice and upon forming tumors were resected and frozen. Xenografts were thawed and process by AutoCUT&RUN in parallel with control DMG samples harvested directly from cell culture. (**b**) Hierarchically clustered correlation matrix of AutoCUT&RUN profiles of histone-H3 modifications that mark active (pink) and repressed (blue) chromatin in VUMC-10 (orange) and SU-DIPG-XIII (purple) cells grown in cell culture (C.C.) or as xenografts (Xeno.). As a quality control H3K27ac was also profiled manually in these cell lines using a different antibody (*). Pearson correlations were calculated using the log_2_ transformed values of paired-end read counts split into 500 bp bins across the genome.

Stratification of patient malignancies is becoming increasingly dependent on molecular diagnostic methods that distinguish tumor subtypes derived from the same tissues. Our VUMC-10 and SU-DIPG-XIII samples provide an excellent opportunity to explore the potential of using AutoCUT&RUN to classify tumor specimens according to their subtype-specific regulatory elements. By applying our model to these samples, we identified 5,006 promoters that show differential activity between VUMC-10 and SU-DIPG-XIII cells (Fig. 5a, Supplementary Table 1). Consistent with the glial origins of these tumors, both the VUMC-10 and SU-DIPG-XIII specific promoters are significantly enriched for genes involved in nervous system development (Supplementary Fig. 4a, b). Genes involved in cell-signaling are also overrepresented in SU-DIPG-XIII cells (Supplementary Fig. 4b); for example, the promoters of the PDGFR gene as well as its ligand PDGF are highly active in SU-DIPG-XIII cells (Fig. 5a). This is consistent with the observation that DMGs frequently contain activating mutations in PDGFR-α that promote tumor growth^5^. In addition, one promoter of the SMAD3 gene, a component of the TGF-β signaling pathway^26^, is specifically active in SU-DIPG-XIII cells, whereas two different SMAD3 promoters are active in VUMC-10 cells (Fig. 5a, Supplementary Fig. 3). In comparison, our model indicates that only 388 promoters differ between VUMC-10 xenografts and cultured cells, and 1,619 promoters differ between SU-DIPG-XIII samples (Fig. 5b, Supplementary Fig. 4c). In addition, comparing promoter chromatin scores in an unbiased correlation matrix also indicates DMG xenografts are far more similar to their corresponding cell culture samples than they are to other DMG subtypes or to H1 or K562 cells (Fig. 5c). This suggests that AutoCUT&RUN can be applied to identify promoters that display tumor subtype-specific activity, providing a reliable method to assign cellular identities to frozen tumor samples, as well as an indication of the signaling pathways that may be driving tumor growth and potential susceptibility to therapeutic agents.

**Figure 5:**
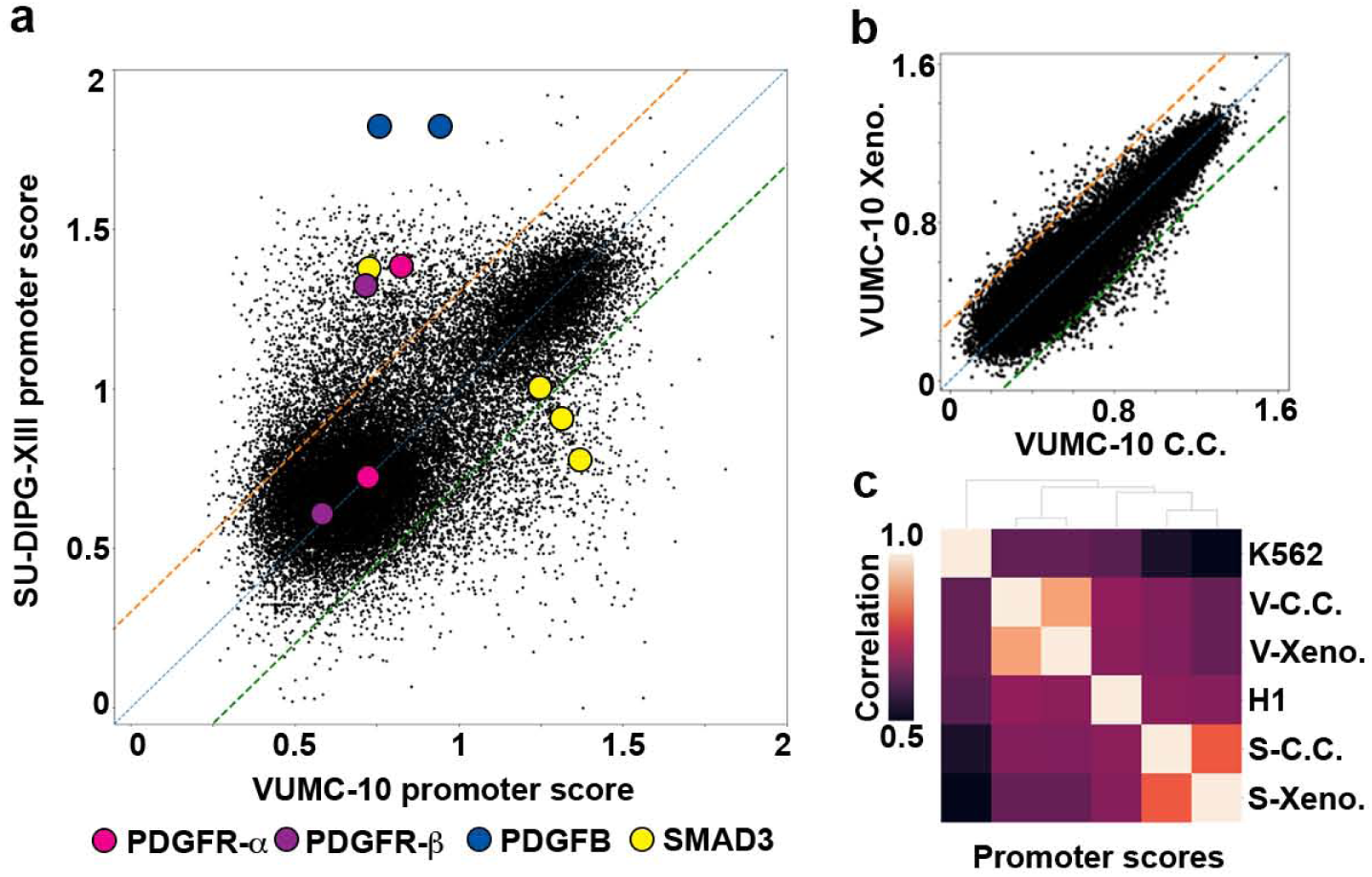
Promoter activity modeling distinguishes gene activities in DMG samples. (**a**) Scatterplot comparing the promoter scores of VUMC-10 and SU-DIPG-XIII cell culture samples. Locations of the promoters of several cell-signaling components implicated in tumor growth are indicated as colored circles. (**b**) Scatterplot comparing the promoter scores of VUMC-10 cell culture (C.C.) and xenograft (Xeno.) samples. Only 388 promoters have a ≥2-fold difference in activity modeling scores between these samples. (**c**) Hierarchically clustered matrix of Spearman correlations of promoter chromatin scores between VUMC-10 (V) and SU-DIPG-XIII (S) cells grown in cell culture (C.C.) or as xenografts (Xeno.), as well as H1 and K562 cells.

### High-throughput mapping of cell-type specific enhancers

The cell-type specific activities of gene promoters are often established by incorporating signals from distal cis-regulatory elements, such as enhancers^1,2^. Similar to promoters, enhancers also display H3K4me2^27^, and active enhancers are typically marked by H3K27ac, whereas repressed enhancers are marked by H3K27me3^20,28,29^. Therefore, we reasoned the AutoCUT&RUN profiles we used to model promoter activity should also allow identification of cell-type specific enhancers. To investigate this possibility, we first compared our H1 data to available chromatin accessibility maps generated by ATAC-seq, which are enriched for both active promoters and enhancers^30,31^. Of the marks we profiled, we find H3K4me2 peaks show the highest overlap with ATAC-seq (Fig 6a, Supplementary Fig. 5a), and identify 36,725/52,270 ATAC-seq peaks (∼70%). Interestingly, H3K4me2 defines an additional 71,397 peaks that were not called by ATAC-seq (Fig. 6a, Supplementary Fig. 5a). Many of these H3K4me2-specific peaks show a low, but detectable ATAC-seq signal (Supplementary Fig. 5b), indicating they may correspond to repressed promoters and enhancers. Consistent with this interpretation, on average H3K4me2+/ATAC-TSSs have higher H3K27me3 signals than H3K4me2+/ATAC+ TSSs (Supplementary Fig. 5c). H3K4me2+/ATAC+ peaks that overlap with annotated TSSs are enriched for H3K4me3, while those peaks that do not overlap TSSs are enriched for H3K4me1 (Fig. 6b&c, Supplementary Fig. 5d), suggesting that many of these distal peaks are enhancers^20,32^. Thus, mapping sites of H3K4me2 by AutoCUT&RUN provides a sensitive method for defining the repertoire of active cis-regulatory elements that control gene expression programs.

**Figure 6:**
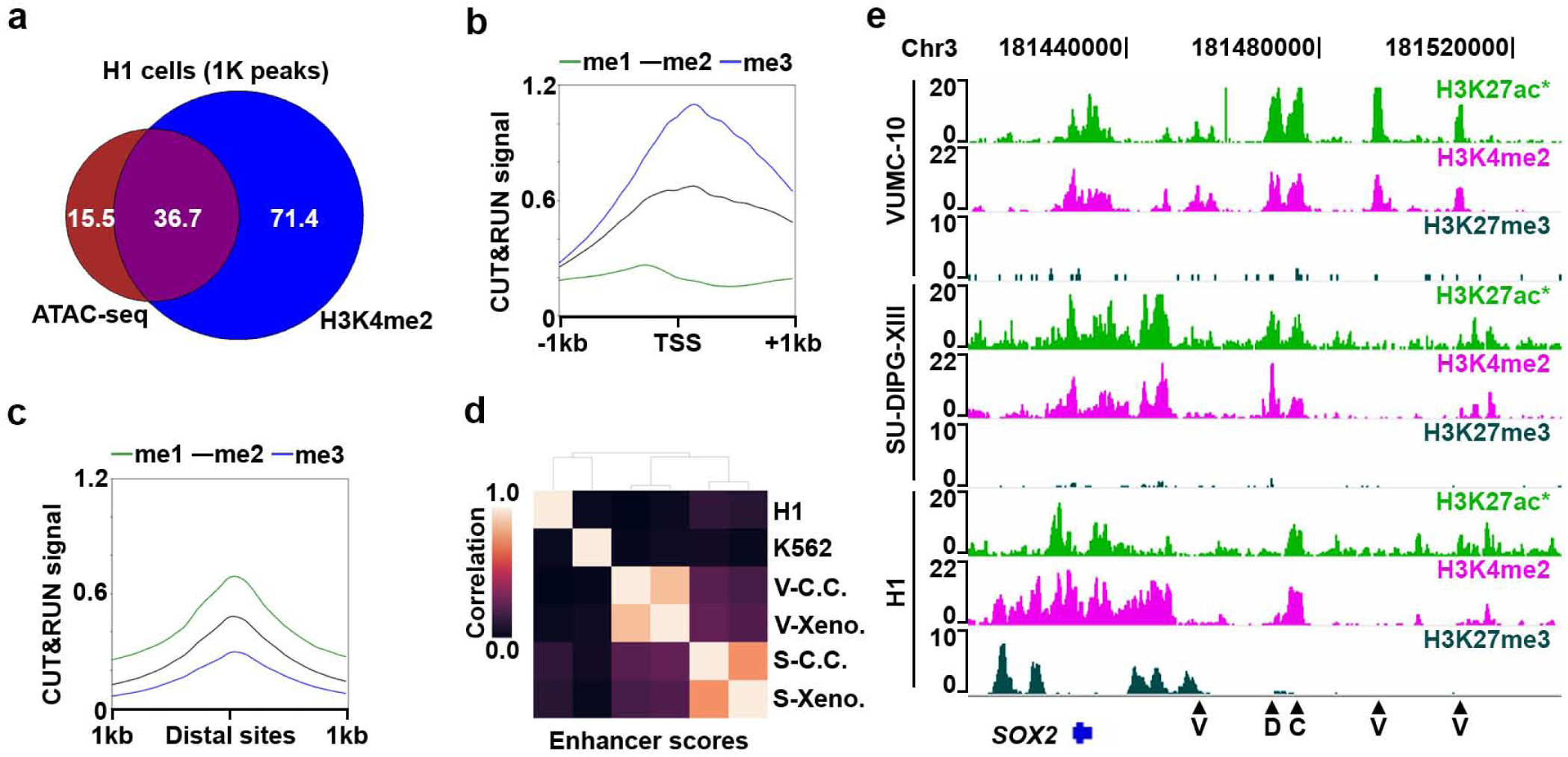
AutoCUT&RUN identifies cell-type specific enhancer elements. (**a**) Scaled Venn diagram showing the overlap of accessible chromatin sites (ATAC-seq peaks) and peaks called on H3K4me2 AutoCUT&RUN profiles in H1 cells. Numbers are provided as 1 thousand peak units. (**b**) Mean enrichment of H3K4me1 (green) H3K4me2 (black) and H3K4me3 (blue) over all H3K4me2+/ATAC+ TSSs. (**c**) Mean enrichment of H3K4me1 (green) H3K4me2 (black) and H3K4me3 (blue) over all H3K4me2+/ATAC+ distal sites. (**d**) Hierarchically clustered matrix of Spearman correlations of enhancer chromatin scores in VUMC-10 (V) and SU-DIPG-XIII (S) cells grown in cell culture (C.C.) or as xenografts (Xeno.), as well as H1 and K562 cells. (**e**) Genome browser tracks showing the location of putative enhancer elements (arrow heads) that are specific to VUMC-10 cells (V), both DMG cell lines (D), or common to DMG cells and H1 cells (C) at a representative locus (*SOX2*). In VUMC-10 and SU-DIPG-XIII cells the H3K27ac track that is shown was also profiled manually as a quality control (*).

Finally, we examined whether AutoCUT&RUN can be used to identify cell-type-specific enhancers. To expand the number of putative enhancer sites, we compiled a list of non-TSS peaks called on H3K4me2 profiles from all six cell lines and xenograft samples. Using our linear regression model, we then assigned these elements chromatin scores and examined their correlations between different cell types. We find that these chromatin scores of DMG cell culture samples and xenografts are highly correlated (r=0.75 and 0.87 for SU-DIPG-XIII and VUMC-10 samples respectively) (Fig. 6d). In contrast, the chromatin scores of SU-DIPG-XIII cells show a weak positive correlation with VUMC-10 cells (e.g. r=0.19), indicating tumor subtype-specific differences. For example, the different enhancers near the *SOX2* pluripotency gene are active in SU-DIPG-XIII and VUMC-10 cells (Fig. 6e), indicating that SU-DIPG-XIII cells resemble a more primitive neural stem cell-type than VUMC-10 cells, as has been previously suggested^33^. Thus, AutoCUT&RUN provides a stringent method for stratifying cell-types and tissue samples.

## Discussion

We adapted the CUT&RUN technique to an automated platform by developing direct ligation of chromatin fragments for Illumina library preparation, and implementing magnetic separation for the wash steps and library purification. AutoCUT&RUN generates 96 genome-wide profiles of antibody-targeted chromatin proteins in just two days, dramatically increasing the throughput and potential scale of studies to interrogate the chromatin landscape at a fraction of the cost of comparable lower-throughput methods. We show that profiling just three histone modifications (H3K27ac, H3K27me3 and H3K4me2) is sufficient to determine the cell-type specific activities of developmentally regulated promoters and enhancers, providing a powerful quantitative metric to compare the epigenetic regulation of different cell-types. This summary metric of chromatin features could be used to assess new cell types and tissue samples and to place them within a reference map of both healthy and diseased cell types. The automated workflow reduces technical variability between experiments, generating consistent profiles from biological replicates and from different sample types.

To continue optimizing AutoCUT&RUN one could envision hardware modifications and computational development. By screening various antibody collections, the repertoire of nuclear proteins that can be efficiently profiled using AutoCUT&RUN would expand dramatically. In addition, the current AutoCUT&RUN protocol is optimized for a popular liquid handling robot, but a custom robot incorporating a reversibly magnetic thermocycler block would allow the CUT&RUN reaction and library preparation to be carried out in place, streamlining the protocol even further. Last, metrics distinguishing cell types could be improved by incorporating additional aspects of the data, such as using a combination of both enhancer and promoter activities.

The excellent reproducibility of profiling frozen tissue samples by AutoCUT&RUN has the potential to transform the field of epigenomic medicine^10^. Compared to other genomics approaches that are currently used for patient diagnosis, AutoCUT&RUN has the unique capacity to profile chromatin proteins within diseased cells. For example, cancers caused by fusions between chromatin proteins could be profiled by AutoCUT&RUN to provide a molecular diagnosis based on their chromatin landscapes, while simultaneously mapping the loci that are disrupted by the mutant fusion protein. This could provide a powerful tool for patient stratification, as well as a direct read-out of whether chromatin-modulating therapies such as histone deacetylase or histone methyltransferase inhibitors are having their intended effects.

## Online Methods

### AutoCUT&RUN

A detailed AutoCUT&RUN protocol is provided (Appendix 1). Briefly, cells or tissue samples are bound to Concanavalin A coated magnetic beads (Bangs Laboratories, ca. no. BP531), permeabilized with digitonin, and bound with a protein specific antibody as previously described^12^. Samples are then arrayed in a 96-well plate and processed on a Beckman Biomek FX liquid handling robot equipped with a 96S Super Magnet Plate (Alpaqua SKU A001322) for magnetic separation of samples during wash steps, and an Aluminum Heat Block Insert for PCR Plates (V&P Scientific, Inc. VP741I6A) routed to a cooling unit to perform the MNase digestion reaction at 0-4 °C after the addition of 2 mM CaCl_2_. MNase digestion reactions are then stopped after 9 min by adding EGTA, which allows Mg^2+^ addition for subsequent enzymatic reactions. This step is critical for automation because it circumvents the need for DNA purification prior to library preparation. Chromatin fragments released into the supernatant during digestion are then used as the substrate for end-repair and ligation with barcoded Y-adapters. Prior to ligation, the A-tailing step is performed at 58°C to preserve sub-nucleosomal fragments in the library^34,35^. End-repair and adapter ligation reactions were performed on a separate thermocycler. Chromatin proteins are then digested with Proteinase K, and adapter ligated DNA fragments are purified on the Biomek FX using two rounds of pre-PCR Ampure bead cleanups with size-selection. PCR enrichment reactions sre performed on a thermocycler using the KAPA PCR kit (KAPA Cat#KK2502). Two rounds post-PCR Ampure bead cleanups with size-selection are performed on the Biomek FX to remove unwanted proteins and self-ligated adapters.

The size distributions of AutoCUT&RUN libraries were analyzed on an Agilent 4200 TapeStation, and library yield was quantified by Qubit Fluorometer (Life Technologies). Up to 24 barcoded AutoCUT&RUN libraries were pooled per lane at equimolar concentration for paired-end 25−25 bp sequencing on a 2-lane flow cell on the Illumina HiSeq 2500 platform at the Fred Hutchinson Cancer Research Center Genomics Shared Resource.

### Antibodies

We used Rabbit anti-CTCF (1:100, Millipore Cat#07-729), Rabbit anti-NPAT (1:100, Termo Fisher Cat#PA5-66839), Rabbit anti-H3K4me1 (1:100, Abcam Cat#ab8895), Rabbit anti-H3K4me2 (1:100, Millipore Cat#07-030), Rabbit anti-H3K4me3 (1:100, Active Motif Cat#39159), Rabbit anti-H3K27me3 (1:100, Cell Signaling Tech Cat#9733S). Since pAMNase does not bind efficiently to many mouse antibodies, we used a rabbit anti-Mouse IgG (1:100, Abcam, Cat#ab46540) as an adapter. H3K27ac was profiled by AutoCUT&RUN in H1 and K562 cells and manually in VUMC-10 and SU-DIPG-XIII cell lines using Rabbit anti-H3K27ac (1:50, Millipore Cat#MABE647). H3K27ac was profiled by AutoCUT&RUN in VUMC-10 and SU-DIPG-XIII cell lines and xenografts using Rabbit anti-H3K27ac (1:100, Abcam Cat#ab45173).

### Cell Culture

Human K562 cells were purchased from ATCC (Manassas, VA, Catalog #CCL-243) and cultured according to supplier’s protocol. H1 hESCs were obtained from WiCell (Cat#WA01-lot#WB35186), and cultured in Matrigel™ (Corning) coated plates in mTeSR™1 Basal Media (STEMCELL Technologies cat# 85851) containing mTeSR™1 Supplement (STEMCELL Technologies cat# 85852). Pediatric DMG cell lines VUMC-DIPG-10 (Esther Hulleman, VU University Medical Center, Amsterdam, Netherlands) and SU-DIPG-XIII (Michelle Monje, Stanford University, CA) were obtained with material transfer agreements from the associated institutions. Cells were maintained in NeuroCult NS-A Basal Medium with NS-A Proliferation Supplement (STEMCELL Technologies, cat# 05751), 100 U/mL of penicillin/streptomycin, 20ng/mL epidermal growth factor (PeproTech, cat# AF-100-15), and 20ng/mL fibroblast growth factor (PeproTech, cat# 100-18B).

### Patient-derived Xenografts

All mouse studies were conducted in accordance with Institute of Animal Care and Use Committee-approved protocols. NSG mice were bred in house and aged to 2-3 months prior to tumor initiation. Intracranial xenografts were established by stereotactic injection of 100,000 cells suspended in 3uL at a position of 2 mm lateral and 1mm posterior to lambda. Symptomatic mice were euthanized and their tumors resected for analysis.

### Annotation and Data Analysis

We aligned paired-end reads using Bowtie2 version 2.2.5 with options: --local --very-sensitive-local --no-unal --no-mixed --no-discordant --phred33 -I 10 -X 700. For mapping spike-in fragments, we also used the --no-overlap --no-dovetail options to avoid cross-mapping of the experimental genome to that of the spike-in DNA^36^. Files were processed using bedtools and UCSC bedGraphToBigWig programs^37,38^.

To examine correlations between the genome-wide distributions of various samples, we generated bins of 500 bp spanning the genome, creating an array with approximately 6 million entries. Reads in each bin were counted and the log_2_ transformed values of these bin counts were used to determine a Pearson correlation score between different experiments. Hierarchal clustering was then performed on a matrix of the Pearson scores.

To examine the distribution of histone mark profiles around promoters, a reference list of genes for build hg19 were downloaded from the UCSC table browser (https://genome.ucsc.edu/cgi-bin/hgTables) and oriented according to the directionality of gene transcription for further analysis. Genes with TSSs within 1 kb of each other were removed, as were genes mapping to the mitochondrial genome, creating a list of 32,042 TSSs. RNA-sequencing data was obtained from the ENCODE project for H1 and K562 cells (ENCSR537BCG and <u>ENCSR000AEL</u>). RNA reads were counted using featureCounts (http://bioinf.wehi.edu.au/featureCounts/), and converted to Fragments Per Kilobase per Million mapped reads (FPKM) and assigned to the corresponding TSS as a gene expression value. ATAC-sequencing data for H1 cells was obtained from gene omnibus expression (GEO) (<u>GSE85330</u>) and mapped to hg19 using bowtie2. Mitochondrial DNA accounted for ∼50% of the reads and were removed in this study.

All heat maps where generated using DeepTools^39^. All of the data was analyzed using either bash or python. The following packages were used in python: Matplotlib, NumPy, Pandas, Scipy, and Seaborn.

### Training the linear regression model

To ensure the accuracy of fitting histone modification data at promoters to RNA-seq values, genes with more than one promoter were removed from the previously generated TSS list. The genes RPPH1 and RMRP were expressed at extremely high levels in H1 cells, and so were considered to be outliers and were removed to avoid skewing the regression, leaving a list of n=12,805 genes.

To assign a relative CUT&RUN signal to promoters for each histone mark, denoted by C, base pair read counts +/– 1 kb of the TSS were normalized by both sequencing depth over the promoters being scored and the total number of promoters examined. The prior normalization is to account for both sequencing depth and sensitivity differences among antibodies, and the latter normalization is included so that the model can be applied to different numbers of *cis*-regulatory elements without changing the relative weight of each element. FPKM values were used for RNA-seq.

The linear regression model was trained to fit profiles of histone marks to RNA expression values as previously described^21^. Briefly, we used a linear combination of histone data fitted to the RNA-seq expression values: *y* = *C*_1_*x*_1_ + … + *C*_*n*_*x*_*n*_, where *C*_*i*_ is the weight for each histone modification and *x*_*i*_ is denoted by *x*_*i*_ = In (*C*_*i*_ + *☐*_*i*_), where C is the normalized base-pair counts described above and *α* is a pseudo-count to accommodate genes with no expression. The RNA-seq values were similarly transformed as *y*_*i*_ = In (*FPKM*_*i*_ + *☐*_*y,i*_). Logarithmic transformations were used to linearize the data. A minimization step was then performed to calculate pseudo-counts and weights for each histone modification that would maximize a regression line between CUT&RUN data and RNA-seq.

We expected that the histone marks H3K27ac, H3K27me3, and H3K4me2 would provide the least redundant information. The optimized three histone mark model for K562 cells is described by:

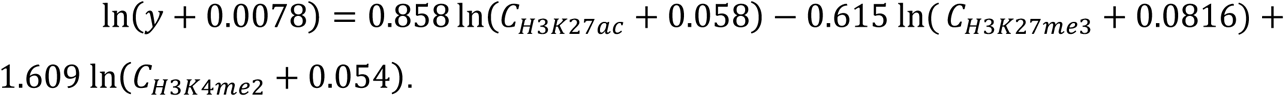

This equation was used to generate all chromatin activity scores.

### Calling chromatin domain for overlap analysis

To compare the global chromatin landscape of H1 and K562 cells, chromatin domains were called using a custom script that enriched for regions relative to an IgG CUT&RUN control. Enriched regions among marks were compared and overlaps were identified by using bedtools intersect. Overlapping regions were quantified by the number of common base pairs, and these were used to generate the Venn diagrams.

### Venn Diagrams

All Venn Diagrams were generated using the BaRC webtool, publicly available from the Whitehead Institute (http://barc.wi.mit.edu/tools/venn/).

### Calculating cell-type specific promoter activity scores

Raw promoter chromatin scores generally fall within a range from −10 to 10, where a smaller number is indicative of less transcriptional activity. To account for outliers in the data when comparing different cell types, promoter scores within 2 standard deviations were z-normalized. Negative and zero values complicate calculating fold change, so the data were shifted in the x and y directions by the most negative values. The fold difference between promoter scores for various cell types was calculated by dividing the inverse log_10_-normalized promoter scores against each other. A conservative 2-fold cutoff was used to determine cell-type specific promoters in each case (Fig 3b-e, 5a-b). Each list of genes was classified by gene ontology (http://geneontology.org/) to identify statistically enriched biological processes.

To examine the relative similarities between cell-types based on their promoter activities, scores for all promoters ≥1 kb apart were used to generate an array, and Spearman correlations were calculated for each pair-wise combination of the samples. Hierarchal clustering of the Spearman correlation values was used to visualize the relative similarities between cell-types.

### Peak calling on AutoCUT&RUN and ATAC-seq data

Biological replicates profiled by AutoCUT&RUN were highly correlated (Fig. 1b), so replicates were joined prior to calling peaks. The tool MACS2 was used to call peaks^40^. The following command was used: “macs2 callpeak -t file -f BEDPE -n name -q 0.01 -- keep-dup all -g 3.137e9”. An FDR cutoff of 0.01 was used

### Calculating cell-type specific enhancer activity scores

To assemble a list of distal cis-regulatory elements in the human genome, we used MACS2 to call peaks on H3K4me2 profiles from each of our samples using the same flags described in the ‘Peak calling on AutoCUT&RUN and ATAC-seq’ methods section. To distinguish between TSSs and putative enhancers, peaks <2.5 kb away from an annotated TSS were removed, and windows +/– 1kb around these putative enhancers were assigned chromatin activity scores using the algorithm trained to predict promoter activity. Correlation matrices comparing the enhancer scores between samples were generated in the same manner as the correlation matrix comparing promoter scores between samples.

## Acknowledgments

We thank the Fred Hutchinson Genomics Shared Resource Facility for technical support, particularly Jenni Risler and Dr. Jeff Delrow for help programming and operating the Biomek FX robot. We thank Terri Bryson for help with cell culture, Christine Codomo for preparing libraries for sequencing, and Jorja Henikoff for preparing the sequencing data for analysis. In addition, we thank Dr. Giancarlo Bonora and Dr. William Noble for helpful discussions related to data analysis. Last, we thank all members of the Henikoff lab as well as Dr. Antoine Molaro for helpful discussions over the course of this project. This work was supported by a grant from the Chan-Zuckerberg Initiative (S.H.), by the Howard Hughes Medical Institute (S.H.) and by a Damon Runyon-Sohn Foundation Fellowship (J.F.S.).

## Author Contributions

D.H.J. and S.H. optimized the AutoCUT&RUN protocol. D.H.J., J.F.S., K.A. and S.H. designed experiments. D.H.J. performed experiments with the help of C.H.M. and J.M.O. who obtained the DMG cell lines and prepared the patient derived xenograft samples. D.H.J., S.J.W. and M.P.M. developed algorithms and analyzed the data. D.H.J., K.A. and S.H. wrote the manuscript.

## Figure Legends

**Appendix 1: AutoCUT&RUN Protocol**

### MATERIALS

#### REAGENTS

- Cell suspension. We have used human K562 and H1 cells (see H1 suspension protocol), as well as several human brain tumor lines, Drosophila S2 cells and dissected Drosophila tissues such as brains and imaginal disks, and spheroplasted yeast.
- CAUTION: The cell lines used in your research should be regularly checked to ensure they are authentic and are not infected with mycoplasma.
- Concanavalin A-coated magnetic beads (Bangs Laboratories, ca. no. BP531)
- Antibody to an epitope of interest. For example, rabbit anti-CTCF polyclonal antibody (Millipore cat. no. 07-729) for mapping 1D and 3D interactions by CUT&RUN
- Positive control antibody to an abundant epitope, *e.g.* anti-H3K27me3 rabbit monoclonal antibody (Cell Signaling Technology, cat. no. 9733)
- Negative control antibody to an absent epitope, *e.g.* guinea pig anti-rabbit antibody
- 5% (wt/vol) Digitonin (EMD Millipore, cat. no. 300410)
- Protein A–Micrococcal Nuclease (pA-MNase) fusion protein (provided in 50% (vol/vol) glycerol by the authors upon request). Store at −20 °C.
- Spike-in DNA (e.g., from *Saccharomyces cerevisiae* micrococcal nuclease-treated chromatin, provided by authors upon request)
- Distilled, deionized or RNAse-free H_2_O (dH_2_O e.g., Promega, cat. no. P1197)
- 1 M Manganese Chloride (MnCl_2_; Sigma-Aldrich, cat. no. 203734)
- 1 M Calcium Chloride (CaCl_2_; Fisher, cat. no. BP510)
- 1 M Potassium Chloride (KCl; Sigma-Aldrich, cat. no. P3911)
- 1M 4-(2-hydroxyethyl)-1-piperazineethanesulfonic acid) pH 7.5 (HEPES (Na^+^); Sigma-Aldrich, cat. no. H3375)
- 1 M 4-(2-hydroxyethyl)-1-piperazineethanesulfonic acid) pH 7.9 (HEPES (K^+^); Sigma-Aldrich, cat. no. H3375)
- 5 M Sodium chloride (NaCl; Sigma-Aldrich, cat. no. S5150-1L)
- 0.5 M Ethylenediaminetetraacetic acid (EDTA; Research Organics, cat. no. 3002E)
- 0.2 M Ethylene glycol-bis(β-aminoethyl ether)-N,N,N’,N’-tetraacetic acid (EGTA; Sigma-Aldrich, cat. no. E3889)
- 2 M Spermidine (Sigma-Aldrich, cat. no. S2501)
- Roche Complete Protease Inhibitor EDTA-Free tablets (Sigma-Aldrich, cat. no. 5056489001)
- RNase A, DNase and protease-free (10 mg/mL; Thermo Fisher Scientific, cat. no. EN0531)
- 10X T4 DNA ligase buffer (NEB #B02025)
- 10mM dNTPs (KAPA #KK1017)
- 10mM ATP (NEB #P0756S)
- 40% PEG 4000 (Sigma-Aldrich, cat. no. 81242)
- 40% PEG 8000 (Sigma-Aldrich, cat. no. 202452)
- T4 PNK (NEB #M0201S)
- T4 DNA polymerse (Invitrogen #18005025)
- Taq DNA polymerase (Thermo #EP0401)
- DNA Seq Adapters. The protocol is optimized for use with TruSeq Y adapters with a free 3’T overhang (*:thiol bond, P-:phosphate group).
- TruSeq Universal Adapter: AATGATACGGCGACCACCGAGATCTACACTCTTTCCCTACACGACGCTCTTCCGATC*T
- TruSeq Indexed Adapters: P-GATCGGAAGAGCACACGTCTGAACTCCAGTCAC(INDEX)ATCTCGTATGCCGTCTTCTGCTT*G
- P5 primer: AATGATACGGCGACCACCGA*G
- P7 primer: CAAGCAGAAGACGGCATACGA*G
- 2X Rapid ligase buffer (Enzymatics #B101L)
- Enzymatics DNA ligase (Enzymatics #L6030-HC-L)
- 10% (wt/vol) Sodium dodecyl sulfate (SDS; Sigma-Aldrich, cat. no. L4509)
- Proteinase K (Thermo Fisher Scientific, cat. no. EO0492)
- Agencourt AMPure XP magnetic beads (Beckman Coulter, cat. no. A63880)
- 80% Ethanol (Decon Labs, cat. no. 2716)
- 5X KAPA buffer (KAPA #KK2502)
- KAPA HS HIFI polymerase (KAPA #KK2502)
- Qubit dsDNA HS kit (Life Technologies, cat. no. Q32851)

#### EQUIPMENT

- Centrifuge Eppendorf 5810, swinging bucket
- Centrifuge Eppendorf 5424, fixed angle rotor
- Centrifuge Eppendorf 5415R, refrigerated fixed angle rotor
- Macsimag magnetic separator (Miltenyi, cat. no. 130-092-168), which allows clean withdrawal of the liquid from the bottom of 1.7 and 2 mL microfuge tubes.
- Vortex mixer (e.g., VWR Vortex Genie)
- Micro-centrifuge (e.g., VWR Model V)
- 1.5-mL microcentrifuge tubes (Genesee, cat. no. 22-282)
- 2-mL microcentrifuge tubes (Axygen, cat. no. MCT-200-C)
- 0.6-mL microcentrifuge tubes (Axygen, cat. no. MCT-060-C)
- 0.6-mL low-retention microcentrifuge tubes (e.g. Thermo Fisher Scientific cat. no. 3446)
- Tube nutator (e.g BD Clay Adams Nutator Mixer; VWR, cat. no. 15172-203)
- Heater block with wells for 1.5-mL microcentrifuge tubes
- Water bath (set to 37 °C)
- Capillary electrophoresis instrument (e.g. Agilent Tapestation 4200)
- Thermal cycler with 3°C/sec ramp rate (e.g. Nexus Eco Thermal Cycler, Thermo Fisher Scientific E6332000029)
- Qubit Fluorometer (Life Technologies, cat. no. Q33216)
- Agilent 4200 TapeStation Instrument
- Biomek FX or FX^P^ equipped with a 96-channel pod and P200 head
- One 1 x 1 Tip Loader ALP (Beckman Coulter C02867)
- Three 1 x 3 Static ALPs (Beckman Coulter B87478)
- One Single Position Cooling/Heating ALP (Beckman Coulter 719361)
- 96S Super Magnet Plate (Alpaqua SKU A001322)
- Aluminum Heat Block Insert for PCR Plates (V&P Scientific, Inc. VP741I6A)
- Cooling Unit filled with antifreeze (e.g Thermo Neslab RTE-7 Digital One Recirculating Chiller Mfr # 271103200000)
- MicroAmp Support Bases (ThermoFisher N801-0531)
- 96 well LoBind PCR plates, Semi-skirted (Eppendorf # 0030129504)
- MicroAmp Clear Adhesive Film (Applied Biosystems Ref# 4306311)
- Biomek AP96 P250 Pre-Sterile Tips with barrier (Beckman Coulter # 717253)
- 96 well Polystyrene V-Bottom Microplates (Greiner Bio-One # 651101)
- MASTERBLOCK™ 96 Deep Well Conical Bottom 2 mL Storage Plates (Greiner Bio-One # 780271)
- 2-20 µL 8-Channel Multi Pipette (e.g. Rainin 17013803)
- 20-200 µL 8-Channel Multi Pipette (e.g. Rainin 17013805)
- Reagent Reservoirs with Dividers (Thermo Scientific 8095)

#### BIOMEK PROGRAMMING

##### Labware Type Editor

| Eppendorf 96 Well LoBind PCR Plate, Semi-skirted (96 Well PCR Plate) |  |  |  |  |
| --- | --- | --- | --- | --- |
| **Basic Info** |  | X | Y |  |
|  | Span: | 12.426 cm | 8.397 cm |  |
|  | Height: | 2.02 cm |  |  |
| **Movement Info** |  | X | Y | Z |
|  | Gripper Offset: | 0 | 0 | 1.6 |
|  | Gripper Squeeze: | 0.45 cm |  |  |
|  | Gripper Unsqueeze: | -1 cm |  |  |
|  | Speed Limit: | 50% |  |  |
| **Wells** |  | X | Y |  |
| **Well Configuration** | Well Offset: | 1.44 cm | 1.12 cm |  |
|  | Well Count: | 12 | 8 |  |
|  | Well Spacing: | 0.9 cm | 0.9 cm |  |
|  | Max Volume: | 235 µL |  |  |
|  | Shape: | Round |  |  |
|  | Upper Radius: | 0.28 cm |  |  |
|  | Lower Radius: | 0.28 cm |  |  |
|  | Height: | 0.5 cm |  |  |
|  | Bottom: | X |  |  |
|  | Shape: | Cone |  |  |
|  | Radius: | 0.28 cm |  |  |
|  | Height: | 1.4 cm |  |  |

| MicroAmp Support Base (PCR Plate Rack) |  |  |  |  |
| --- | --- | --- | --- | --- |
| **Basic Info** |  |  |  |  |
|  |  | X | Y |  |
|  | Span: | 12.7762 cm | 8.5471 cm |  |
|  | Height: | 1.4 cm |  |  |
| **Stacking** |  | X | Y | Z |
|  | Stack Offset: | 0 | 0 | 0.1 |
|  | Stacking Speed: | 100% |  |  |
| **Stack Offsets Edit** | 96 Well PCR Plate: | X | Stack Offset X: | 0 cm |
|  |  |  | Stack Offset Y: | 0.1 cm |
|  |  |  | Stack Offset Z: | 0.65 cm |
| **Wells** |  | X | Y |  |
| **Well Configuration** | Well Offset: | 1.438 cm | 1.123 cm |  |
|  | Well Count: | 12 | 8 |  |
|  | Well Spacing: | 0.9 cm | 0.9 cm |  |
|  | Max Volume: | 250 uL |  |  |
|  | Shape: | Round |  |  |
|  | Upper Radius: | 0.3 cm |  |  |
|  | Lower Radius: | 0.28 cm |  |  |
|  | Height: | 1 cm |  |  |
|  | Bottom: | X |  |  |
|  | Shape: | Cone |  |  |
|  | Radius: | 0.28 cm |  |  |
|  | Height: | 1.4 cm |  |  |

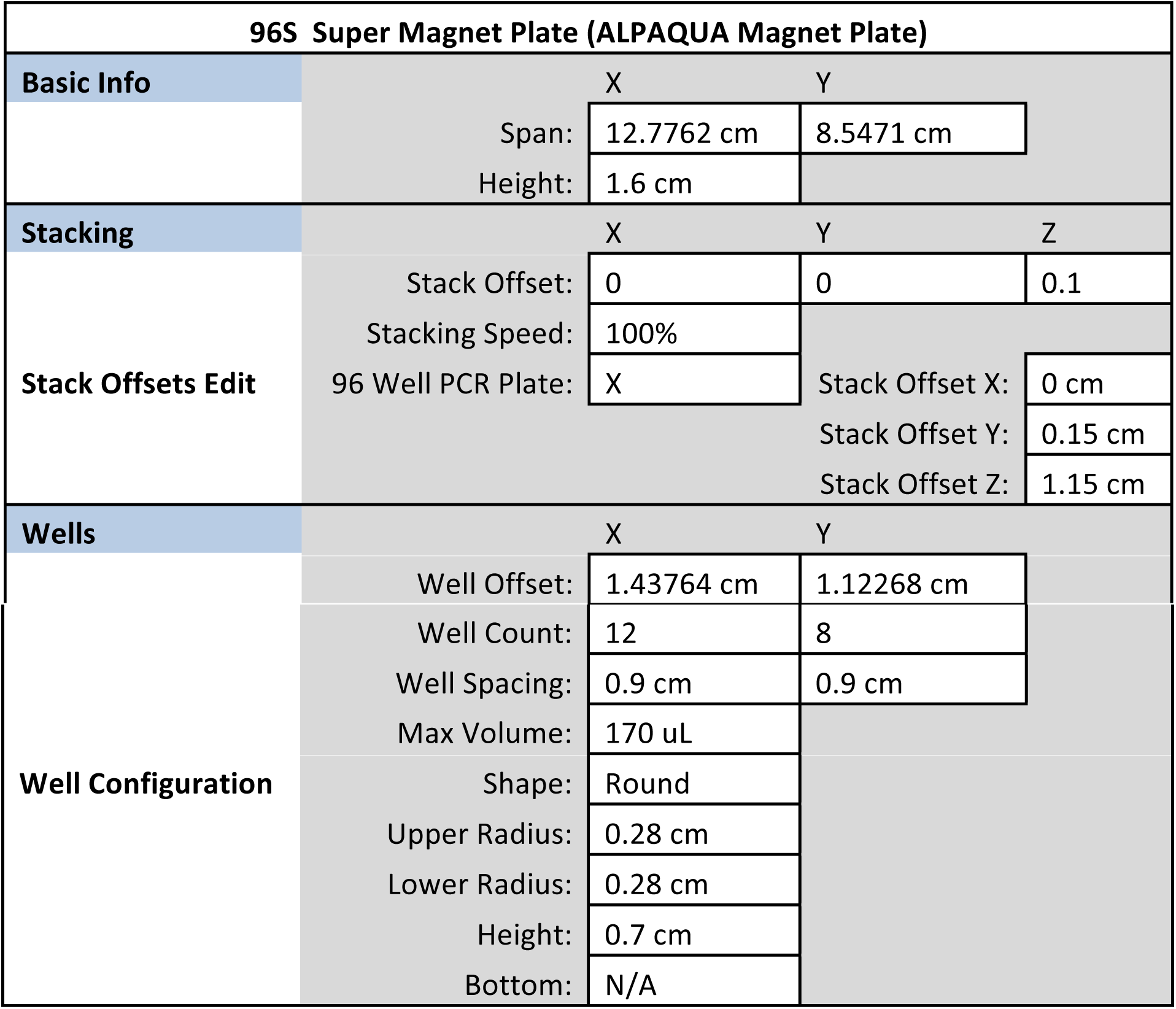

| 96S Super Magnet Plate (ALPAQUA Magnet Plate) |  |  |  |  |
| --- | --- | --- | --- | --- |
| **Basic Info** |  |  |  |  |
|  |  | X | Y |  |
|  | Span: | 12.7762 cm | 8.5471 cm |  |
|  | Height: | 1.6 cm |  |  |
| **Stacking** |  | X | Y | Z |
|  | Stack Offset: | 0 | 0 | 0.1 |
|  | Stacking Speed: | 100% |  |  |
| **Stack Offsets Edit** | 96 Well PCR Plate: | X | Stack Offset X: | 0 cm |
|  |  |  | Stack Offset Y: | 0.15 cm |
|  |  |  | Stack Offset Z: | 1.15 cm |
| **Wells** |  | X | Y |  |
|  | Well Offset: | 1.43764 cm | 1.12268 cm |  |

|  |  |  |  |
| --- | --- | --- | --- |
| **Well Configuration** | Well Count: | 12 | 8 |
|  | Well Spacing: | 0.9 cm | 0.9 cm |
|  | Max Volume: | 170 uL |  |
|  | Shape: | Round |  |
|  | Upper Radius: | 0.28 cm |  |
|  | Lower Radius: | 0.28 cm |  |
|  | Height: | 0.7 cm |  |
|  | Bottom: | N/A |  |

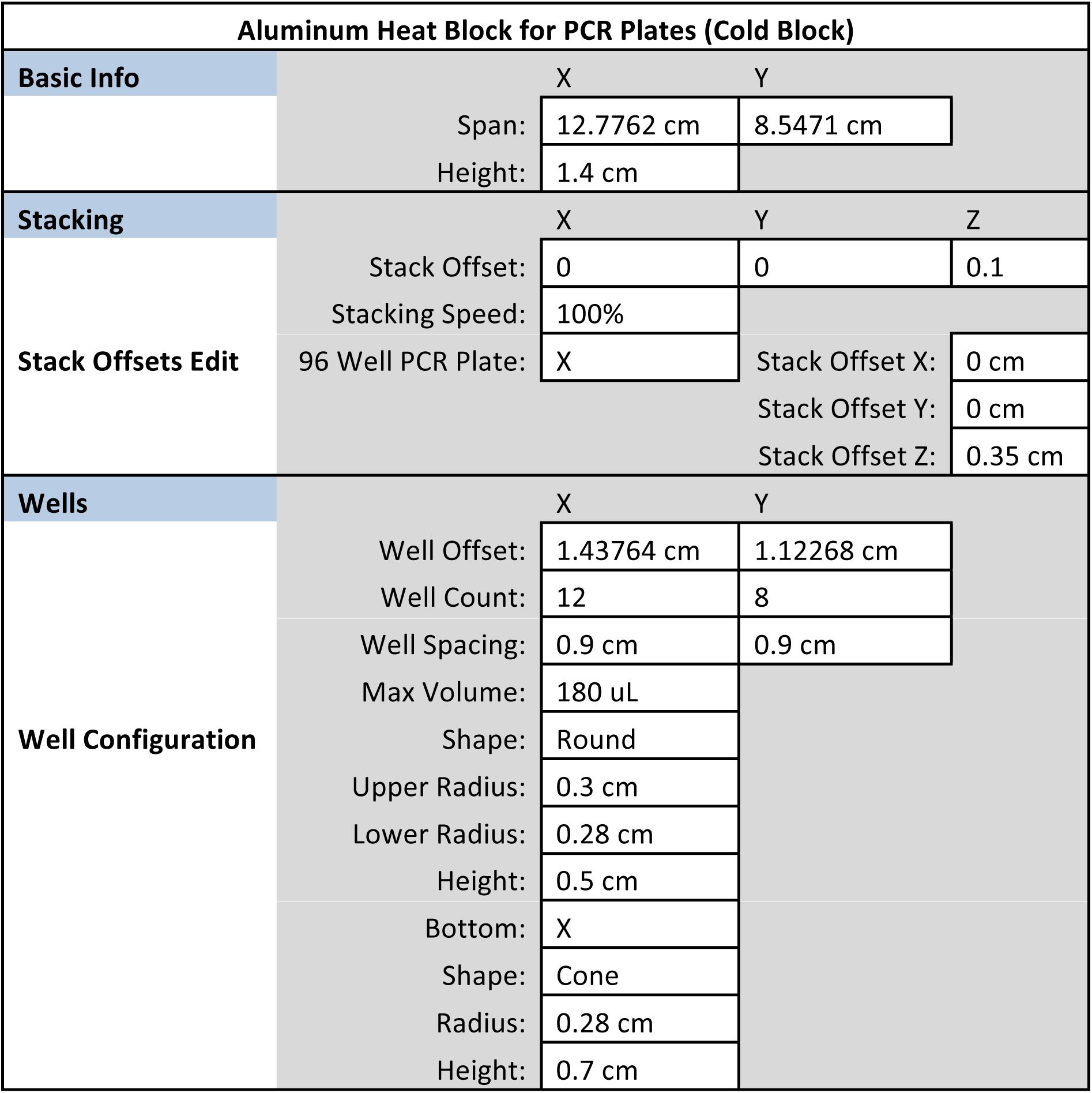

| 96 Well Polystyrene V-Bottom Microplate (V-Bottom Plate) |  |  |  |
| --- | --- | --- | --- |
| **Basic Info** |  | X | Y |
|  | Span: | 12.78 cm | 8.56 cm |
|  | Height: | 1.41 cm |  |

| Wells |  | X | Y |
| --- | --- | --- | --- |
| Well Configuration | Well Offset: | 1.44 cm | 1.12 cm |
|  | Well Count: | 12 | 8 |
|  | Well Spacing: | 0.9 cm | 0.9 cm |
|  | Max Volume: | 235 uL |  |
|  | Shape: | Round |  |
|  | Upper Radius: | 0.309 cm |  |
|  | Lower Radius: | 0.2735 cm |  |
|  | Height: | 0.83 cm |  |
|  | Bottom: | X |  |
|  | Shape: | Cone |  |
|  | Radius: | 0.2735 cm |  |
|  | Height: | 0.2 cm |  |

| 96 Deep Well 2 mL Plate (Deep Well Plate) |  |  |  |
| --- | --- | --- | --- |
| Basic Info |  | X | Y |
|  | Span: | 12.78 cm | 8.56 cm |
|  | Height: | 4.4 cm |  |
| Wells |  | X | Y |
| Well Configuration | Well Offset: | 1.44 cm | 1.13 cm |
|  | Well Count: | 12 | 8 |
|  | Well Spacing: | 0.9 cm | 0.9 cm |
|  | Max Volume: | 2300 uL |  |
|  | Shape: | Rectangle |  |

| Biomek AP96 P250 barrier tips (AP96_200uL) |  |  |  |
| --- | --- | --- | --- |
| Basic Info |  | X | Y |
|  | Span: | 12.789 cm | 8.56 cm |
|  | Height: | 5.6 cm |  |

| Tips |  | X | Y |
| --- | --- | --- | --- |
|  | Tip Offset: | 1.447 cm | 1.132 cm |
|  | Tip Count: | 12 | 8 |
|  | Tip Spacing: | 0.9 cm | 0.9 cm |
|  | Tip Load Z Offset: | -5.3 cm |  |
|  | Tip Unload Z Offset: | 0 cm |  |
|  | Tip Type: | P200 |  |

##### Liquid Type Editor

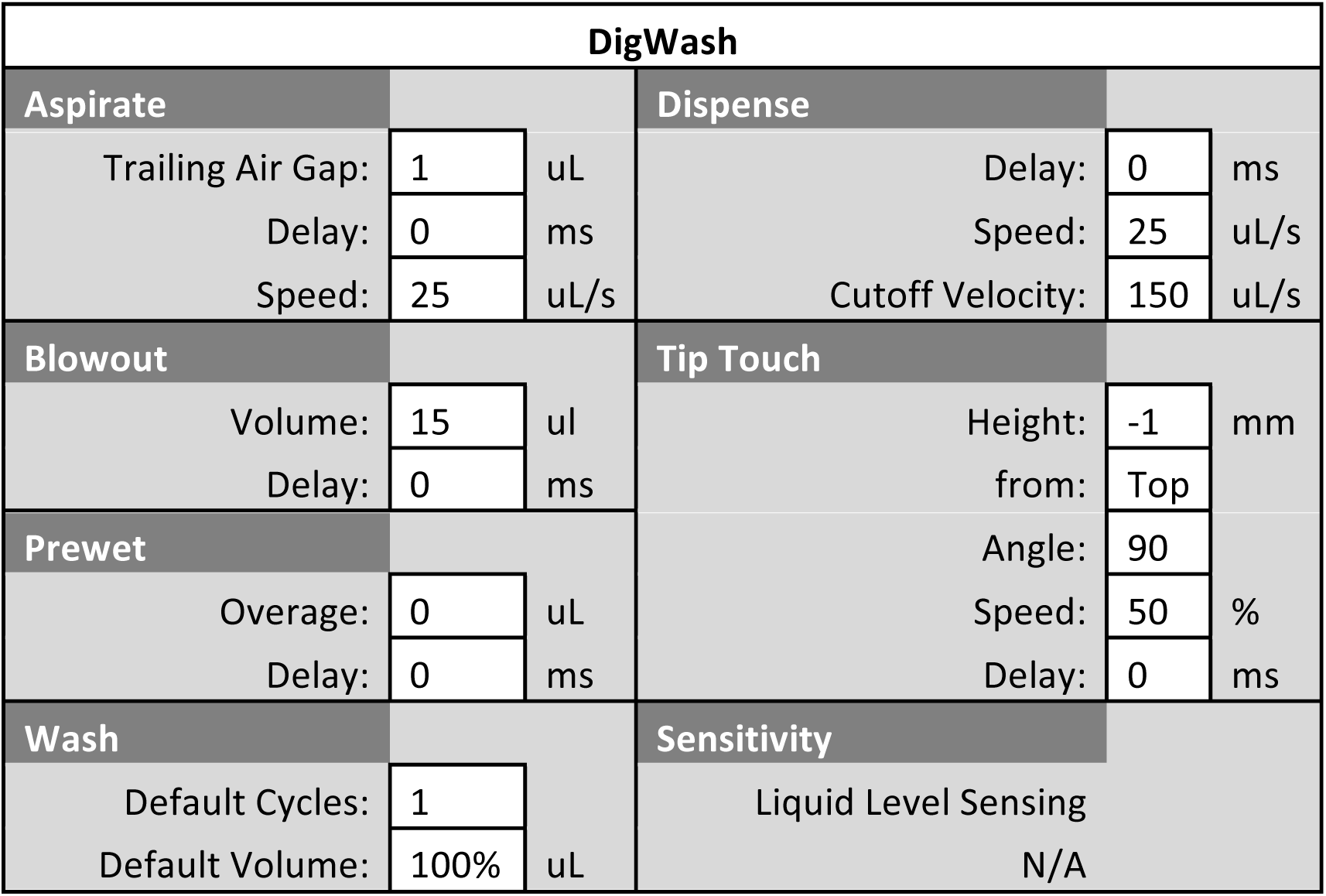

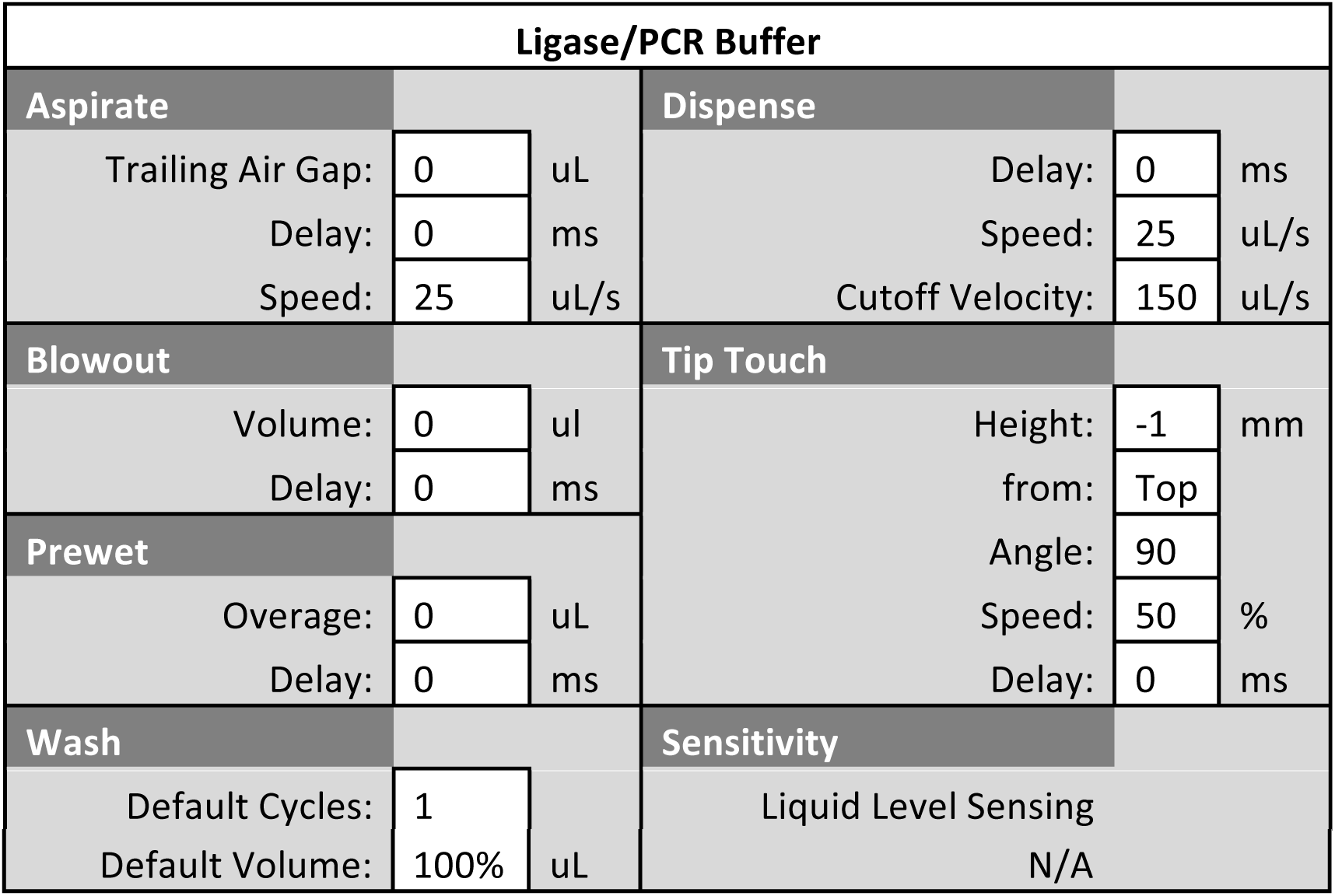

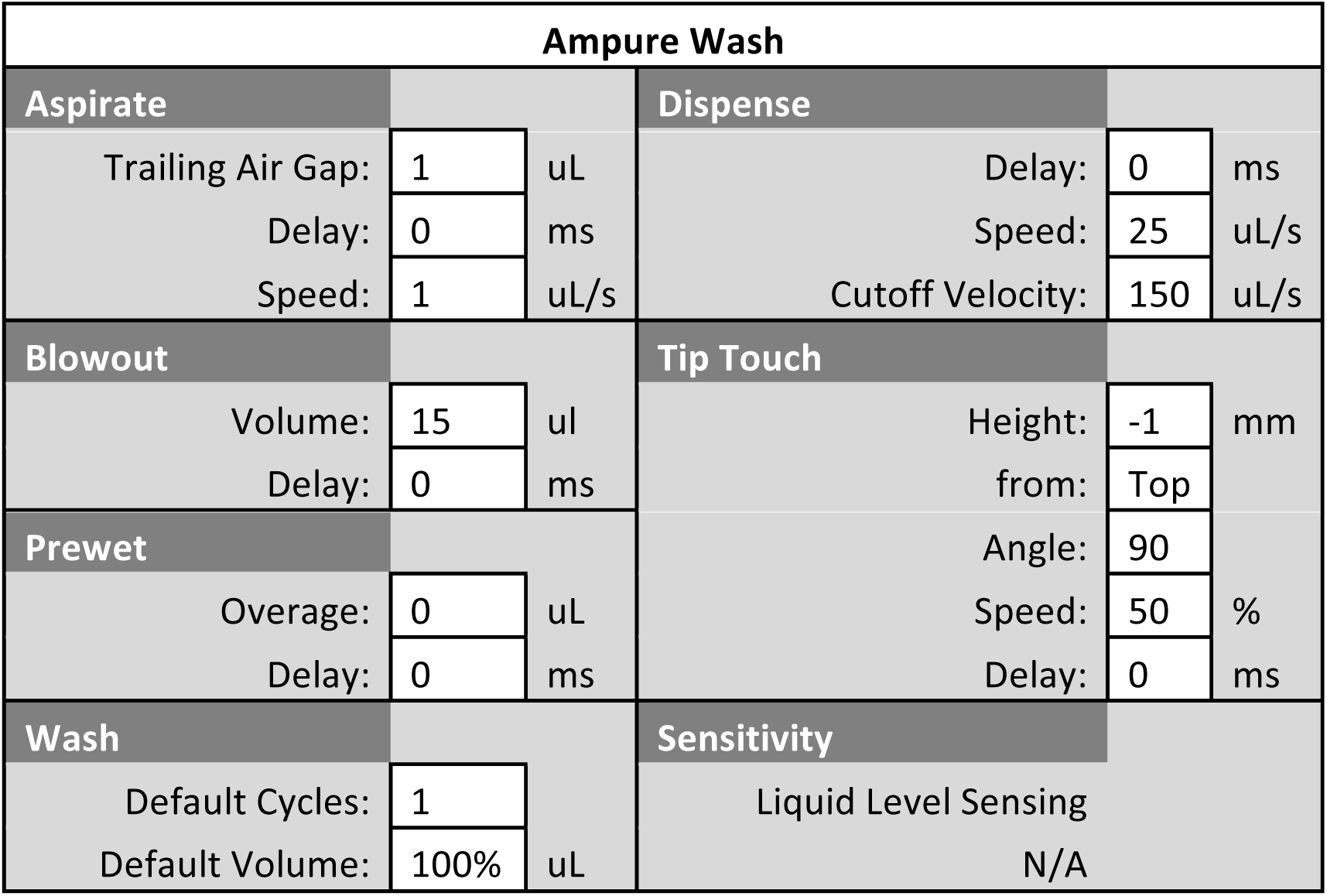

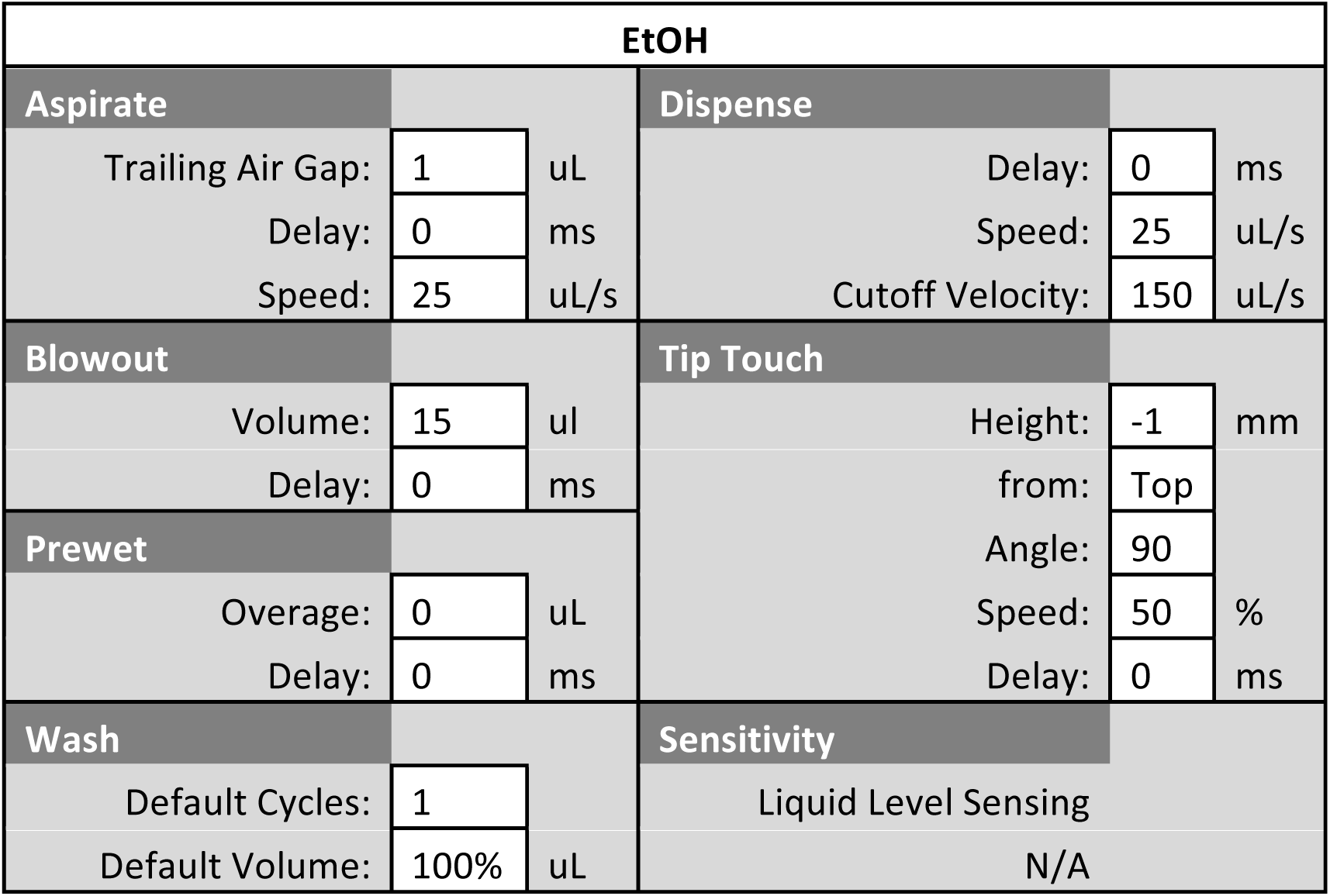

| Tris-HCl-1 |  |  |  |  |  |
| --- | --- | --- | --- | --- | --- |
| Aspirate |  |  | Dispense |  |  |
| Trailing Air Gap: | 1 | uL | Delay: | 0 | ms |
| Delay: | 0 | ms | Speed: | 25 | uL/s |
| Speed: | 25 | uL/s | Cutoff Velocity: | 150 | uL/s |
| **Blowout** |  |  | **Tip Touch** |  |  |
| Volume: | 15 | ul | Height: | -1 | mm |
| Delay: | 0 | ms | from: | Top |  |
| **Prewet** |  |  | Angle: | 90 |  |
| Overage: | 0 | uL | Speed: | 50 | % |
| Delay: | 0 | ms | Delay: | 0 | ms |
| **Wash** |  |  | **Sensitivity** |  |  |
| Default Cycles: | 1 |  | Liquid Level Sensing |  |  |
| Default Volume: | 100% | uL | N/A |  |  |

| Tris-HCl-2 |  |  |  |  |  |
| --- | --- | --- | --- | --- | --- |
| **Aspirate** |  |  | **Dispense** |  |  |
| Trailing Air Gap: | 1 | uL | Delay: | 0 | ms |
| Delay: | 0 | ms | Speed: | 25 | uL/s |
| Speed: | 1 | uL/s | Cutoff Velocity: | 150 | uL/s |
| **Blowout** |  |  | **Tip Touch** |  |  |
| Volume: | 15 | ul | Height: | -1 | mm |
| Delay: | 0 | ms | from: | Top |  |
| **Prewet** |  |  | Angle: | 90 |  |
| Overage: | 0 | uL | Speed: | 50 | % |
| Delay: | 0 | ms | Delay: | 0 | ms |
| **Wash** |  |  | **Sensitivity** |  |  |
| Default Cycles: | 1 |  | Liquid Level Sensing |  |  |
| Default Volume: | 100% | uL | N/A |  |  |

##### Techniques

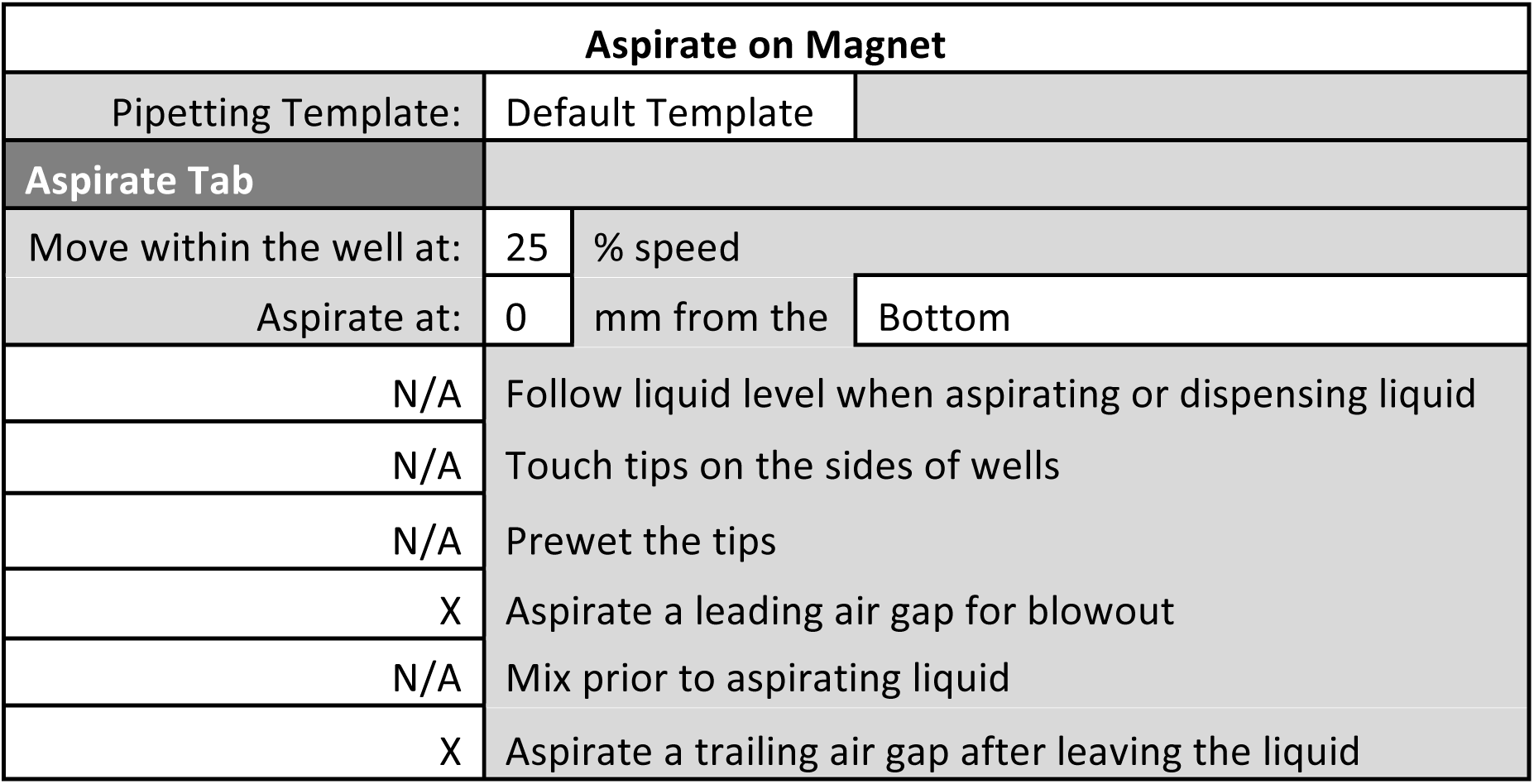

| Dispense in Waste |  |  |  |
| --- | --- | --- | --- |
| Pipetting Template: | Default Template |  |  |
| Dispense Tab |  |  |  |
| Move within the Well at: | 100 | % speed |  |
| Dispense at: | 30 | mm from the | Bottom |
| N/A | Follow liquid level when aspirating or dispensing liquid |  |  |
| X | Touch tips on the sides of the wells |  |  |
| X | Blowout all leading air gaps |  |  |
| N/A | Mix after dispensing |  |  |

| Aspirate from DWP |  |  |  |
| --- | --- | --- | --- |
| Pipetting Template: | Default Template |  |  |
| Aspirate Tab |  |  |  |
| Move within the well at: | 25 | % speed |  |
| Aspirate at: | 0 | mm from the | Bottom |
| N/A | Follow liquid level when aspirating or dispensing liquid |  |  |
| N/A | Touch tips on the sides of wells |  |  |
| N/A | Prewet the tips |  |  |
| X | Aspirate a leading air gap for blowout |  |  |
| N/A | Mix prior to aspirating liquid |  |  |
| X | Aspirate a trailing air gap after leaving the liquid |  |  |

| Dispense on Magnet |  |  |  |
| --- | --- | --- | --- |
| Pipetting Template: | Default Template |  |  |
| Dispense Tab |  |  |  |
| Move within the Well at: | 25 | % speed |  |
| Dispense at: | 2 | mm from the | Bottom |
| N/A | Follow liquid level when aspirating or dispensing liquid |  |  |
| X | Touch tips on the sides of the wells |  |  |
| X | Blowout all leading air gaps |  |  |
| N/A | Mix after dispensing |  |  |

| Aspirate from VBP |  |
| --- | --- |
| Pipetting Template: | Default Template |

| Aspirate tab |  |  |  |
| --- | --- | --- | --- |
| Move within the well at: | 25 | % speed |  |
| Aspirate at: | 1.5 | mm from the | Bottom |
| N/A | Follow liquid level when aspirating or dispensing liquid |  |  |
| N/A | Touch tips on the sides of wells |  |  |
| N/A | Prewet the tips |  |  |
| X | Aspirate a leading air gap for blowout |  |  |
| N/A | Mix prior to aspirating liquid |  |  |
| X | Aspirate a trailing air gap after leaving the liquid |  |  |

| Dispense and Mix-1 |  |  |  |  |  |  |  |  |  |
| --- | --- | --- | --- | --- | --- | --- | --- | --- | --- |
| Pipetting Template: | Default Template |  |  |  |  |  |  |  |  |
| Dispense Tab |  |  |  |  |  |  |  |  |  |
| Move within the Well at: | 25 | % speed |  |  |  |  |  |  |  |
| Dispense at: | 5 | mm from the |  | Bottom |  |  |  |  |  |
| N/A | Follow liquid level when aspirating or dispensing liquid |  |  |  |  |  |  |  |  |
| N/A | Touch tips on the sides of the wells |  |  |  |  |  |  |  |  |
| X | Blowout all leading air gaps |  |  |  |  |  |  |  |  |
| X | Mix after dispensing liquid |  |  |  |  |  |  |  |  |
| Mix: | 30 | ul |  | 10 |  | times |  |  |  |
| Aspirate at: | 5 | mm from the |  | Bottom |  | at |  | 25 | uL/s |
| Dispense at: | 5 | mm from the |  | Bottom |  | at |  | 25 | uL/s |

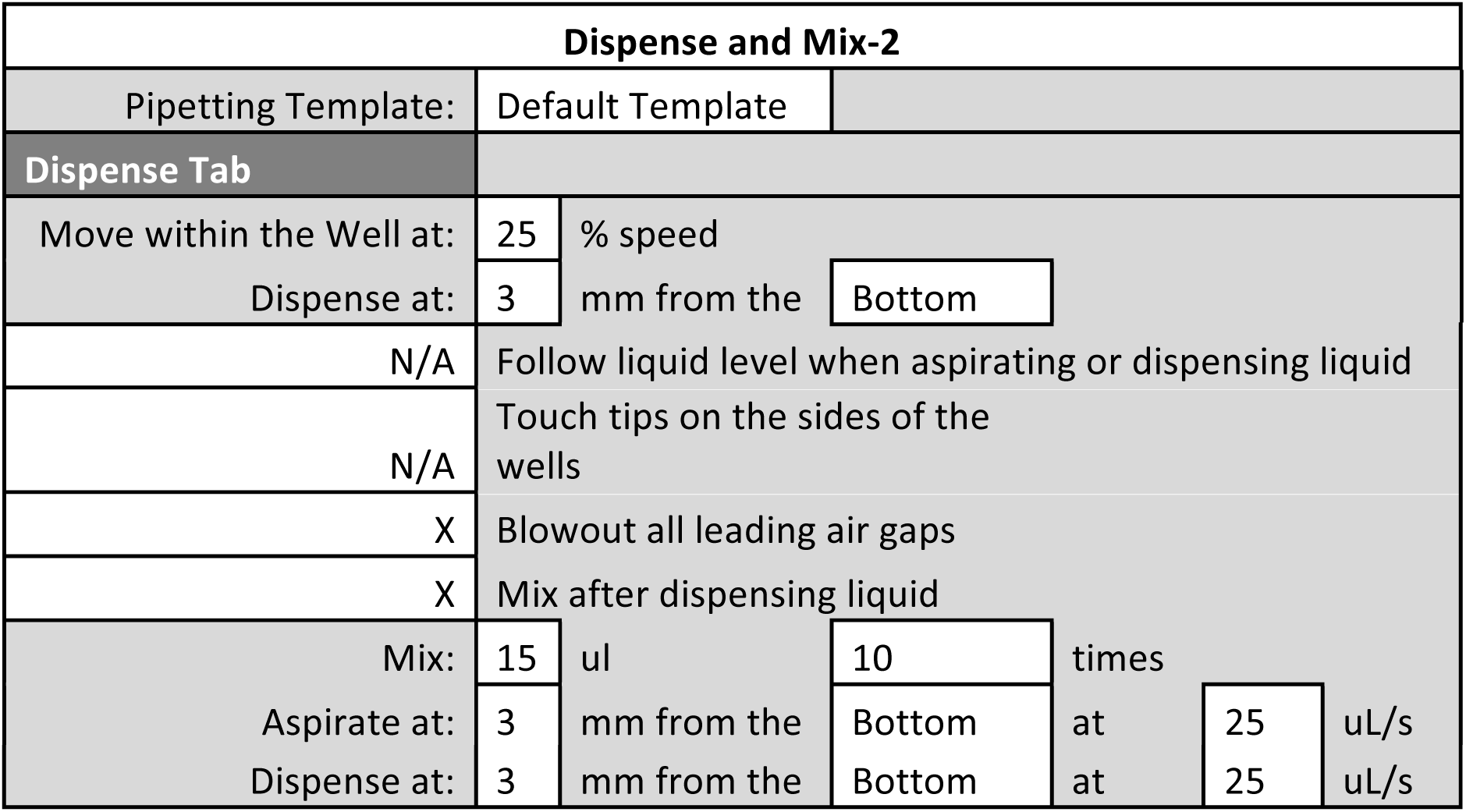

| Dispense and Mix-2 |  |  |  |  |  |  |  |  |  |
| --- | --- | --- | --- | --- | --- | --- | --- | --- | --- |
| Pipetting Template: | Default Template |  |  |  |  |  |  |  |  |
| Dispense Tab |  |  |  |  |  |  |  |  |  |
| Move within the Well at: | 25 | % speed |  |  |  |  |  |  |  |
| Dispense at: | 3 | mm from the |  | Bottom |  |  |  |  |  |
| N/A | Follow liquid level when aspirating or dispensing liquid |  |  |  |  |  |  |  |  |
| N/A | Touch tips on the sides of the wells |  |  |  |  |  |  |  |  |
| X | Blowout all leading air gaps |  |  |  |  |  |  |  |  |
| X | Mix after dispensing liquid |  |  |  |  |  |  |  |  |
| Mix: | 15 | ul |  | 10 |  | times |  |  |  |
| Aspirate at: | 3 | mm from the |  | Bottom |  | at |  | 25 | uL/s |

|  |  |  |  |  |  |  |
| --- | --- | --- | --- | --- | --- | --- |
| Dispense at: | 3 | mm from the | Bottom | at | 25 | uL/s |

| Dispense and Mix-3 |  |  |  |  |  |  |
| --- | --- | --- | --- | --- | --- | --- |
| Pipetting Template: | Default Template |  |  |  |  |  |
| Dispense Tab |  |  |  |  |  |  |
| Move within the Well at: | 25 | % speed |  |  |  |  |
| Dispense at: | 3 | mm from the | Bottom |  |  |  |
| N/A | Follow liquid level when aspirating or dispensing liquid |  |  |  |  |  |
| N/A | Touch tips on the sides of the wells |  |  |  |  |  |
| X | Blowout all leading air gaps |  |  |  |  |  |
| X | Mix after dispensing liquid |  |  |  |  |  |
| Mix: | 25 | ul | 10 | times |  |  |
| Aspirate at: | 3 | mm from the | Bottom | at | 25 | uL/s |
| Dispense at: | 3 | mm from the | Bottom | at | 25 | uL/s |

| Aspirate on Rack |  |  |  |
| --- | --- | --- | --- |
| Pipetting Template: | Default Template |  |  |
| Aspirate tab |  |  |  |
| Move within the well at: | 25 | % speed |  |
| Aspirate at: | 0.5 | mm from the | Bottom |
| N/A | Follow liquid level when aspirating or dispensing liquid |  |  |
| N/A | Touch tips on the sides of wells |  |  |
| N/A | Prewet the tips |  |  |
| N/A | Aspirate a leading air gap for blowout |  |  |
| N/A | Mix prior to aspirating liquid |  |  |
| N/A | Aspirate a trailing air gap after leaving the liquid |  |  |

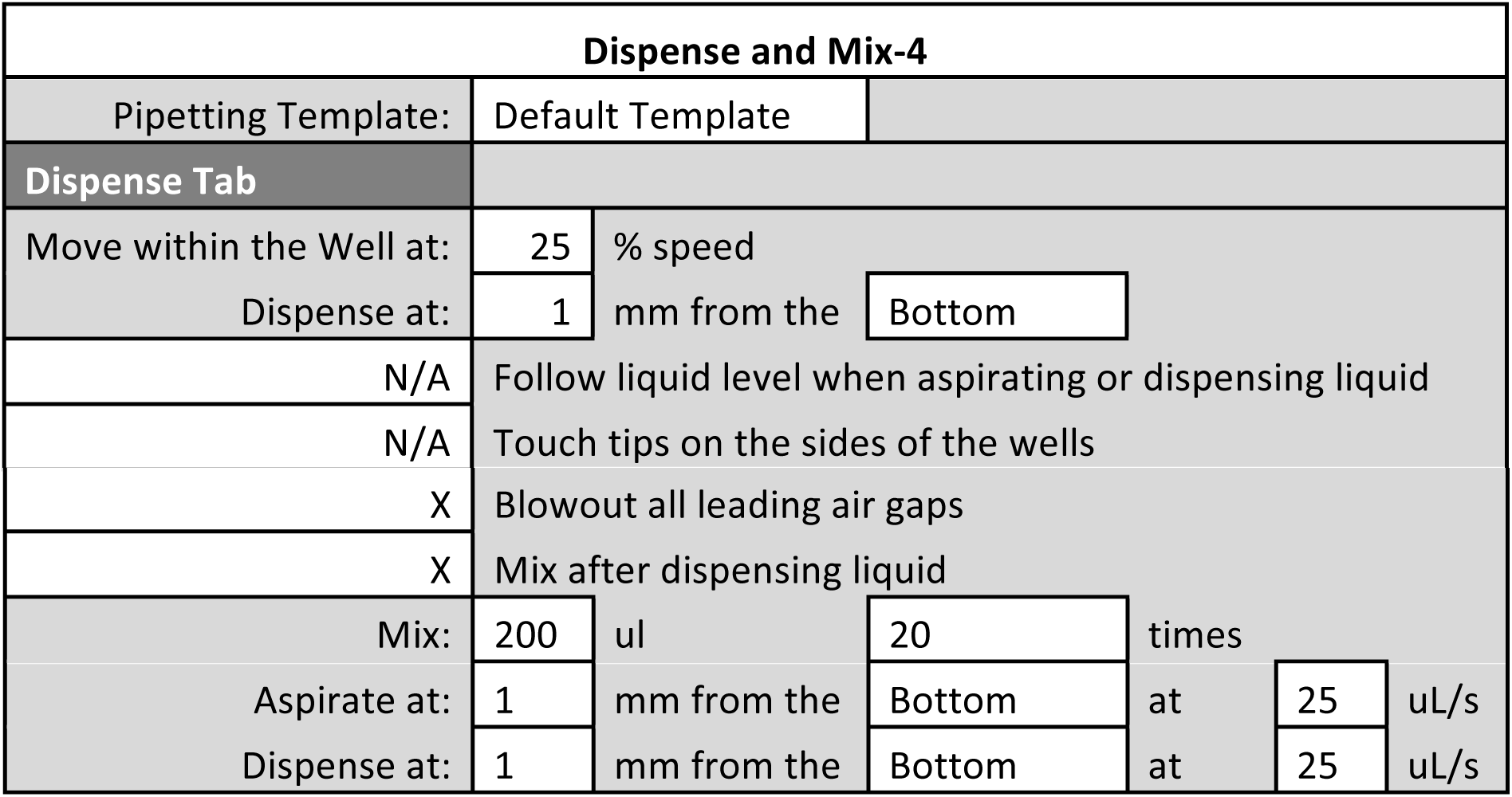

| Dispense and Mix-5 |  |  |  |  |  |  |
| --- | --- | --- | --- | --- | --- | --- |
| Pipetting Template: | Default Template |  |  |  |  |  |
| Dispense Tab |  |  |  |  |  |  |
| Move within the Well at: | 25 | % speed |  |  |  |  |
| Dispense at: | 1 | mm from the | Bottom |  |  |  |
| N/A | Follow liquid level when aspirating or dispensing liquid |  |  |  |  |  |
| X | Touch tips on the sides of the wells |  |  |  |  |  |
| X | Blowout all leading air gaps |  |  |  |  |  |
| X | Mix after dispensing liquid |  |  |  |  |  |
| Mix: | 30 | ul | 10 | times |  |  |
| Aspirate at: | 1 | mm from the | Bottom | at | 50 | uL/s |
| Dispense at: | 1 | mm from the | Bottom | at | 50 | uL/s |

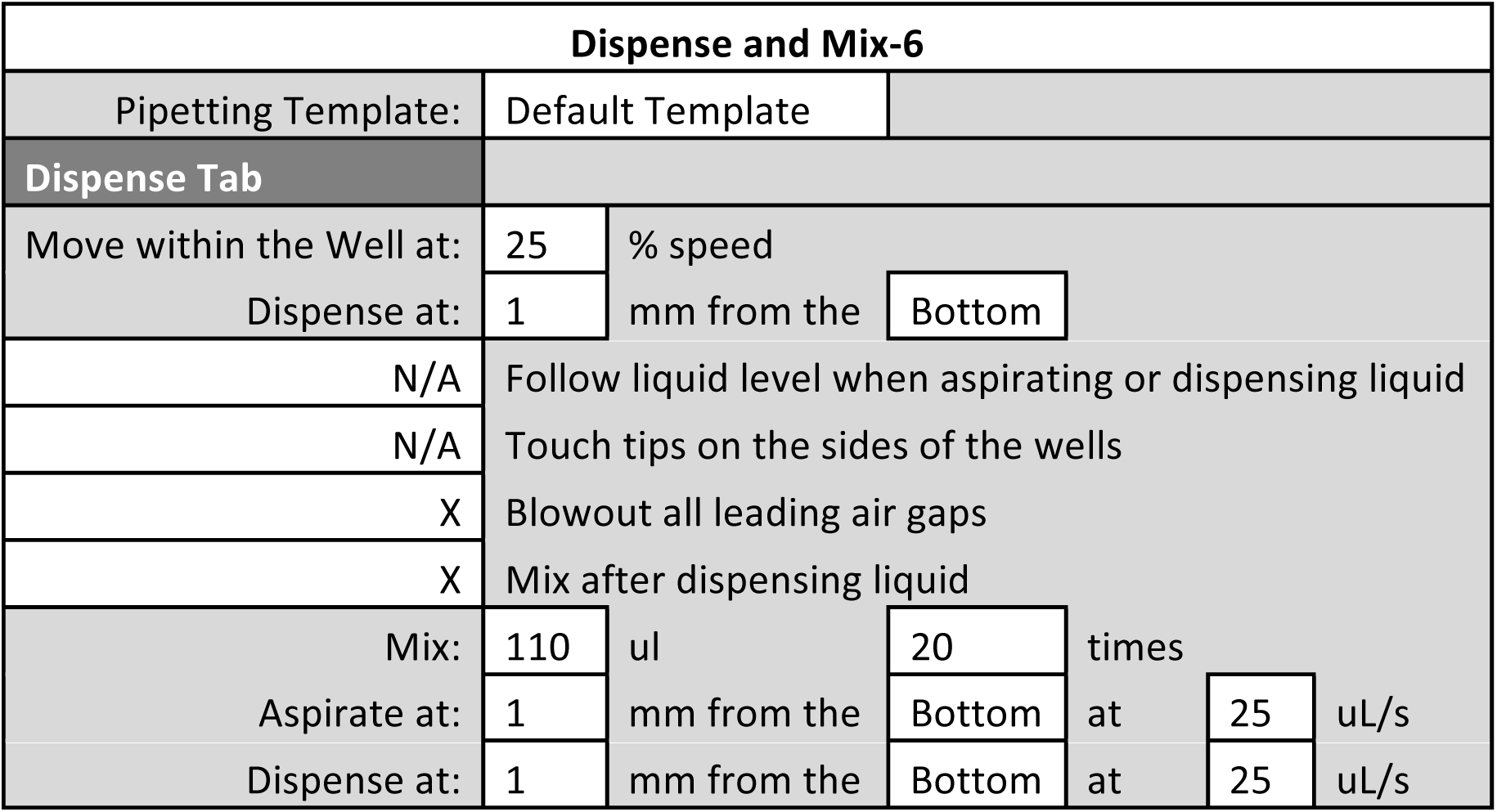

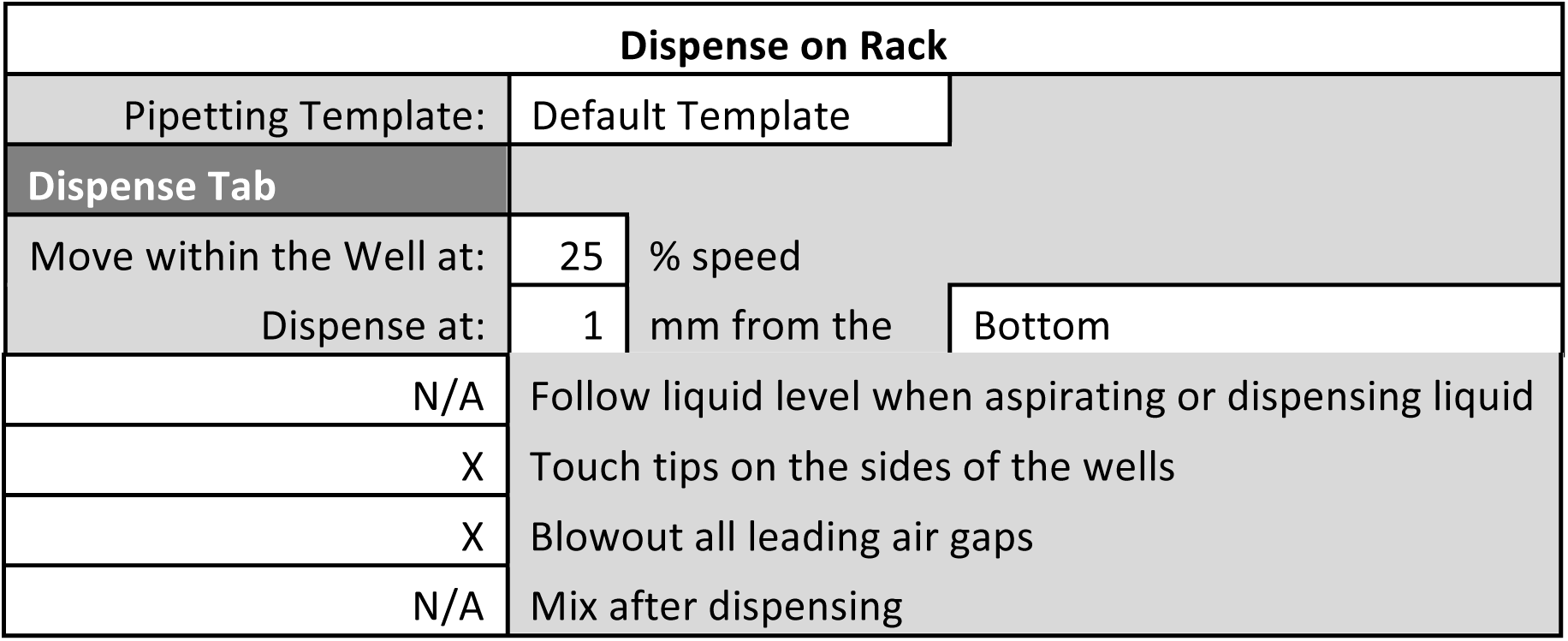

| Dispense on Rack |  |  |  |
| --- | --- | --- | --- |
| Pipetting Template: | Default Template |  |  |
| Dispense Tab |  |  |  |
| Move within the Well at: | 25 | % speed |  |
| Dispense at: | 1 | mm from the | Bottom |

|  |  |
| --- | --- |
| N/A | Follow liquid level when aspirating or dispensing liquid |
| X | Touch tips on the sides of the wells |
| X | Blowout all leading air gaps |
| N/A | Mix after dispensing |

### Methods

#### pA-MNase Binding

**1) Start**

**2) Instrument Setup**

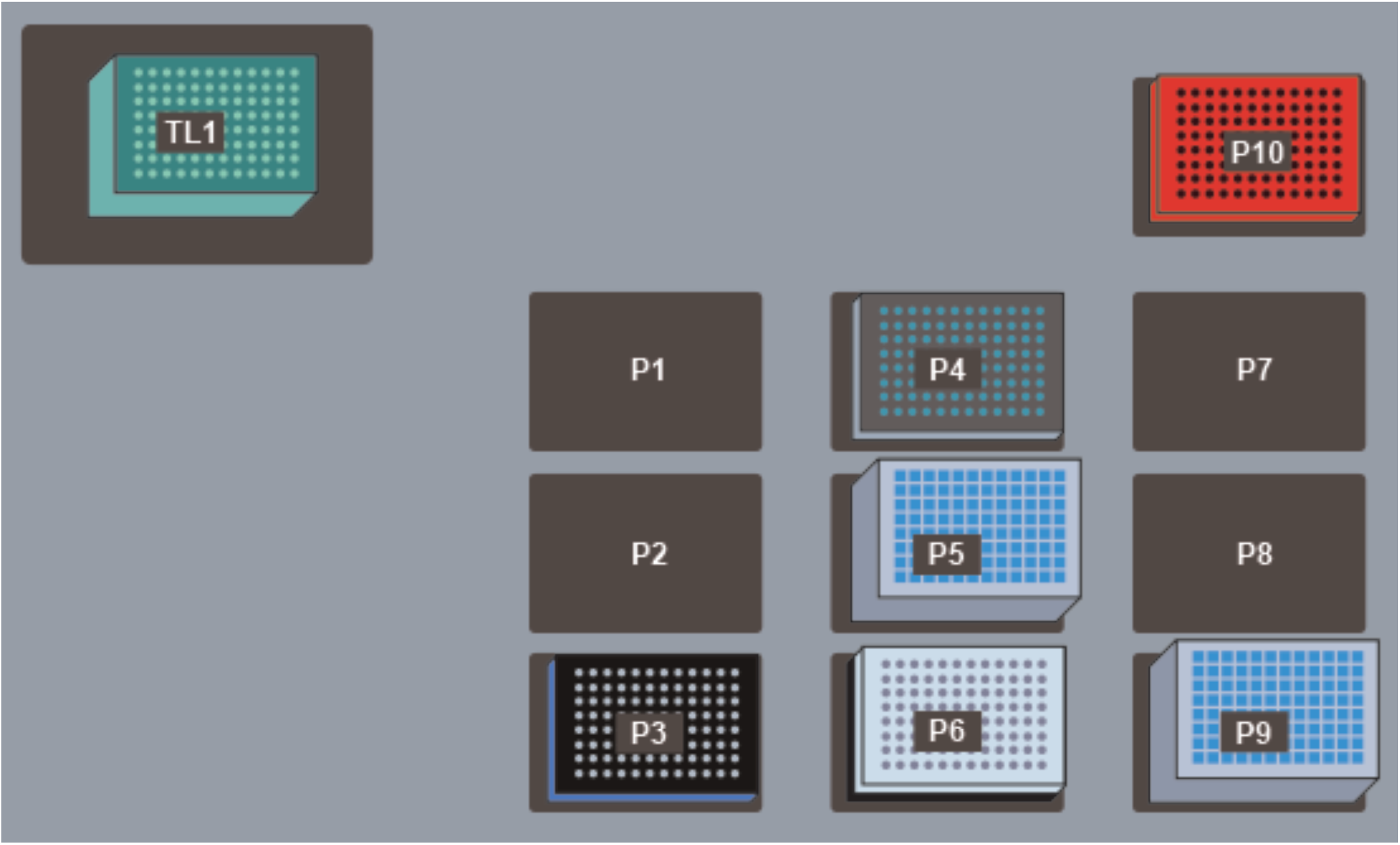

TL1: Fresh AP96 200 µL Tips (double click to increase the # of load times)

P3: ALPAQUA Magnet Plate

P4: V-Bottom Plate preloaded with 175 µL pA-MNase solution

P5: Deep Well Plate preloaded with 1 mL of Dig-Wash Buffer

P6: 96 Well PCR Plate preloaded with up to 150 µL of ConA bead-bound cells + Antibody stacked on a PCR Plate Rack

P9: Deep Well Plate for receiving liquid waste

P10: Cold Block seated on a Cooling/Heating ALP routed to a Cooling Unit pre-chilled to 0°C

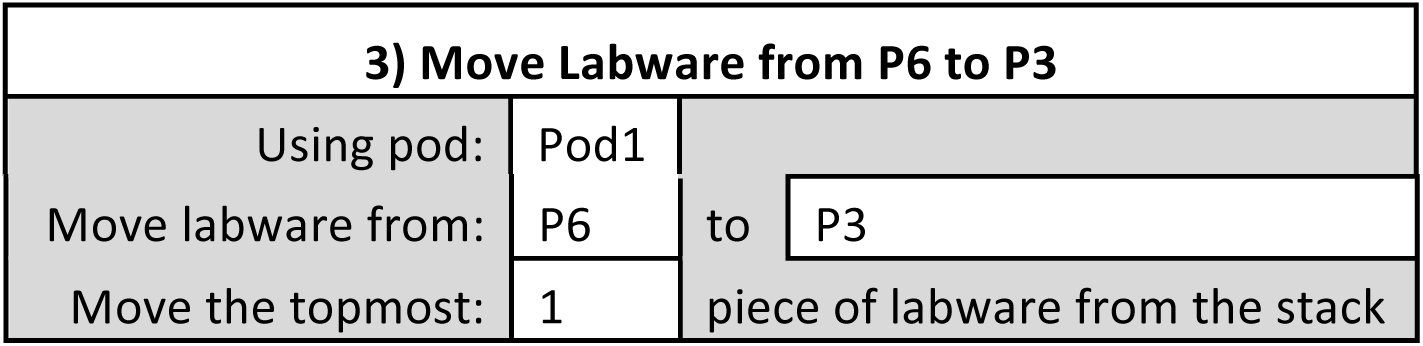

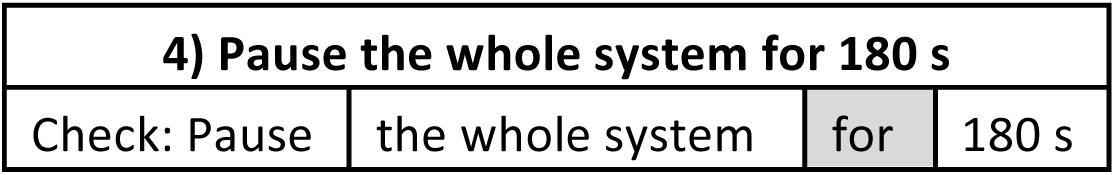

|  |  |
| --- | --- |
| **5) Load tips from TL1** |  |
| Tips: | Load |
| Location: | TL1 |
| Pod: | Pod1 |

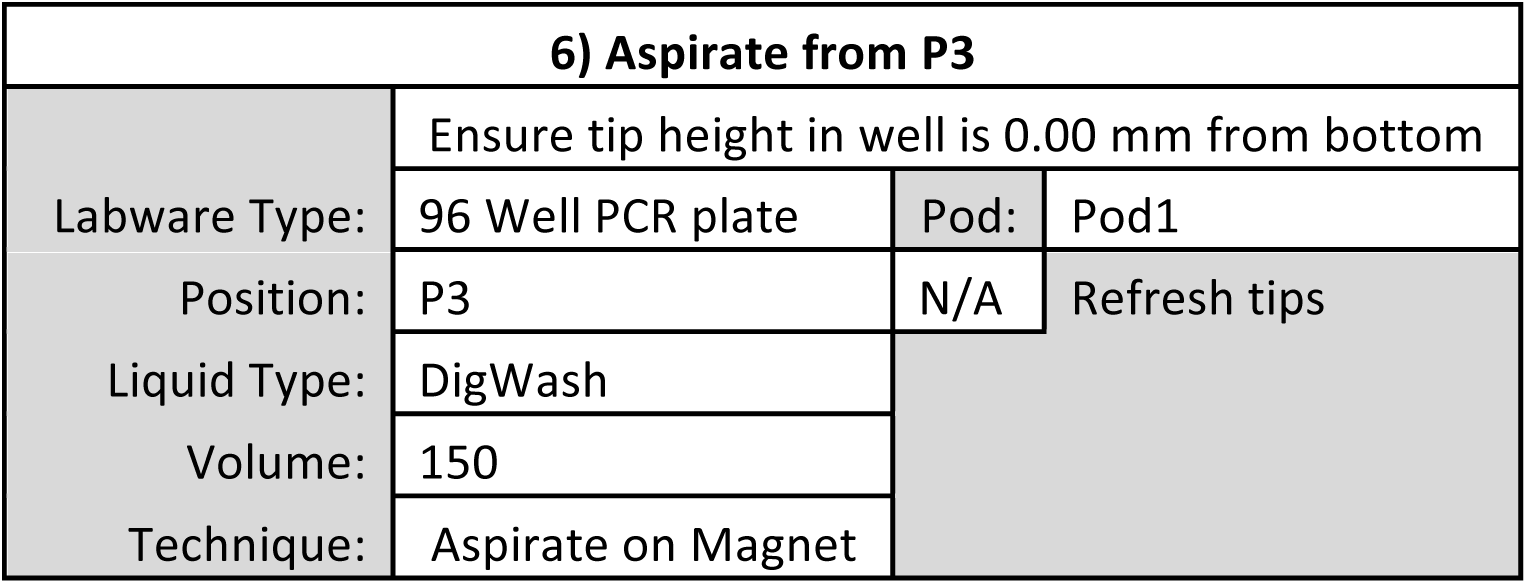

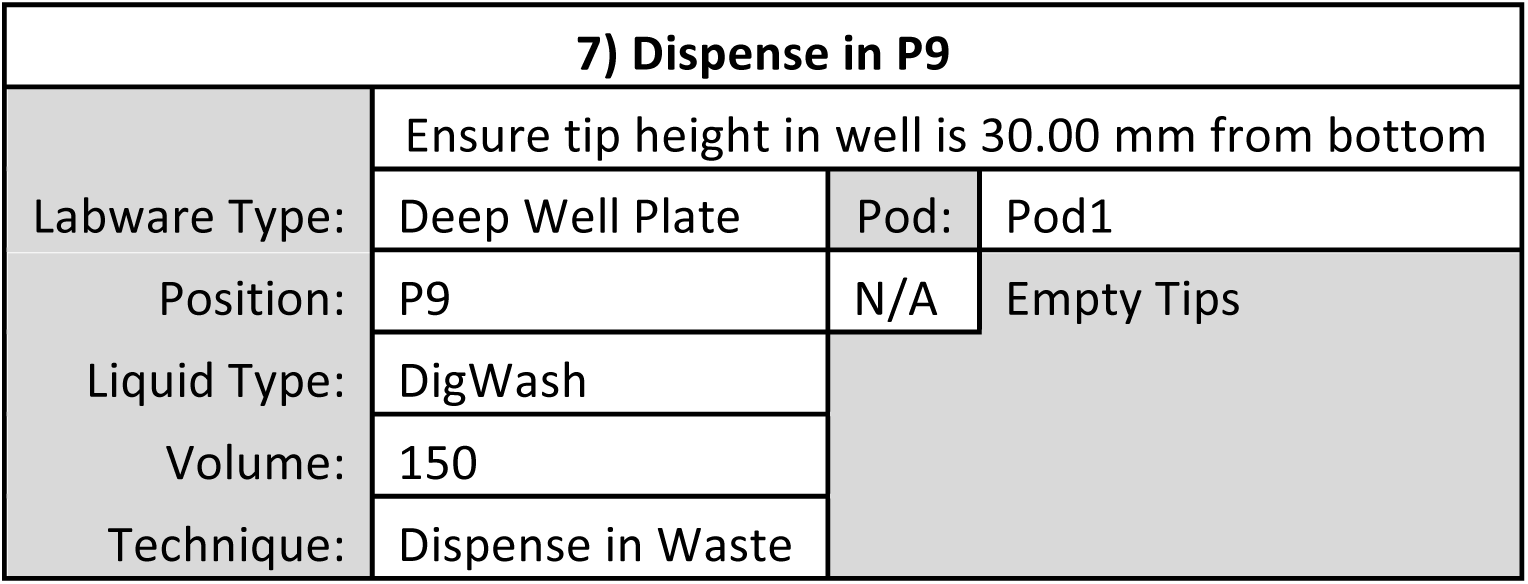

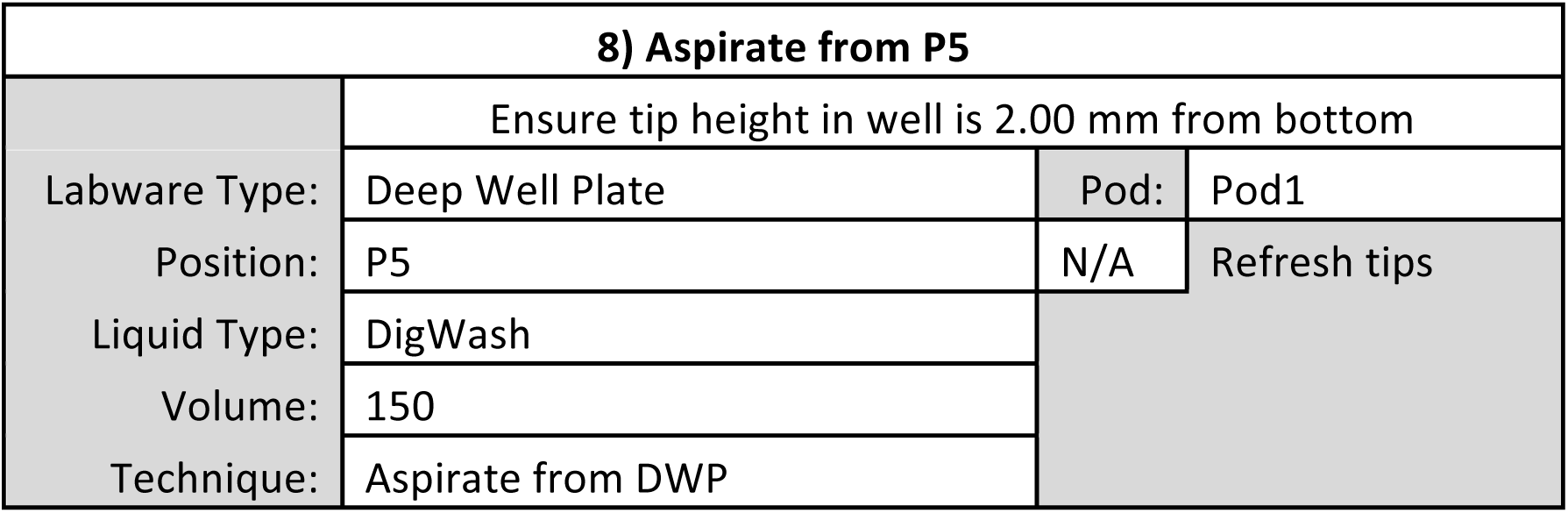

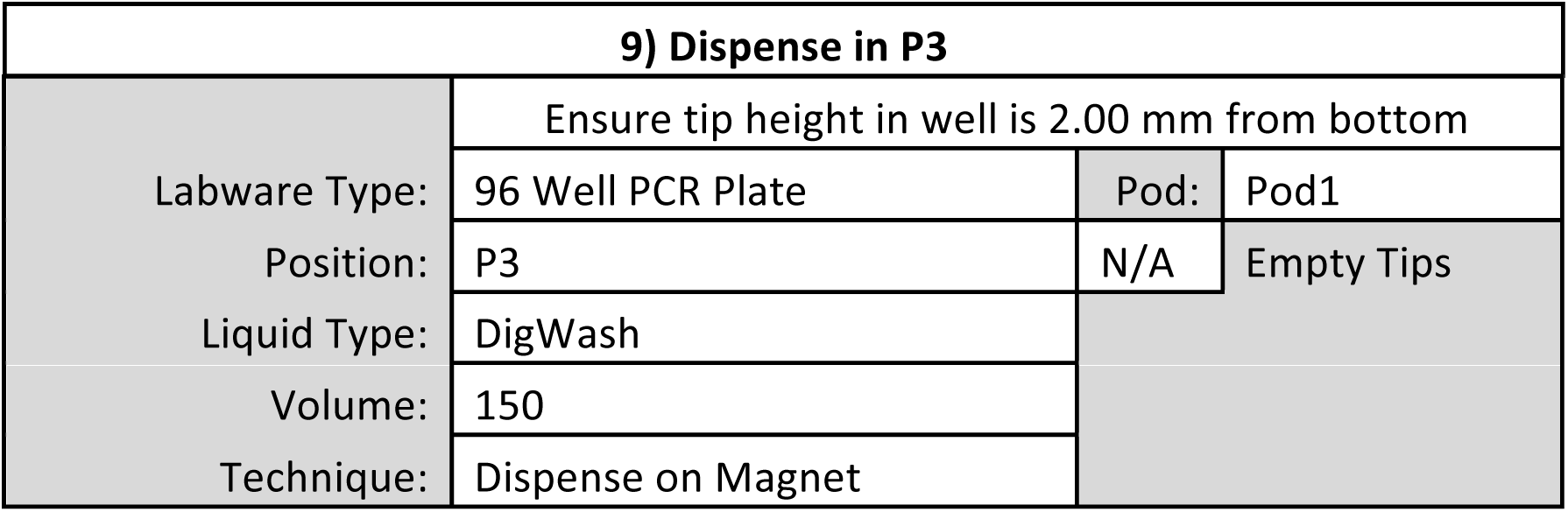

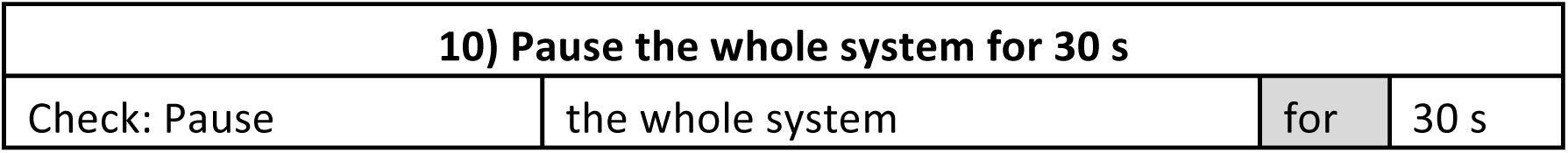

**11) Repeat 6-10 to wash cells a second time**

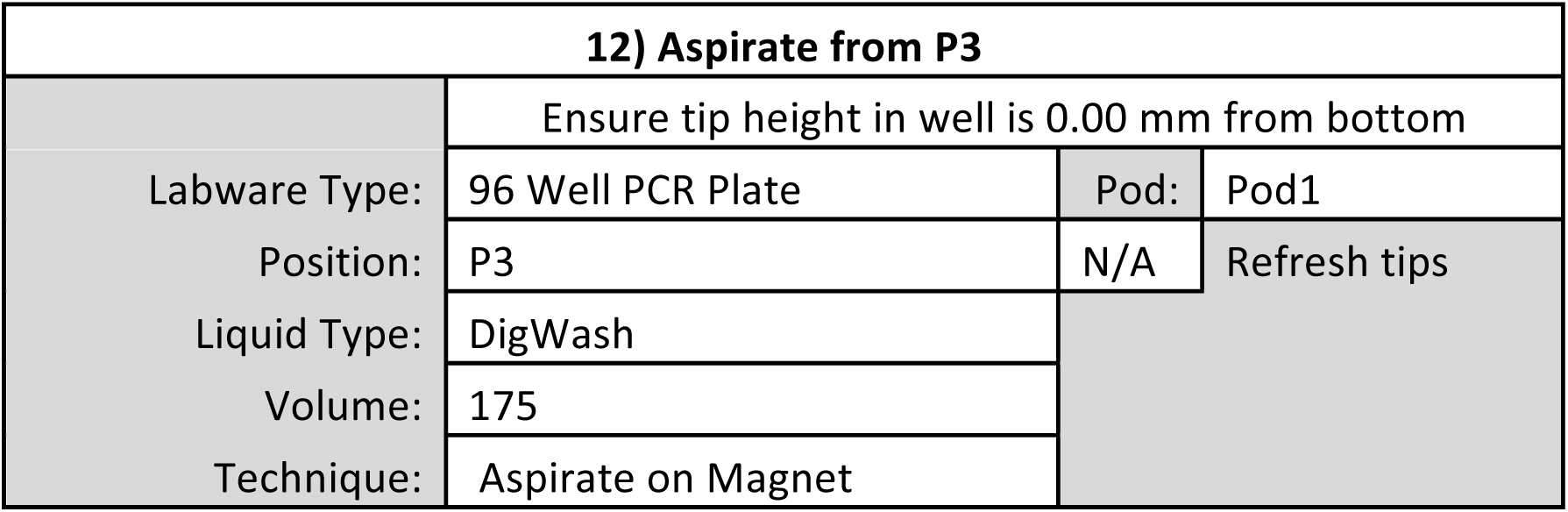

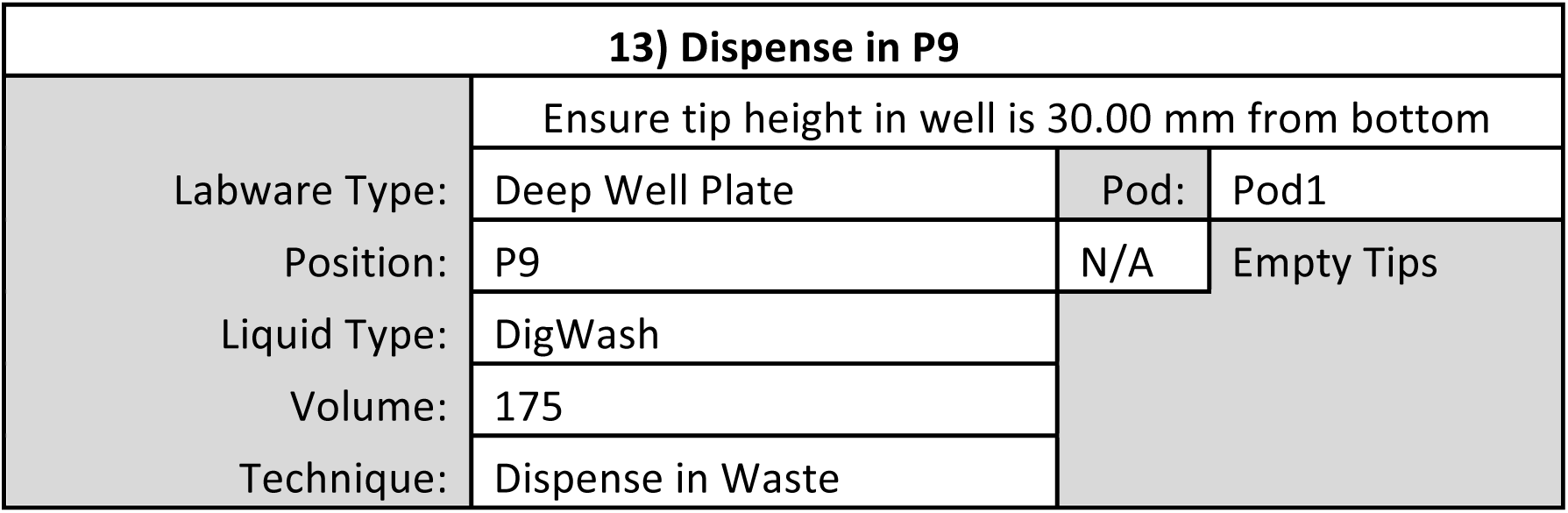

|  |  |
| --- | --- |
| **14) Unload tips to TL1** |  |
| Tips: | Unload |
| Location: | TL1 |
| Pod: | Pod1 |

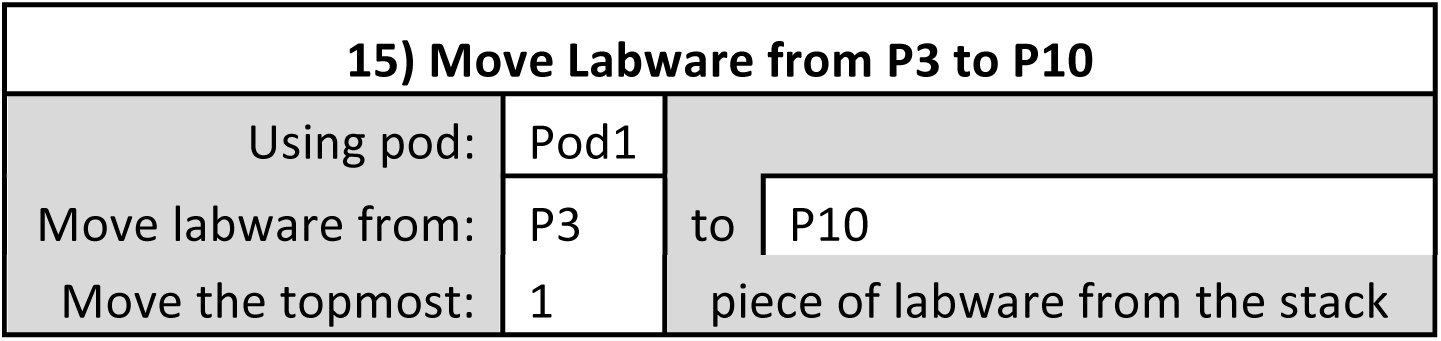

|  |  |  |  |
| --- | --- | --- | --- |
| **15) Move Labware from P3 to P10** |  |  |  |
| Using pod: | Pod1 |  |  |
| Move labware from: | P3 | to | P10 |

|  |  |  |
| --- | --- | --- |
| Move the topmost: | 1 | piece of labware from the stack |

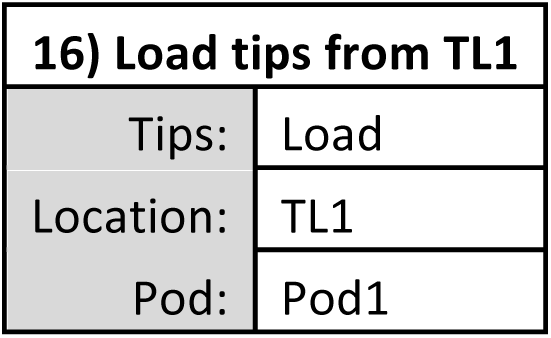

17) Aspirate from P4
|  |  |  |  |
| --- | --- | --- | --- |
|  | Ensure tip height in well is 1.5 mm from bottom |  |  |
| Labware Type: | V-Bottom Plate | Pod: | Pod1 |
| Position: | P4 | N/A | Refresh tips |
| Liquid Type: | DigWash |  |  |
| Volume: | 150 |  |  |
| Technique: | Aspirate from VBP |  |  |

18) Dispense in P10
|  |  |  |  |
| --- | --- | --- | --- |
|  | Ensure tip height in well is 5.00 mm from bottom |  |  |
| Labware Type: | 96 Well PCR plate | Pod: | Pod1 |
| Position: | P10 | N/A | Empty Tips |
| Liquid Type: | DigWash |  |  |
| Volume: | 150 |  |  |
| Technique: | Dispense and Mix-1 |  |  |

19) Pause the whole system for 510 s
|  |  |  |  |
| --- | --- | --- | --- |
| Check: Pause | the whole system | for | 510 s |

20) Mix in P10
|  |  |  |  |
| --- | --- | --- | --- |
|  | Ensure tip height in well is 5.00 mm from bottom |  |  |
| Labware Type: | 96 Well PCR plate | Pod | Pod1 |
| Position: | P10 | N/A | Refresh tips |
| Liquid Type: | DigWash | Mix | 25 times |
| Volume: | 30 | uL |  |
| Technique: | Custom |  |  |
| Customize |  |  |  |
| Pipetting Template: | Default Template |  |  |
| Mix Tab |  |  |  |

|  |  |  |  |  |  |
| --- | --- | --- | --- | --- | --- |
| Move within the well at: | 25 | % speed |  |  |  |
| Aspirate at: | 5 | mm from the | Bottom | at | 25 ul/s |
| Dispense at: | 5 | mm from the | Bottom | at | 25 ul/s |
| N/A | Follow liquid level when aspirating or dispensing liquid |  |  |  |  |
| N/A | Touch tips on the sides of the wells |  |  |  |  |
| N/A | Aspirate a leading air gap prior to mix and blowout after mix is complete |  |  |  |  |

**21) Repeat Steps 19&20 four times in order to mix cells in pA-MNase solution for ∼1hr**

**21) Repeat Steps 19&20 four times in order to mix cells in pA-MNase solution for ~1hr**
| **22) Unload tips to TL1** |  |
| --- | --- |
| Tips: | Unload |
| Location: | TL1 |
| Pod: | Pod1 |

| **23) Finish** |
| --- |
| Check: Clear current instrument setup of all labware after the method completes |
| Check: Clear current device setup of all labware after the method completes |
| Check: Unload disposable tips from all pods after the method completes |
| Check: Clear all global variables after the method completes |

**pA-MNase Digest**

**1) Start**

**2) Instrument Setup**

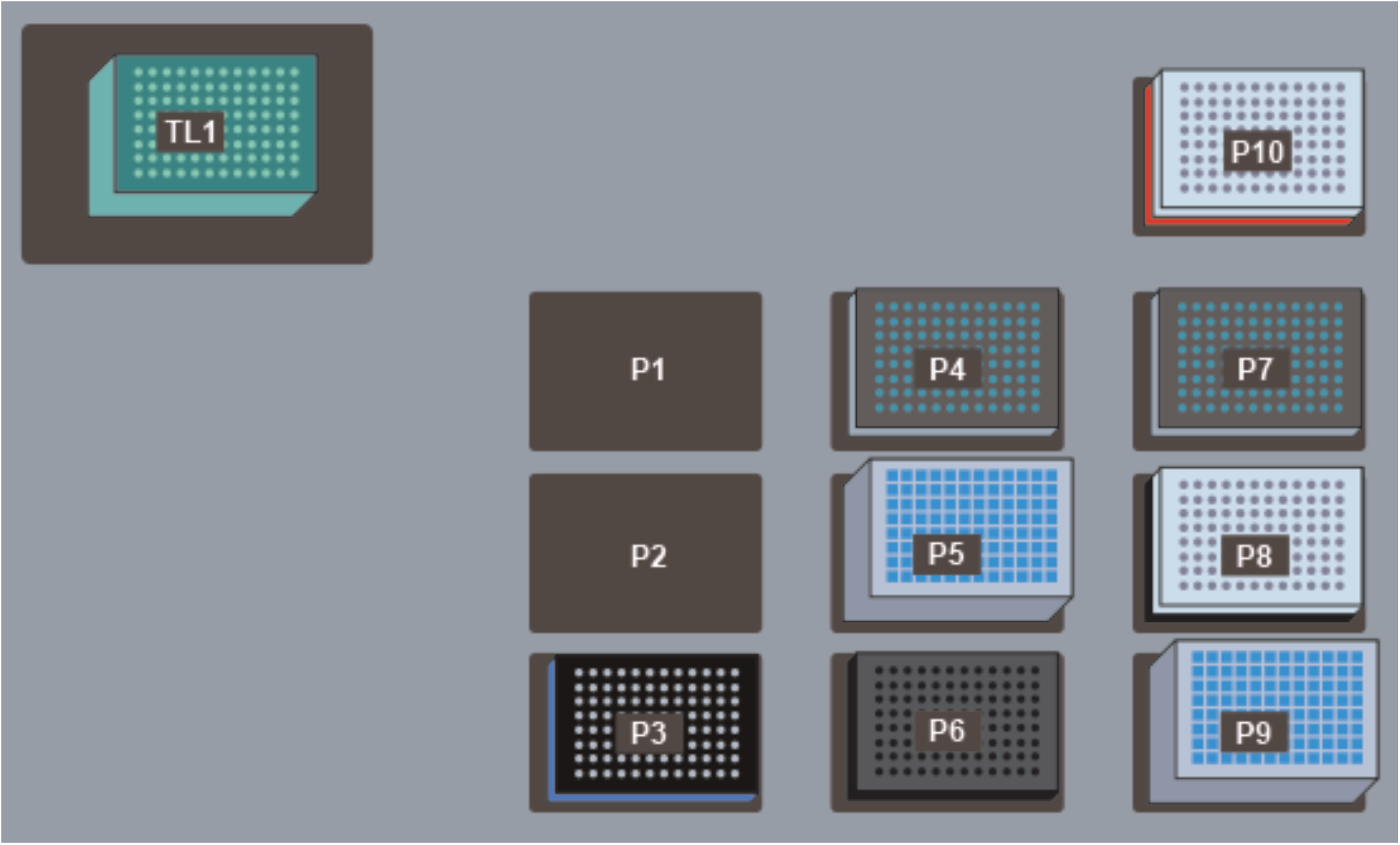

TL1: Continue with same AP96 200 µL Tips (double click to increase the # of load times)

P3: ALPAQUA Magnet Plate

P4: V-Bottom Plate preloaded with 50 µL pA-MNase Reaction Mix

P5: Deep Well Plate containing leftover Dig-Wash buffer from pA-MNase Binding Method

P6: PCR Plate Rack

P7: V-Bottom Plate preloaded with 25 µL 4X STOP Buffer

P8: 96 Well PCR Plate for accepting digested chromatin stacked on a PCR Plate Rack

P9: Deep Well Plate for receiving liquid waste

P10: 96 Well PCR Plate preloaded with 150 µL Sample in pA-MNase solution on a Cold Block seated on a Cooling/Heating ALP routed to a Cooling Unit pre-chilled to 0°C

| 3) Move Labware from P10 to P3 |  |  |  |
| --- | --- | --- | --- |
| Using pod: | Pod1 |  |  |
| Move labware from: | P10 | to | P3 |
| Move the topmost: | 1 | piece of labware from the stack |  |

| 4) Pause the whole system for 180 s |  |  |  |
| --- | --- | --- | --- |
| Check: Pause | the whole system | for | 180 s |

| 5) Load tips from TL1 |  |
| --- | --- |
| Tips: | Load |
| Location: | TL1 |
| Pod: | Pod1 |

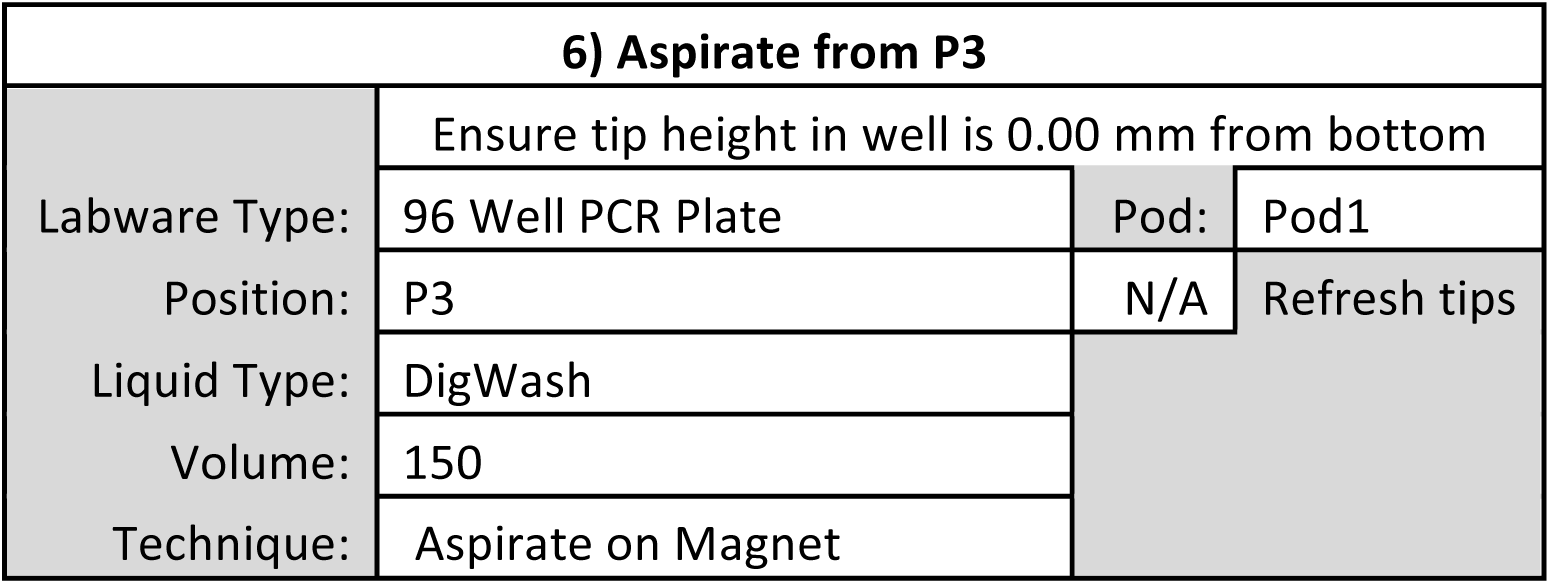

| 7) Dispense in P9 |  |  |  |
| --- | --- | --- | --- |
|  | Ensure tip height in well is 30.00 mm from bottom |  |  |
| Labware Type: | Deep Well Plate | Pod: | Pod1 |
| Position: | P9 | N/A | Empty Tips |
| Liquid Type: | DigWash |  |  |
| Volume: | 150 |  |  |
| Technique: | Dispense in Waste |  |  |

| 8) Aspirate from P5 |  |  |  |
| --- | --- | --- | --- |
|  | Ensure tip height in well is 2.00 mm from bottom |  |  |
| Labware Type: | Deep Well Plate | Pod: | Pod1 |
| Position: | P5 | N/A | Refresh tips |
| Liquid Type: | DigWash |  |  |
| Volume: | 150 |  |  |
| Technique: | Aspirate from DWP |  |  |

| 9) Dispense in P3 |  |  |  |
| --- | --- | --- | --- |
|  | Ensure tip height in well is 2.00 mm from bottom |  |  |
| Labware Type: | 96 Well PCR Plate | Pod: | Pod1 |
| Position: | P3 | N/A | Empty Tips |
| Liquid Type: | DigWash |  |  |
| Volume: | 150 |  |  |
| Technique: | Dispense on Magnet |  |  |

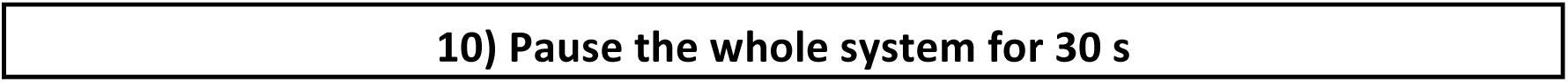
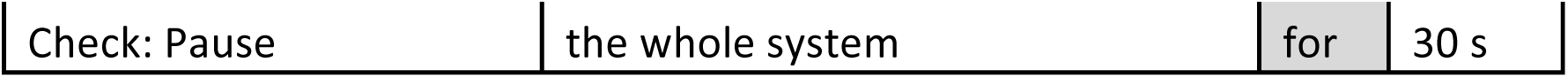

|  |
| --- |
| 10) Pause the whole system for 30 s |

|  |  |  |  |
| --- | --- | --- | --- |
| Check: Pause | the whole system | for | 30 s |

**11) Repeat Steps 6-10 to wash cells a second time**

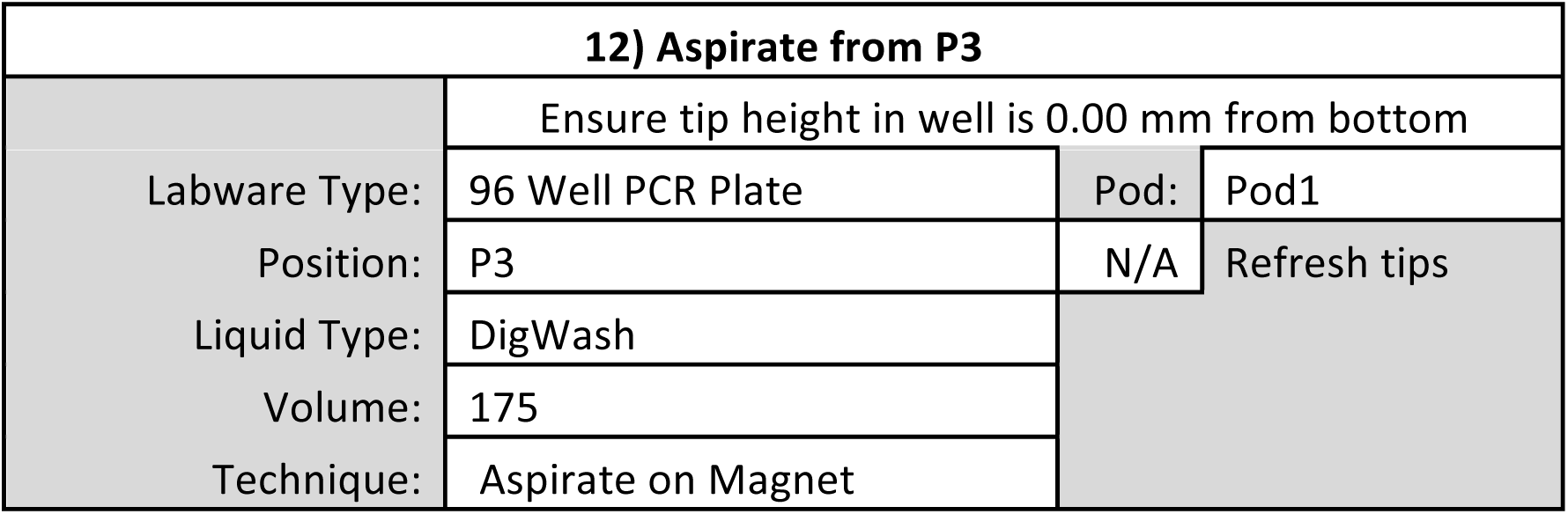

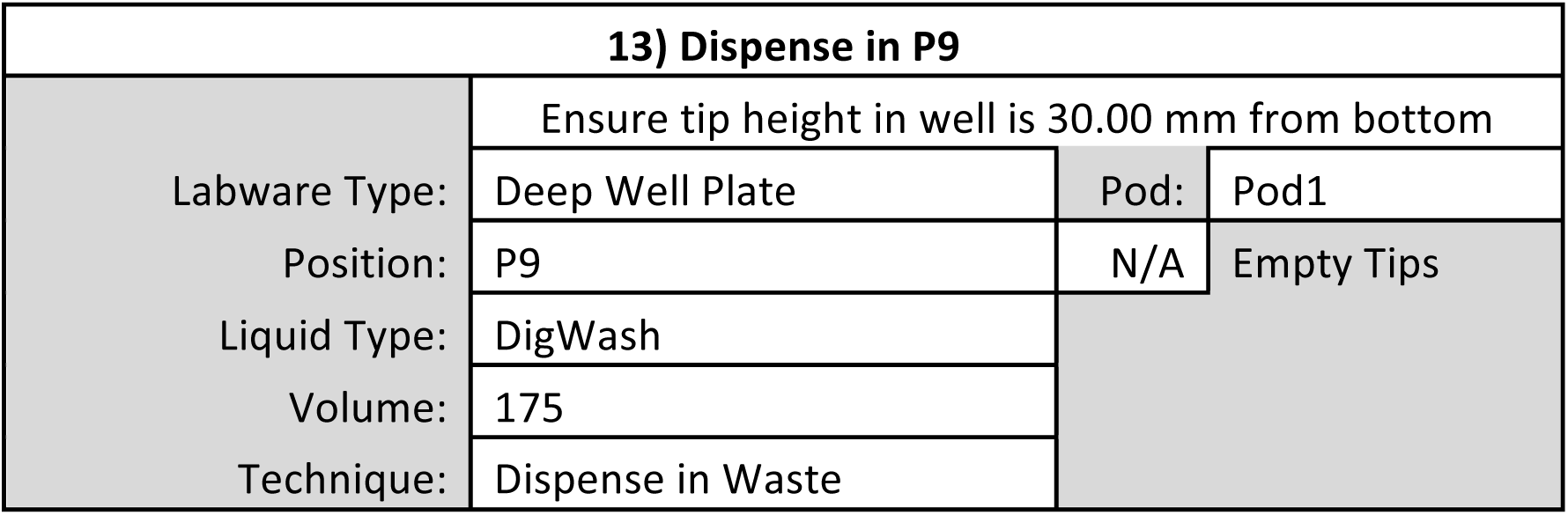

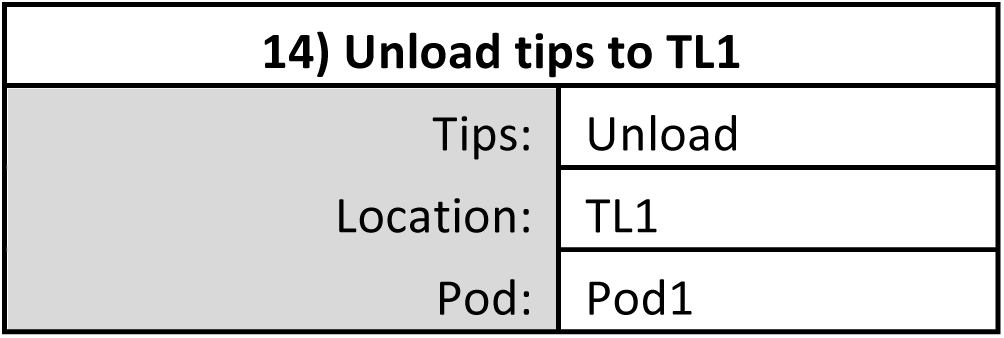

| 15) Move Labware from P3 to P10 |  |  |
| --- | --- | --- |
| Using pod: | Pod1 |  |
| Move labware from: | P3 | to P10 |
| Move the topmost: | 1 | piece of labware from the stack |

| 16) Load tips from TL1 |  |
| --- | --- |
| Tips: | Load |
| Location: | TL1 |
| Pod: | Pod1 |

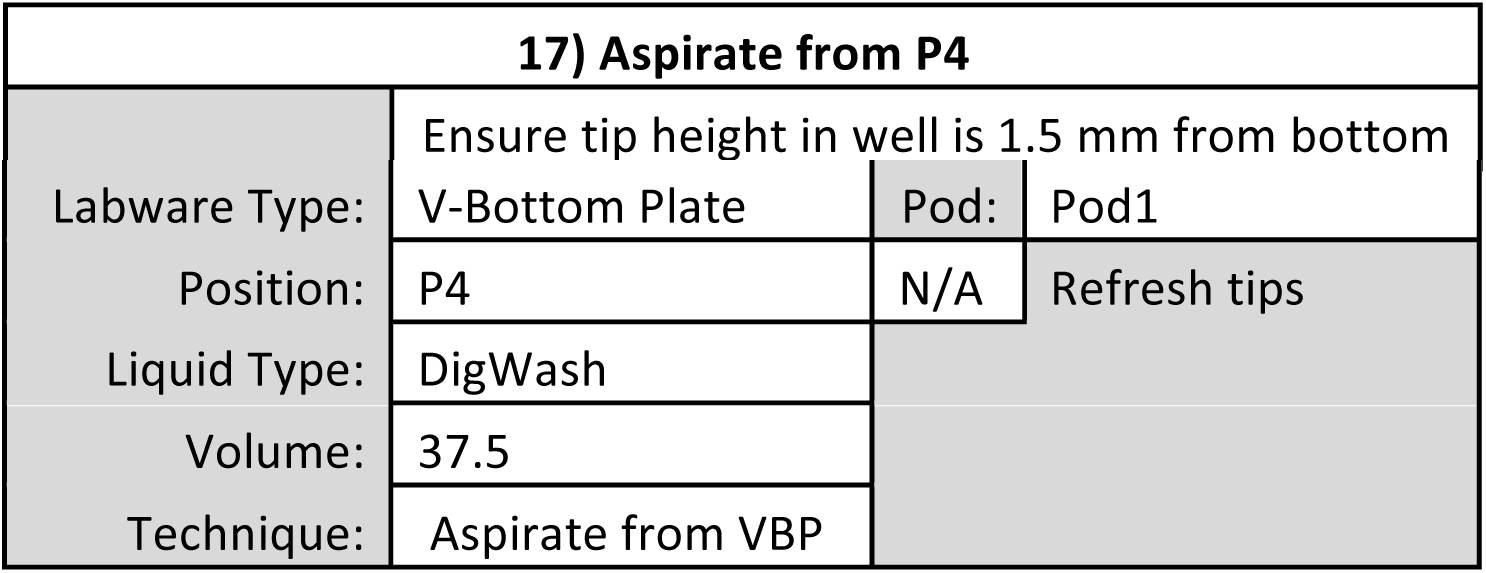

| 17) Aspirate from P4 |  |
| --- | --- |
|  | Ensure tip height in well is 1.5 mm from bottom |

|  |  |  |  |
| --- | --- | --- | --- |
| Labware Type: | V-Bottom Plate | Pod: | Pod1 |
| Position: | P4 | N/A | Refresh tips |
| Liquid Type: | DigWash |  |  |
| Volume: | 37.5 |  |  |
| Technique: | Aspirate from VBP |  |  |

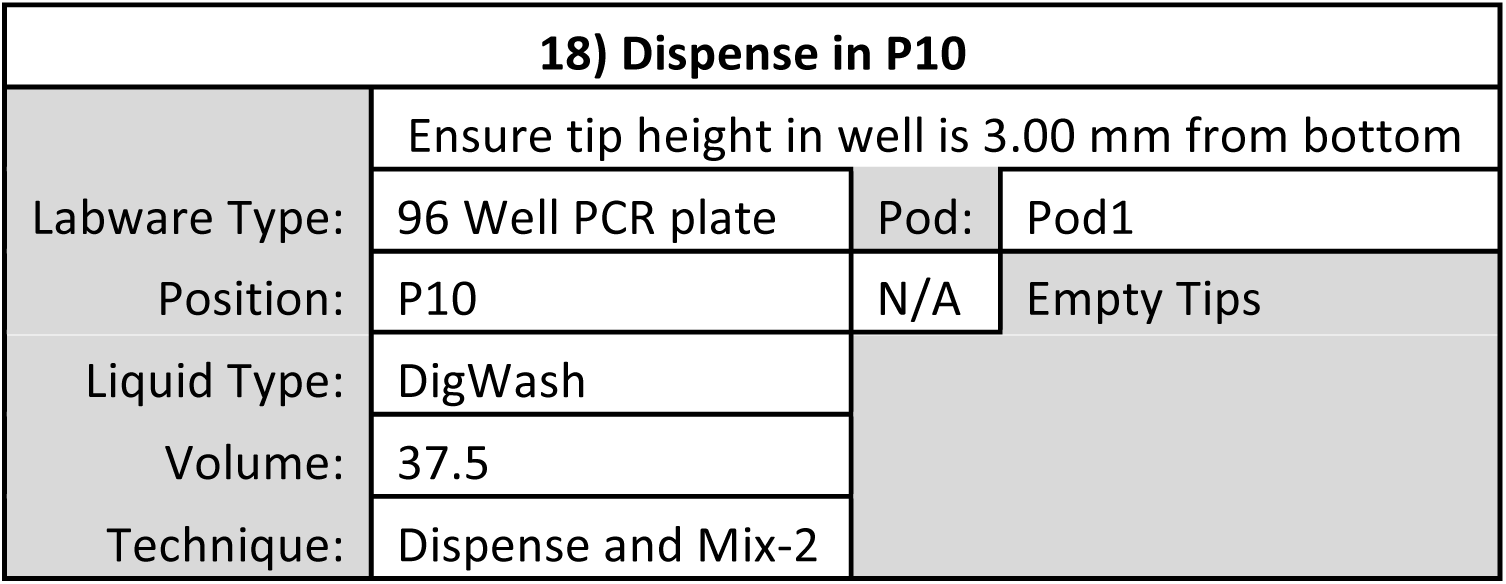

| 18) Dispense in P10 |  |  |  |
| --- | --- | --- | --- |
|  | Ensure tip height in well is 3.00 mm from bottom |  |  |
| Labware Type: | 96 Well PCR plate | Pod: | Pod1 |
| Position: | P10 | N/A | Empty Tips |
| Liquid Type: | DigWash |  |  |
| Volume: | 37.5 |  |  |
| Technique: | Dispense and Mix-2 |  |  |

| 19) Pause the whole system for 540 - 1800 s |  |  |  |
| --- | --- | --- | --- |
| Check: Pause | the whole system | for | 540 - 1800 s |

| 20) Aspirate from P7 |  |  |  |
| --- | --- | --- | --- |
|  | Ensure tip height in well is 1.5 mm from bottom |  |  |
| Labware Type: | V-Bottom Plate | Pod: | Pod1 |
| Position: | P7 | N/A | Refresh tips |
| Liquid Type: | DigWash |  |  |
| Volume: | 15 |  |  |
| Technique: | Aspirate from VBP |  |  |

| 21) Dispense in P10 |  |  |  |
| --- | --- | --- | --- |
|  | Ensure tip height in well is 3.00 mm from bottom |  |  |
| Labware Type: | 96 Well PCR Plate | Pod: | Pod1 |
| Position: | P10 | N/A | Empty Tips |
| Liquid Type: | DigWash |  |  |
| Volume: | 15 |  |  |
| Technique: | Dispense and Mix-3 |  |  |

| 22) Unload tips to TL1 |  |
| --- | --- |
| Tips: | Unload |
| Location: | TL1 |
| Pod: | Pod1 |

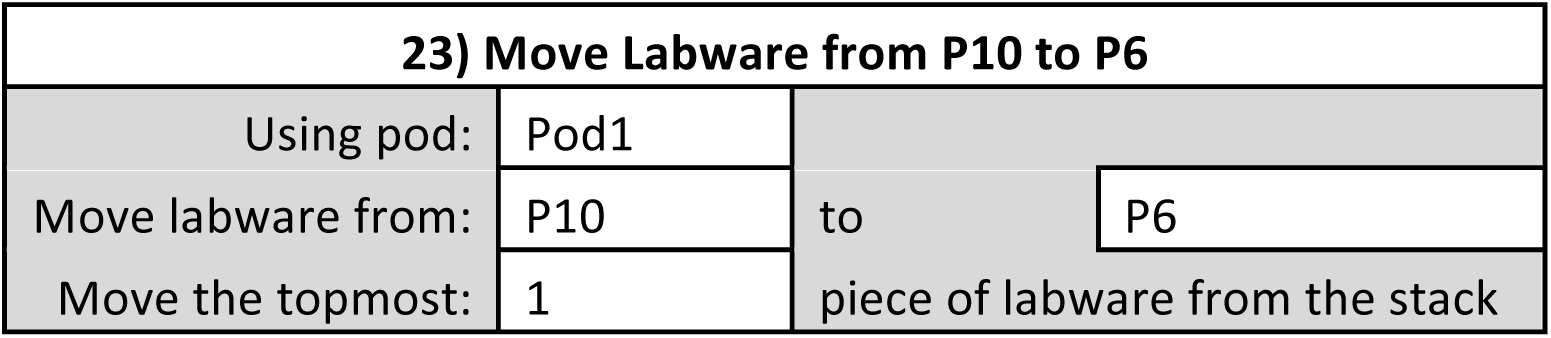

| 23) Move Labware from P10 to P6 |  |  |  |
| --- | --- | --- | --- |
| Using pod: | Pod1 |  |  |
| Move labware from: | P10 | to | P6 |
| Move the topmost: | 1 | piece of labware from the stack |  |

| 24) Load tips from TL1 |  |
| --- | --- |
| Tips: | Load |
| Location: | TL1 |
| Pod: | Pod1 |

| 25) Pause the whole system for 510 s |  |  |  |
| --- | --- | --- | --- |
| Check: Pause | the whole system | for | 510 s |

| 26) Mix in P6 |  |  |  |
| --- | --- | --- | --- |
|  |  | Ensure tip height in well is 3.00 mm from bottom |  |
| Labware Type: | 96 Well PCR plate | Pod | Pod1 |
| Position: | P6 | N/A | Refresh tips |
| Liquid Type: | DigWash | Mix: | 25 times |
| Volume: | 25 | uL |  |
| Technique: | Custom |  |  |
| Customize |  |  |  |
| Pipetting Template: | Default Template |  |  |
| Mix Tab |  |  |  |
| Move within the well at: | 25 | % speed |  |
| Aspirate at: | 3 | mm from the | Bottom at 25 ul/s |
| Dispense at: | 3 | mm from the | Bottom at 25 ul/s |
| N/A | Follow liquid level when aspirating or dispensing liquid |  |  |
| N/A | Touch tips on the sides of the wells |  |  |
| N/A | Aspirate a leading air gap prior to mix and blowout after mix is complete |  |  |

**27) Repeat Steps 25&26 two more times to release digested chromatin for ∼30min.**

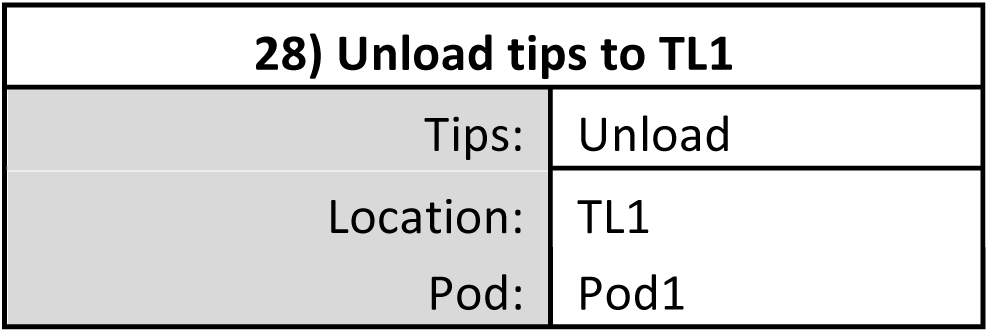

| 29) Move Labware from P6 to P3 |  |  |  |
| --- | --- | --- | --- |
| Using pod: | Pod1 |  |  |
| Move labware from: | P6 | to | P3 |
| Move the topmost: | 1 | piece of labware from the stack |  |

| 30) Pause the whole system for 180 s |  |  |  |
| --- | --- | --- | --- |
| Check: Pause | the whole system | for | 180 s |

| 31) Load tips from TL1 |  |
| --- | --- |
| Tips: | Load |
| Location: | TL1 |
| Pod: | Pod1 |

| 32) Aspirate from P3 |  |  |  |
| --- | --- | --- | --- |
|  | Ensure tip height in well is 0.0 mm from bottom |  |  |
| Labware Type: | 96 Well PCR plate | Pod: | Pod1 |
| Position: | P3 | N/A | Refresh tips |
| Liquid Type: | DigWash |  |  |
| Volume: | 37.5 |  |  |
| Technique: | Aspirate on Magnet |  |  |

| 33) Dispense in P8 |  |  |  |
| --- | --- | --- | --- |
|  | Ensure tip height in well is 3.00 mm from bottom |  |  |
| Labware Type: | 96 Well PCR Plate | Pod: | Pod1 |
| Position: | P8 | N/A | Empty Tips |
| Liquid Type: | DigWash |  |  |
| Volume: | 37.5 |  |  |
| Technique: | Dispense on Rack |  |  |

| 34) Unload tips to TL1 |  |
| --- | --- |
| Tips: | Unload |
| Location: | TL1 |
| Pod: | Pod1 |

| 35) Finish |
| --- |
| Check: Clear current instrument setup of all labware after the method completes |
| Check: Clear current device setup of all labware after the method completes |
| Check: Unload disposable tips from all pods after the method completes |
| Check: Clear all global variables after the method completes |

**Pre-PCR DNA Cleanup**

**1) Start**

**2) Instrument Setup**

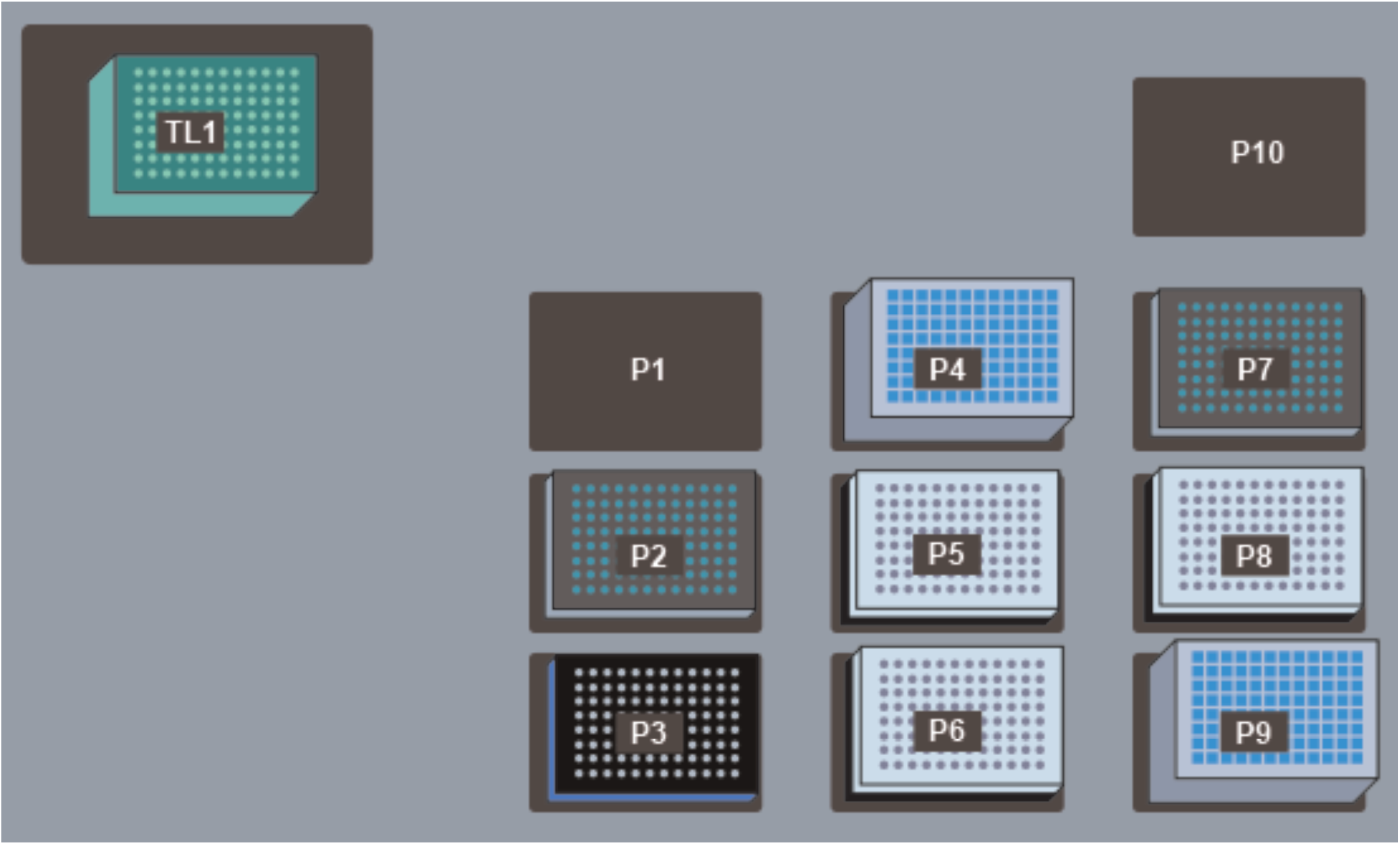

TL1: Fresh AP96 200 µL Tips (double click to increase the # of load times)

P2: V-Bottom Plate preloaded with 100 µL 10mM Tris-HCl pH 8

P3: ALPAQUA Magnet Plate

P4: Deep Well Plate preloaded with 1 mL 80% Ethanol

P5: 96 Well PCR Plate containing 110 µL of Adapter Ligated DNA stacked on a PCR Plate Rack

P6: 96 Well PCR Plate preloaded with 90 µL of Ampure Beads stacked on a PCR Plate Rack

P7: V-Bottom Plate preloaded with 100 µL HXP Mix

P8: 96 Well PCR Plate for accepting cleaned-up DNA stacked on a PCR Plate Rack

P9: Deep Well Plate for receiving liquid waste

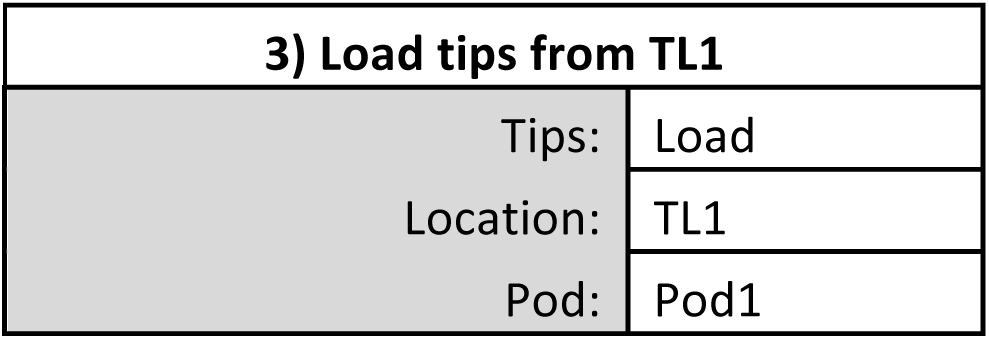

| 4) Aspirate from P5 |  |  |  |
| --- | --- | --- | --- |
|  | Ensure tip height in well is 0.50 mm from bottom |  |  |
| Labware Type: | 96 Well PCR plate | Pod: | Pod1 |
| Position: | P5 | N/A | Refresh tips |
| Liquid Type: | Ligase/PCR Buffer |  |  |
| Volume: | 110 |  |  |
| Technique: | Aspirate on Rack |  |  |

| 5) Dispense in P6 |  |  |  |
| --- | --- | --- | --- |
|  | Ensure tip height in well is 1.00 mm from bottom |  |  |
| Labware Type: | 96 Well PCR Plate | Pod: | Pod1 |
| Position: | P6 | N/A | Empty Tips |
| Liquid Type: | Ligase/PCR Buffer |  |  |
| Volume: | 110 |  |  |
| Technique: | Dispense and Mix-4 |  |  |

| 6) Unload tips to TL1 |  |
| --- | --- |
| Tips: | Unload |
| Location: | TL1 |
| Pod: | Pod1 |

| 7) Pause the whole system for 300 s |  |  |  |
| --- | --- | --- | --- |
| Check: Pause | the whole system | for | 300 s |

| 8) Move Labware from P6 to P3 |  |  |  |
| --- | --- | --- | --- |
| Using pod: | Pod1 |  |  |
| Move labware from: | P6 | to | P3 |
| Move the topmost: | 1 | piece of labware from the stack |  |

| 9) Pause the whole system for 300 s |  |  |  |
| --- | --- | --- | --- |
| Check: Pause | the whole system | for | 300 s |

| 10) Load tips from TL1 |  |  |
| --- | --- | --- |
|  | Tips: | Load |
|  | Location: | TL1 |
|  | Pod: | Pod1 |

| 11) Aspirate from P3 |  |  |  |
| --- | --- | --- | --- |
|  | Ensure tip height in well is 0.00 mm from bottom |  |  |
|  | Labware Type: | 96 Well PCR Plate | Pod: Pod1 |
|  | Position: | P3 | N/A Refresh tips |
|  | Liquid Type: | Ampure Wash |  |
|  | Volume: | 105 |  |
|  | Technique: | Aspirate on Magnet |  |

| 12) Dispense in P9 |  |  |  |
| --- | --- | --- | --- |
|  | Ensure tip height in well is 30.00 mm from bottom |  |  |
|  | Labware Type: | Deep Well Plate | Pod: Pod1 |
|  | Position: | P9 | N/A Empty Tips |
|  | Liquid Type: | Ampure Wash |  |
|  | Volume: | 105 |  |
|  | Technique: | Dispense in Waste |  |

**13) Repeat Steps 11&12 to remove remaining Ampure Buffer.**

| 14) Aspirate from P4 |  |  |  |
| --- | --- | --- | --- |
|  | Ensure tip height in well is 2.00 mm from bottom |  |  |
|  | Labware Type: | Deep Well Plate | Pod: Pod1 |
|  | Position: | P4 | N/A Refresh tips |
|  | Liquid Type: | EtOH |  |
|  | Volume: | 150 |  |
|  | Technique: | Aspirate from DWP |  |

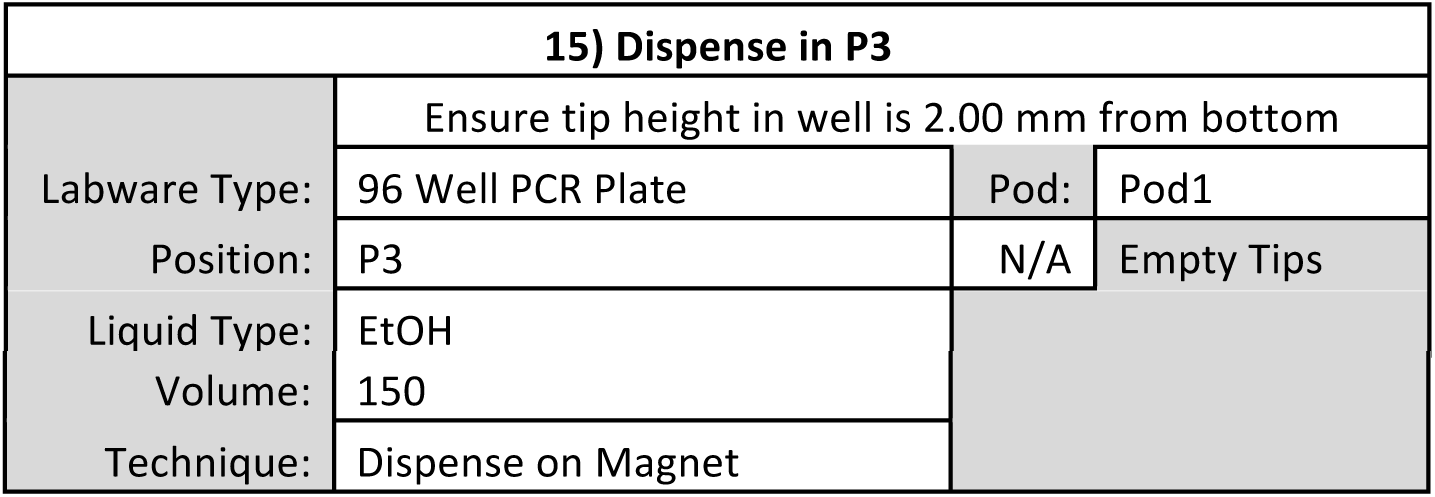

| 15) Dispense in P3 |  |  |  |
| --- | --- | --- | --- |
|  | Ensure tip height in well is 2.00 mm from bottom |  |  |
|  | Labware Type: | 96 Well PCR Plate | Pod: Pod1 |
|  | Position: | P3 | N/A Empty Tips |
|  | Liquid Type: | EtOH |  |

|  |  |
| --- | --- |
| Volume: | 150 |
| Technique: | Dispense on Magnet |

| 16) Pause the whole system for 30 s |  |  |  |
| --- | --- | --- | --- |
| Check: Pause | the whole system | for | 30 s |

| 17) Aspirate from P3 |  |  |  |
| --- | --- | --- | --- |
|  | Ensure tip height in well is 0.00 mm from bottom |  |  |
| Labware Type: | 96 Well PCR Plate | Pod: | Pod1 |
| Position: | P3 | N/A | Refresh tips |
| Liquid Type: | EtOH |  |  |
| Volume: | 155 |  |  |
| Technique: | Aspirate on Magnet |  |  |

| 18) Dispense in P9 |  |  |  |
| --- | --- | --- | --- |
|  | Ensure tip height in well is 30.00 mm from bottom |  |  |
| Labware Type: | Deep Well Plate | Pod: | Pod1 |
| Position: | P9 | N/A | Empty Tips |
| Liquid Type: | EtOH |  |  |
| Volume: | 155 |  |  |
| Technique: | Dispense in Waste |  |  |

**19) Repeat Steps 14-18 to wash bead-bound DNA a second time with Ethanol**

| 20) Pause the whole system for 600 s |  |  |  |
| --- | --- | --- | --- |
| Check: Pause | the whole system | for | 600 s |

| 21) Unload tips to TL1 |  |
| --- | --- |
| Tips: | Unload |
| Location: | TL1 |
| Pod: | Pod1 |

| 22) Move Labware from P3 to P6 |  |  |  |
| --- | --- | --- | --- |
| Using pod: | Pod1 |  |  |
| Move labware from: | P3 | to | P6 |
| Move the topmost: | 1 | piece of labware from the stack |  |

| 23) Load tips from TL1 |  |  |
| --- | --- | --- |
|  | Tips: | Load |
|  | Location: | TL1 |
|  | Pod: | Pod1 |

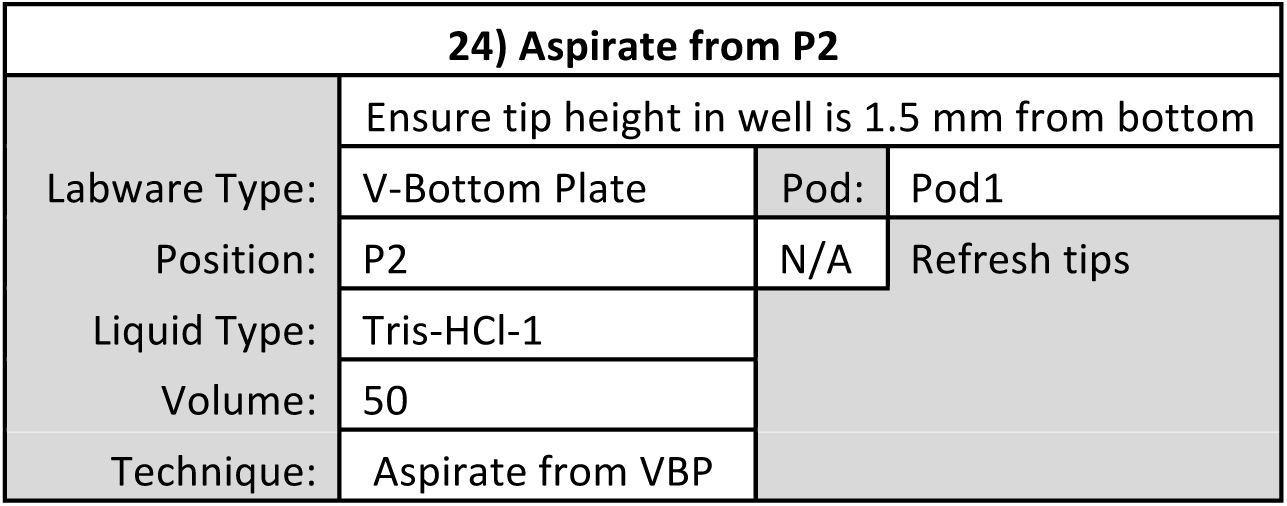

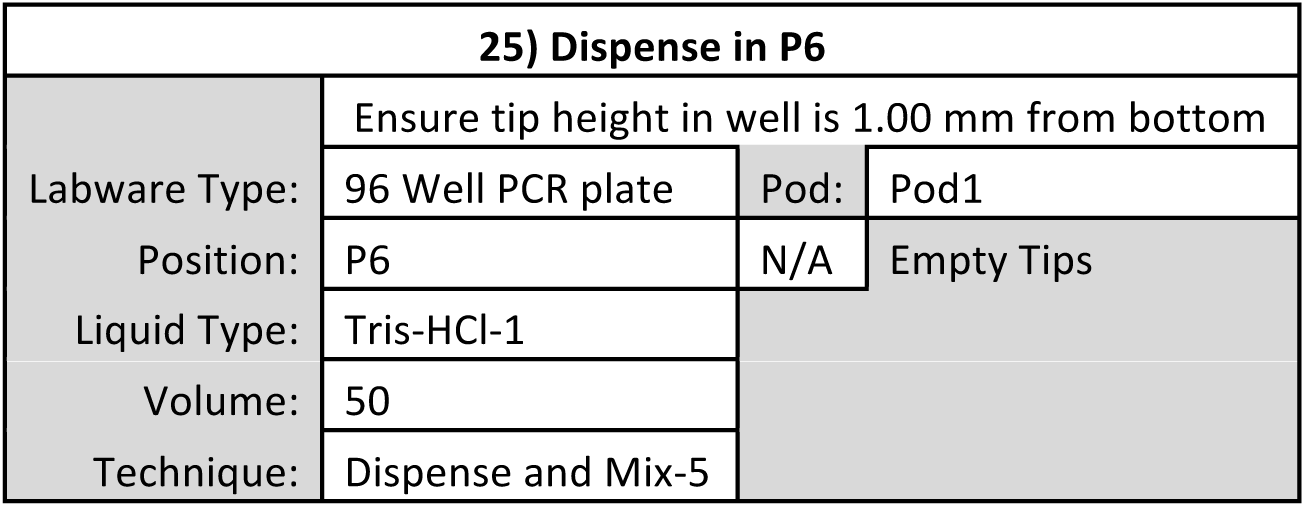

| 26) Pause the whole system for 300 s |  |  |  |
| --- | --- | --- | --- |
| Check: Pause | the whole system | for | 300 s |

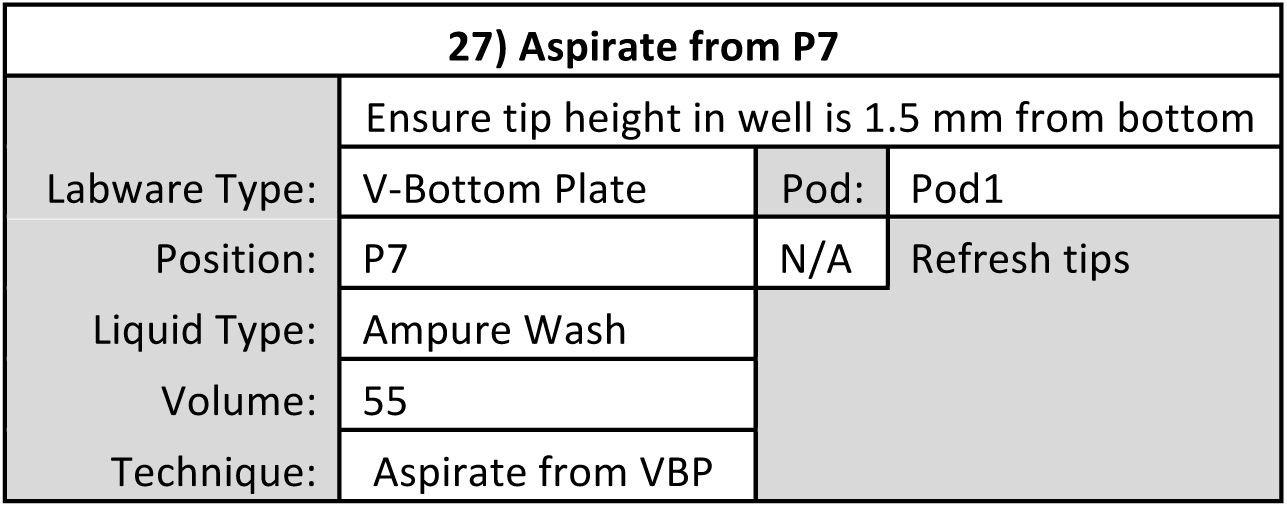

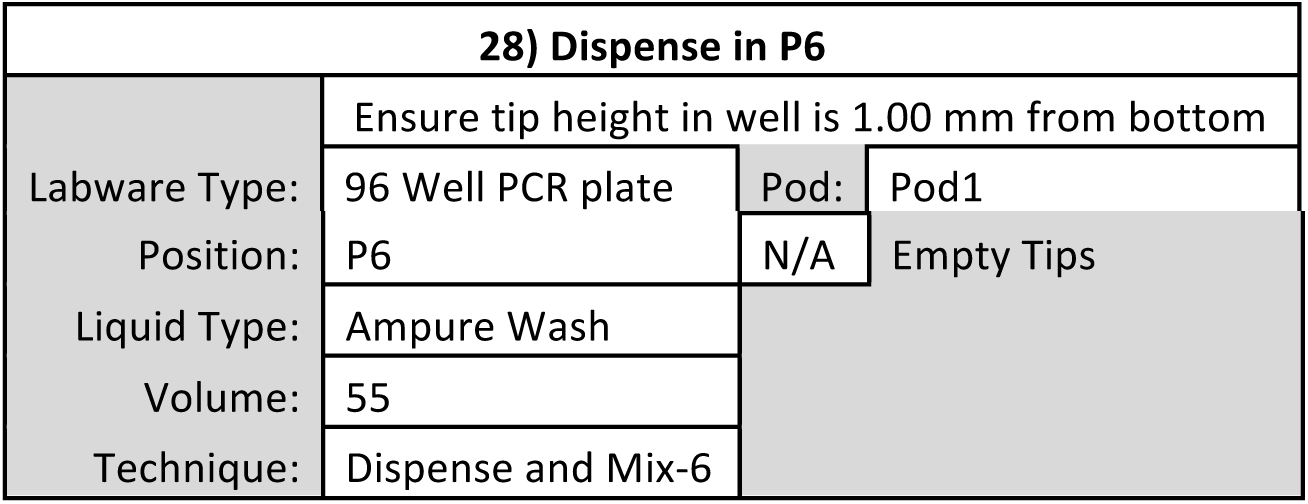

| 29) Unload tips to TL1 |  |
| --- | --- |
| Tips: | Unload |
| Location: | TL1 |
| Pod: | Pod1 |

| 30) Pause the whole system for 300 s |  |  |  |
| --- | --- | --- | --- |
| Check: Pause | the whole system | for | 300 s |

| 31) Move Labware from P6 to P3 |  |  |
| --- | --- | --- |
| Using pod: | Pod1 |  |
| Move labware from: | P6 | to P3 |
| Move the topmost: | 1 | piece of labware from the stack |

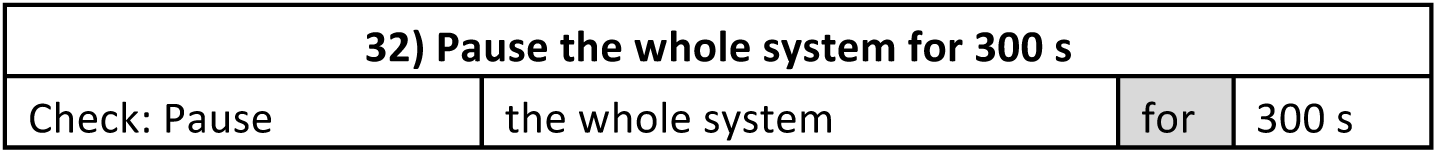

| 33) Load tips from TL1 |  |
| --- | --- |
| Tips: | Load |
| Location: | TL1 |
| Pod: | Pod1 |

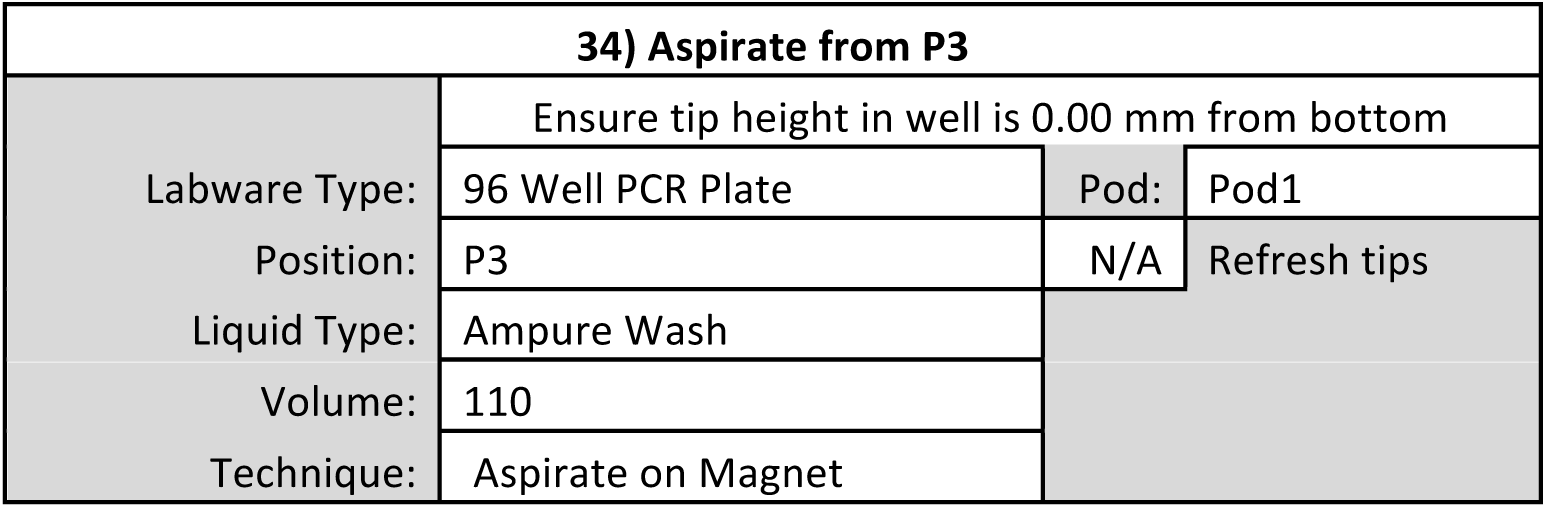

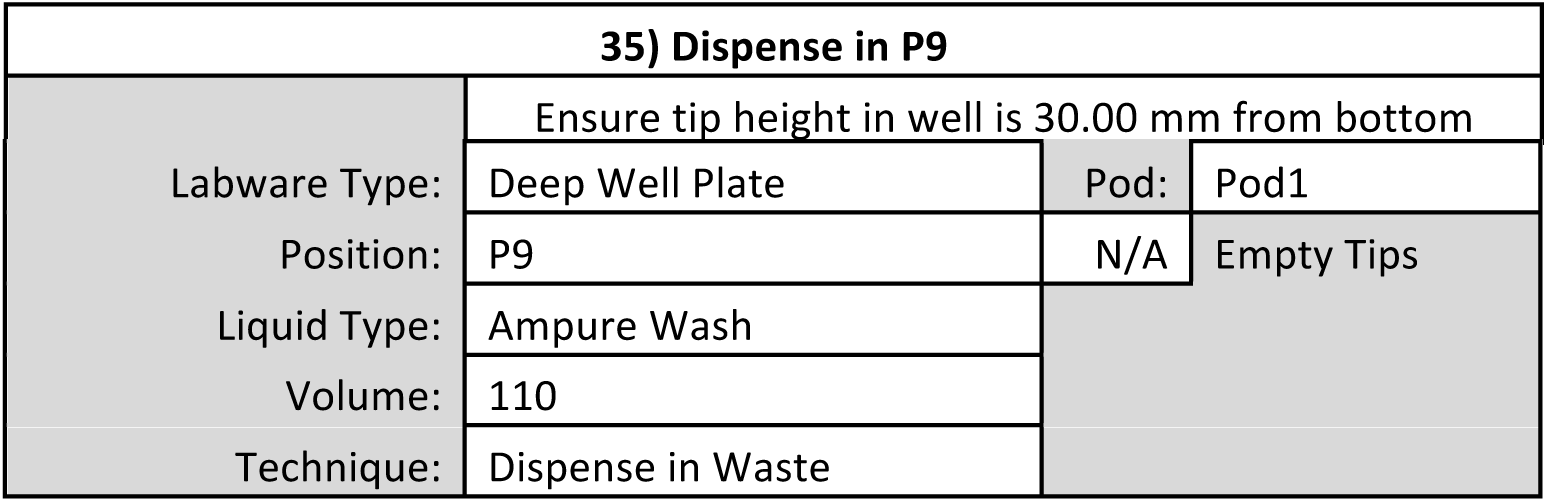

| 35) Dispense in P9 |  |
| --- | --- |
|  | Ensure tip height in well is 30.00 mm from bottom |

|  |  |  |  |
| --- | --- | --- | --- |
| Labware Type: | Deep Well Plate | Pod: | Pod1 |
| Position: | P9 | N/A | Empty Tips |
| Liquid Type: | Ampure Wash |  |  |
| Volume: | 110 |  |  |
| Technique: | Dispense in Waste |  |  |

**36) Perform Steps 14-21 again to wash bead-bound DNA twice with Ethanol and then allow to air dry.**

| 37) Move Labware from P3 to P6 |  |  |  |
| --- | --- | --- | --- |
| Using pod: | Pod1 |  |  |
| Move labware from: | P3 | to | P6 |
| Move the topmost: | 1 | piece of labware from the stack |  |

| 38) Load tips from TL1 |  |
| --- | --- |
| Tips: | Load |
| Location: | TL1 |
| Pod: | Pod1 |

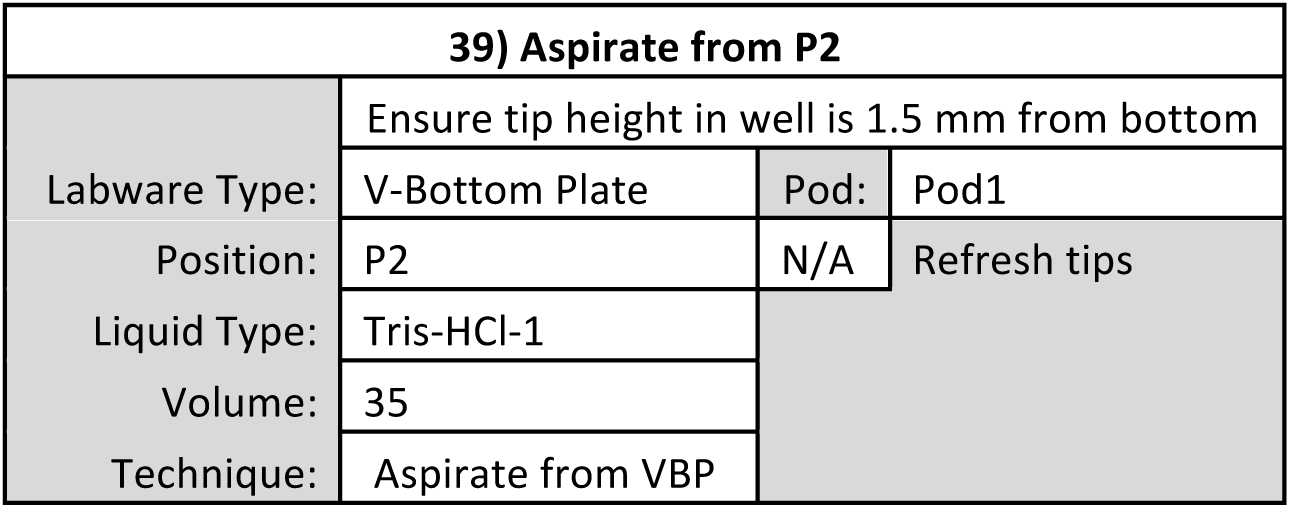

| 40) Dispense in P6 |  |  |  |
| --- | --- | --- | --- |
|  | Ensure tip height in well is 1.00 mm from bottom |  |  |
| Labware Type: | 96 Well PCR plate | Pod: | Pod1 |
| Position: | P6 | N/A | Empty Tips |
| Liquid Type: | Tris-HCl-1 |  |  |
| Volume: | 35 |  |  |
| Technique: | Dispense and Mix-5 |  |  |

| 41) Unload tips to TL1 |  |
| --- | --- |
| Tips: | Unload |
| Location: | TL1 |
| Pod: | Pod1 |

| 42) Pause the whole system for 300 s |  |  |  |
| --- | --- | --- | --- |
| Check: Pause | the whole system | for | 300 s |

| 43) Move Labware from P6 to P3 |  |  |  |
| --- | --- | --- | --- |
| Using pod: | Pod1 |  |  |
| Move labware from: | P6 | to | P3 |
| Move the topmost: | 1 | piece of labware from the stack |  |

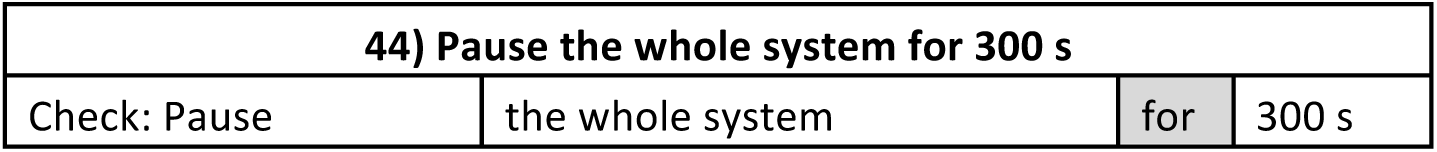

| 45) Load tips from TL1 |  |
| --- | --- |
| Tips: | Load |
| Location: | TL1 |
| Pod: | Pod1 |

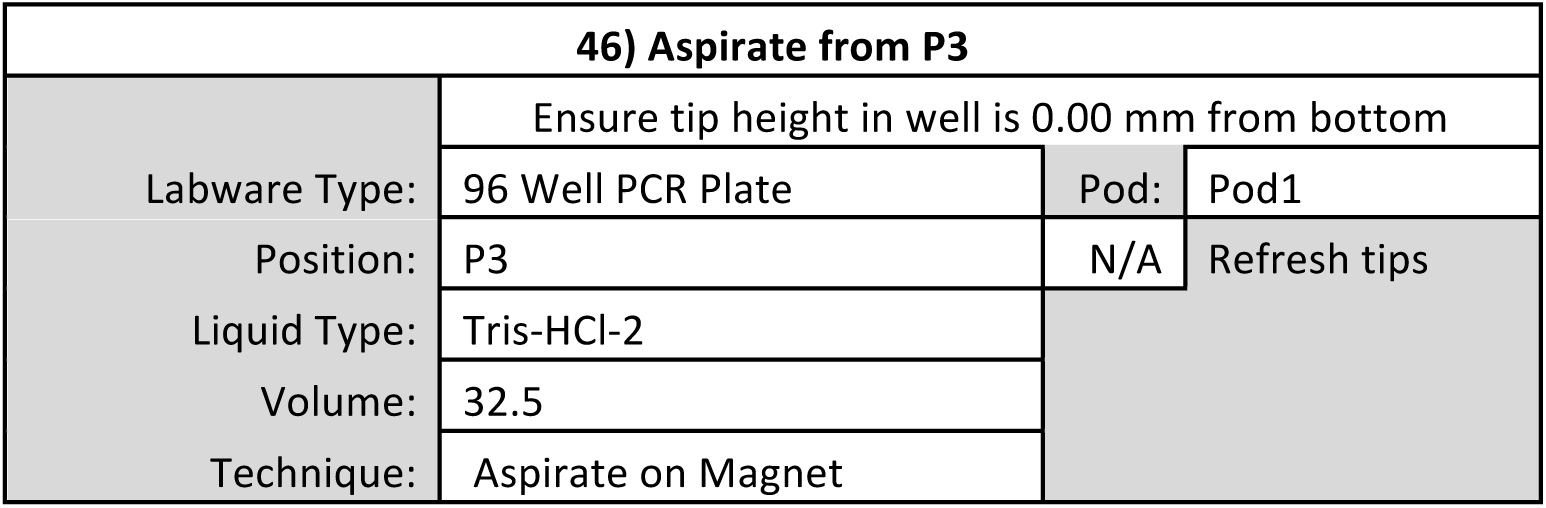

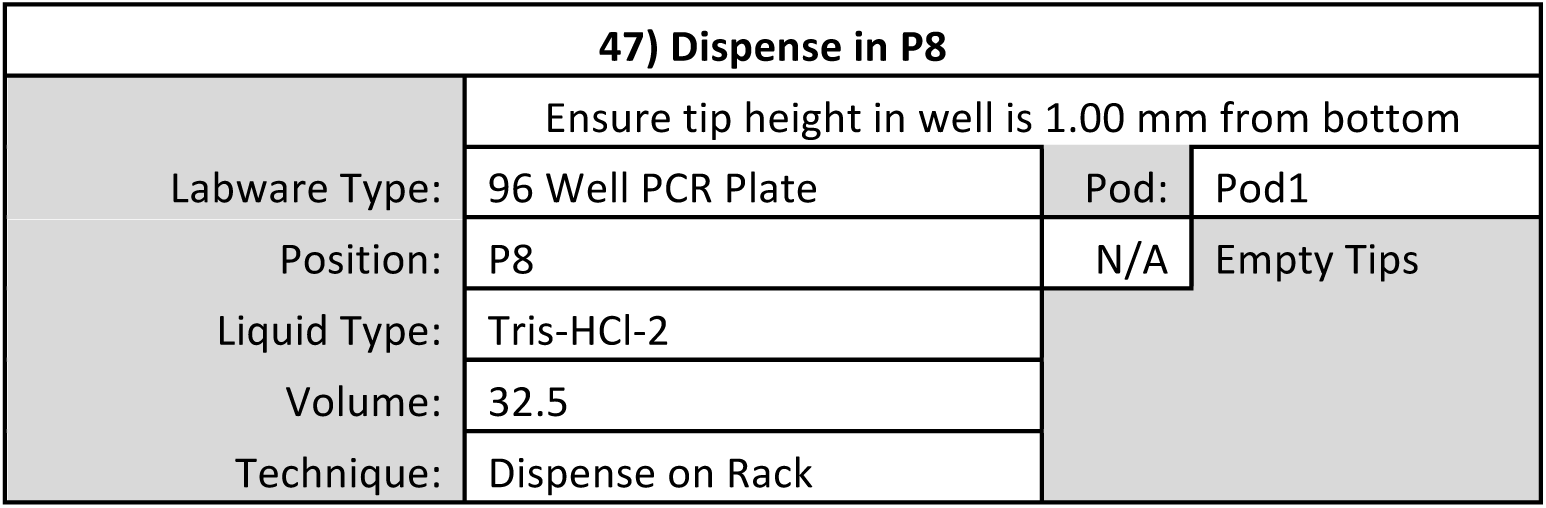

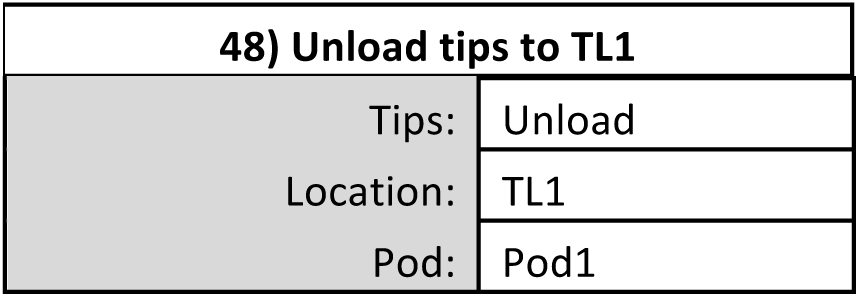

| 48) Unload tips to TL1 |
| --- |

|  |  |
| --- | --- |
| Tips: | Unload |
| Location: | TL1 |
| Pod: | Pod1 |

| 49) Finish |
| --- |
| Check: Clear current instrument setup of all labware after the method completes |
| Check: Clear current device setup of all labware after the method completes |
| Check: Unload disposable tips from all pods after the method completes |
| Check: Clear all global variables after the method completes |

**Post-PCR DNA Cleanup**

**Start**

**Instrument Setup**

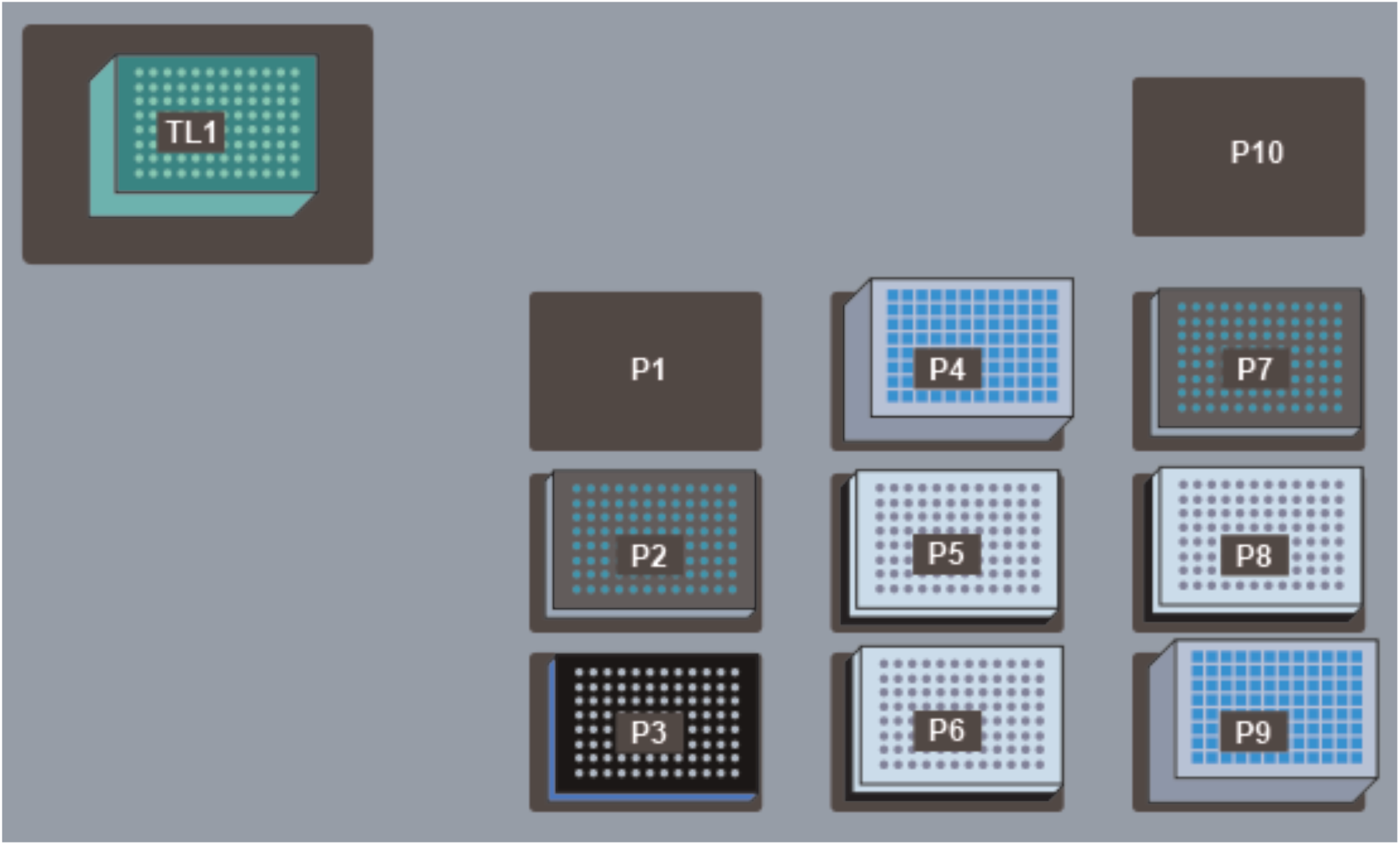

TL1: Fresh AP96 200 µL Tips (double click to increase the # of load times)

P2: V-Bottom Plate preloaded with 100 µL 10mM Tris-HCl pH 8

P3: ALPAQUA Magnet Plate

P4: Deep Well Plate preloaded with 1 mL 80% Ethanol

P5: 96 Well PCR Plate containing 50 µL of PCR product stacked on a PCR Plate Rack

P6: 96 Well PCR Plate preloaded with 55 µL of Ampure Beads stacked on a PCR Plate Rack

P7: V-Bottom Plate preloaded with 100 µL HXP Mix

P8: 96 Well PCR Plate for accepting cleaned-up DNA stacked on a PCR Plate Rack

P9: Deep Well Plate for receiving liquid waste

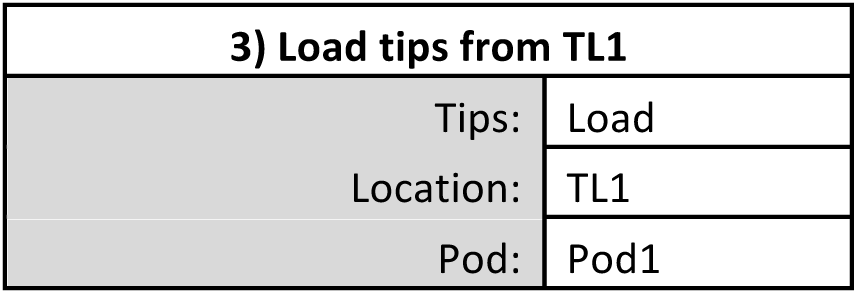

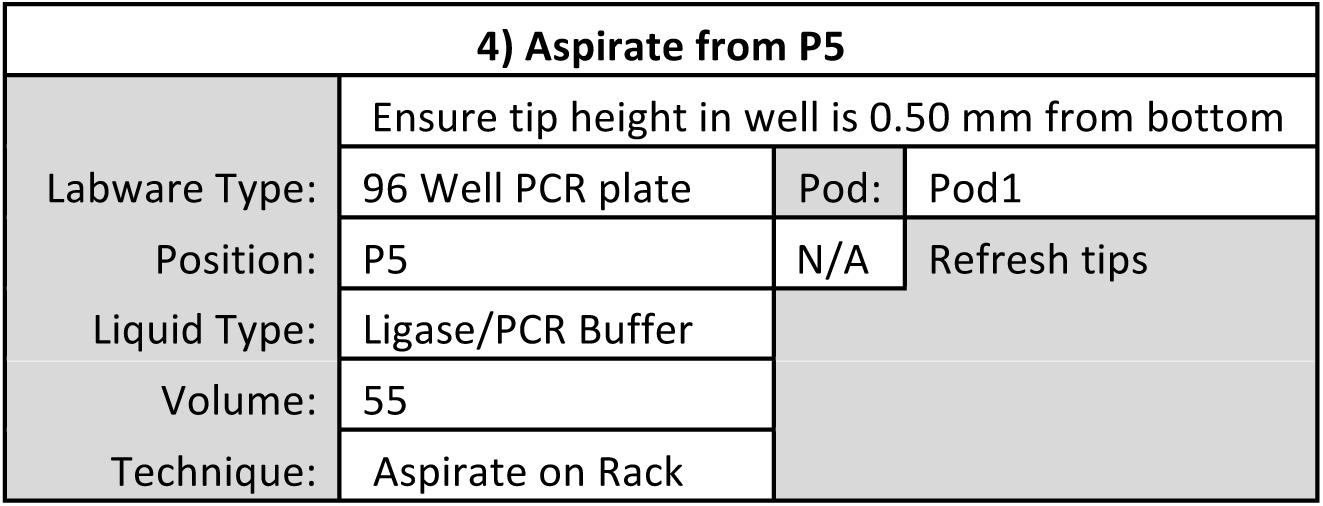

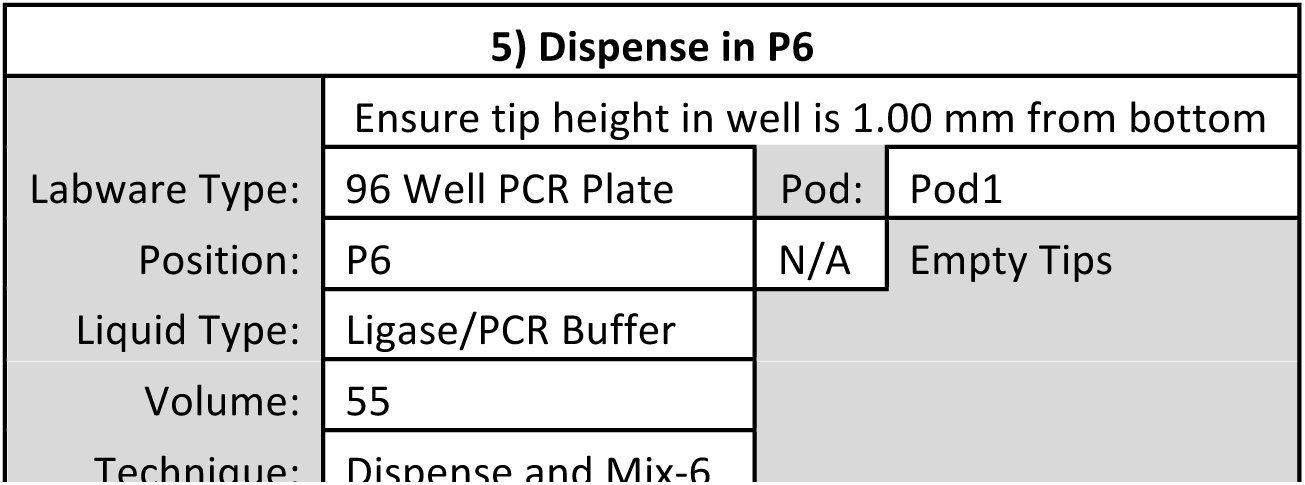

| 6) Unload tips to TL1 |  |  |
| --- | --- | --- |
|  | Tips: | Unload |
|  | Location: | TL1 |
|  | Pod: | Pod1 |

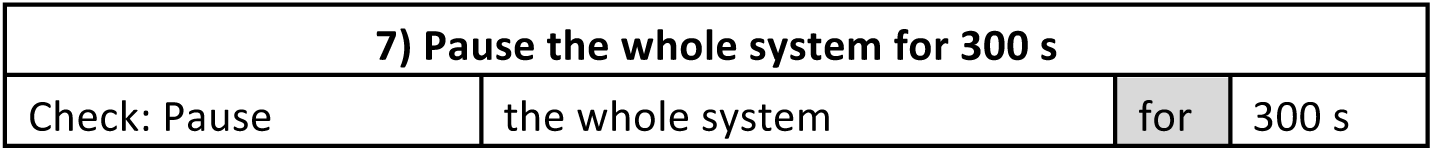

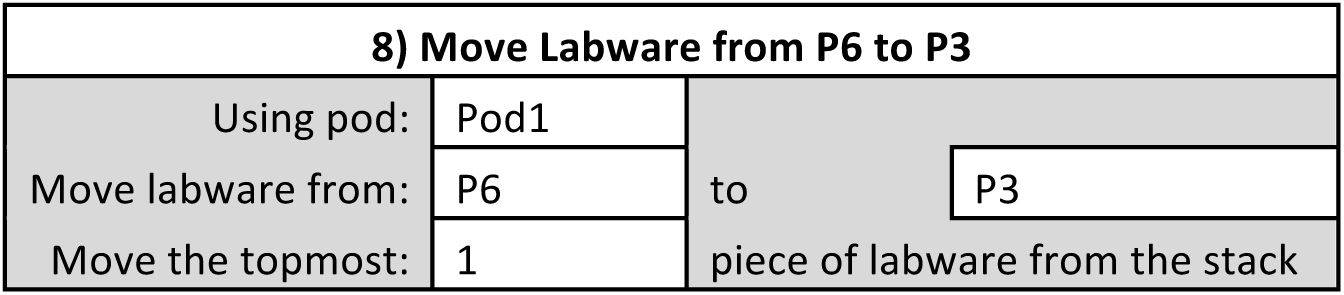

| 9) Pause the whole system for 300 s |  |  |  |
| --- | --- | --- | --- |
| Check: Pause | the whole system | for | 300 s |

| 10) Load tips from TL1 |  |
| --- | --- |
| Tips: | Load |
| Location: | TL1 |
| Pod: | Pod1 |

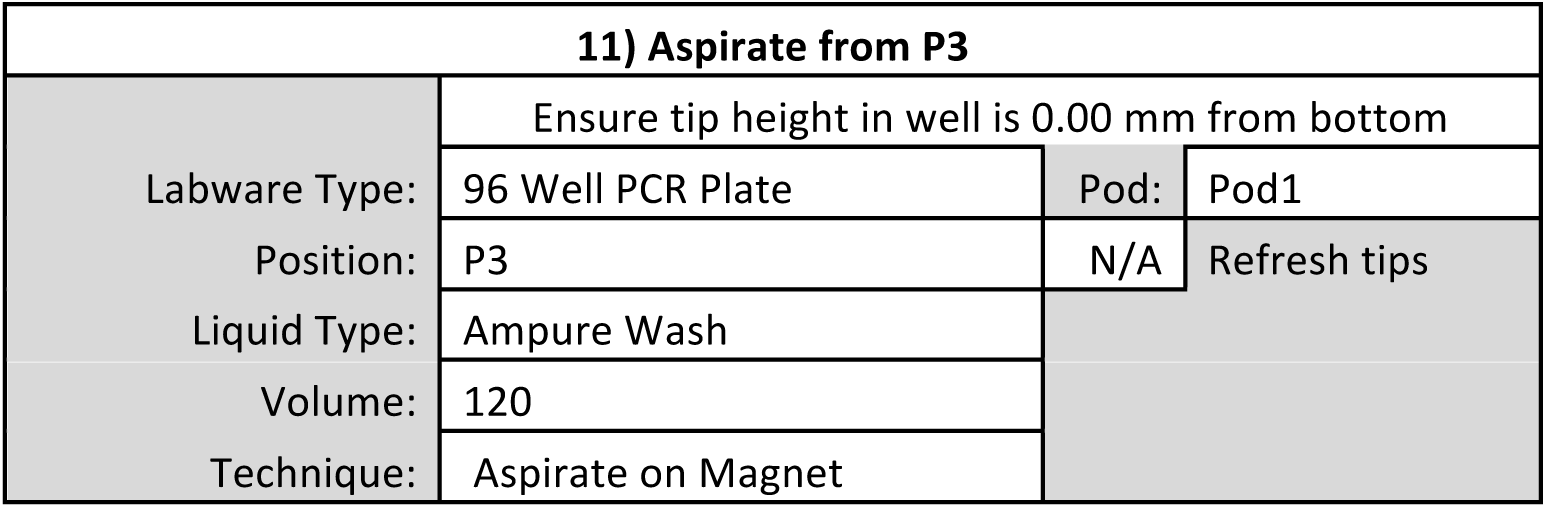

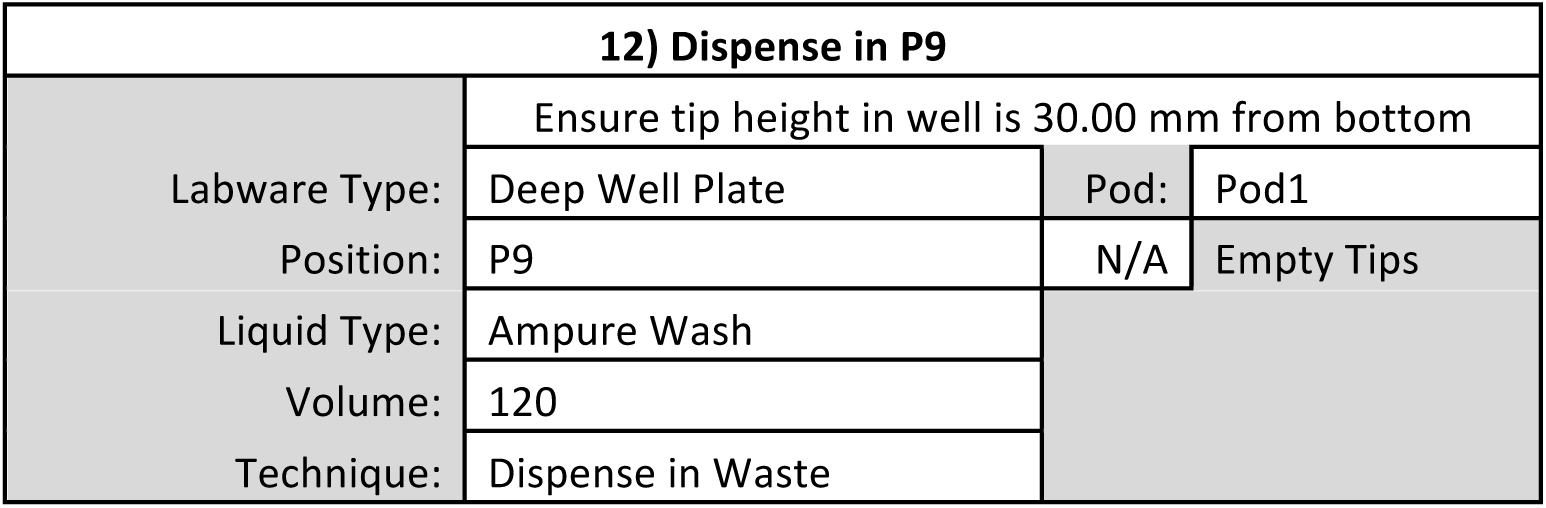

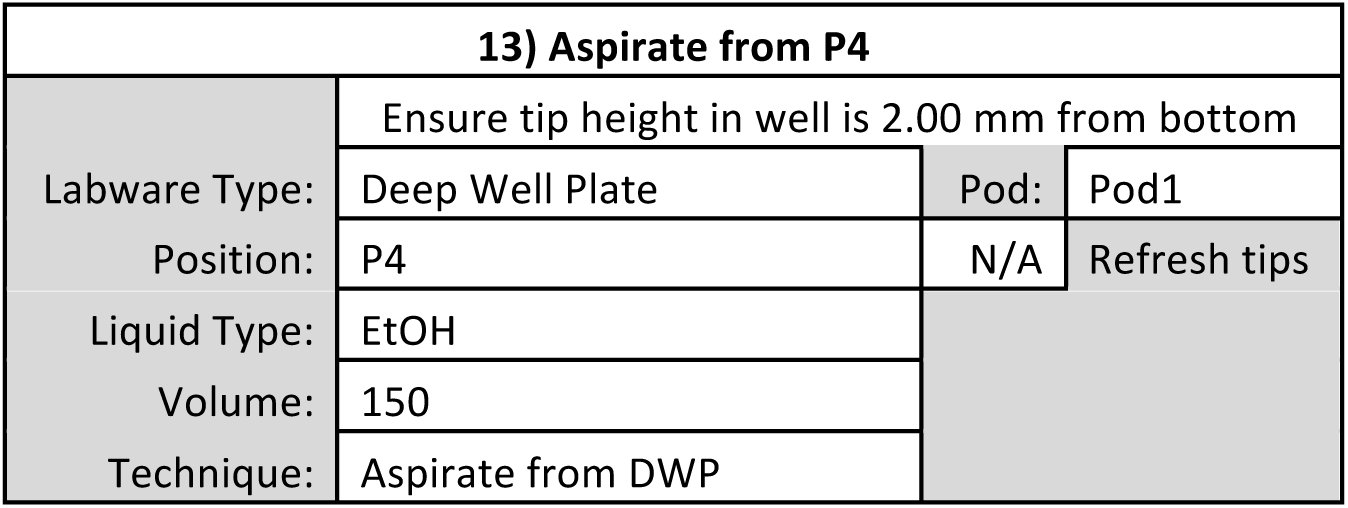

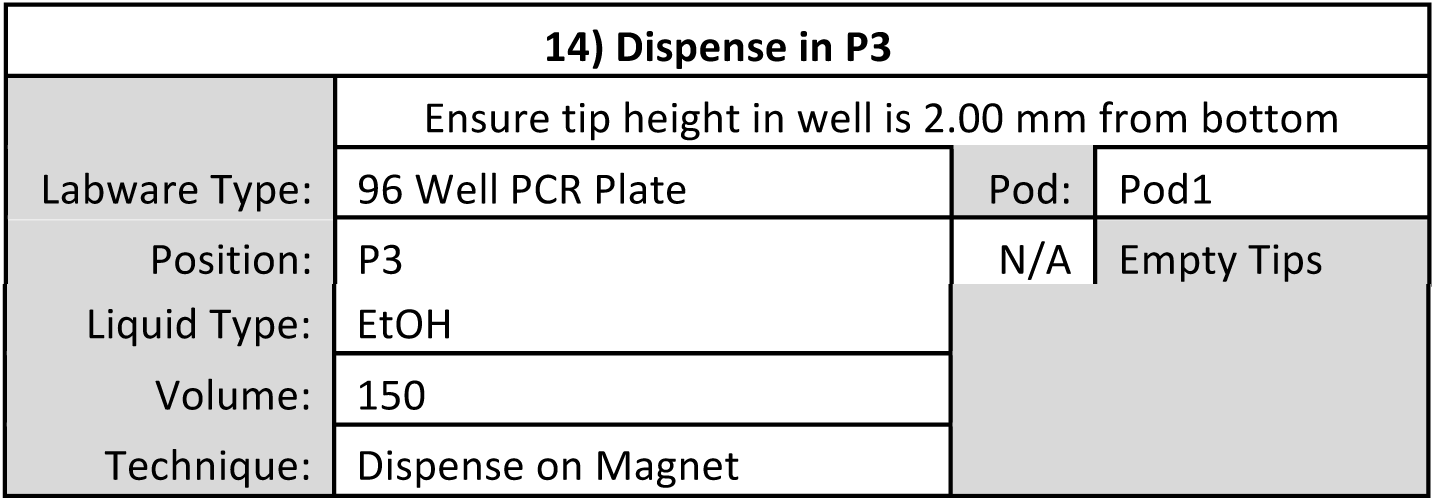

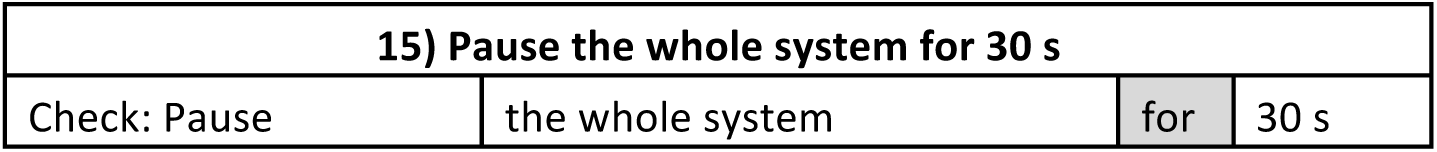

| 15) Pause the whole system for 30 s |  |  |  |
| --- | --- | --- | --- |
| Check: Pause | the whole system | for | 30 s |

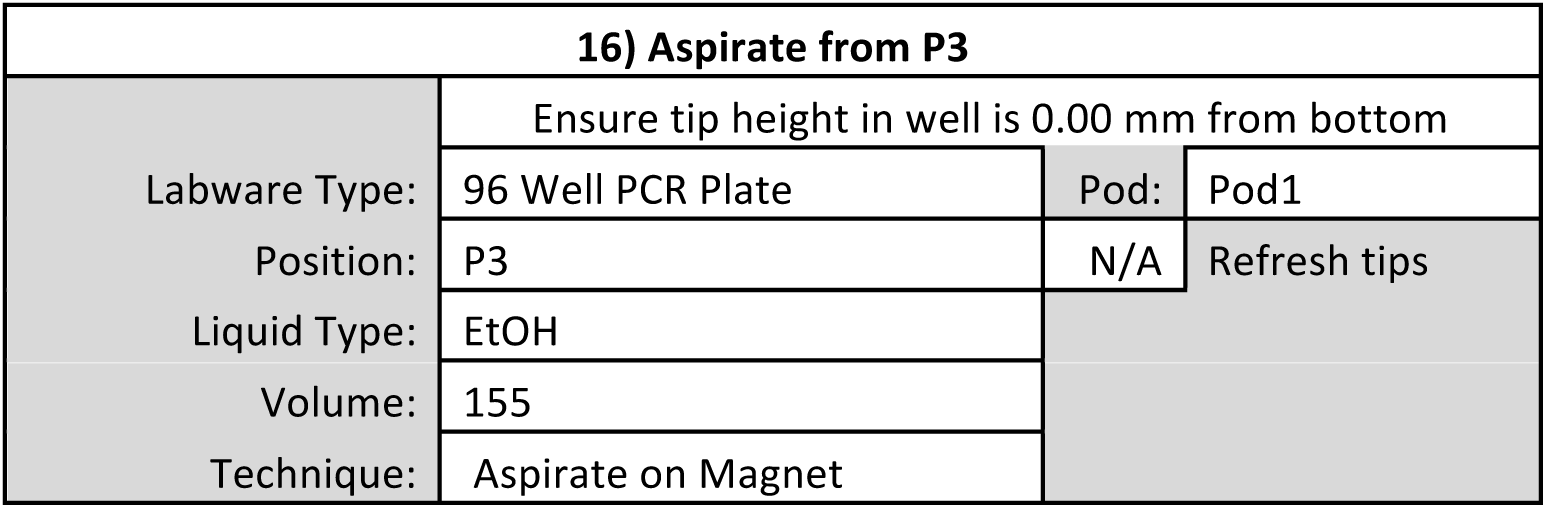

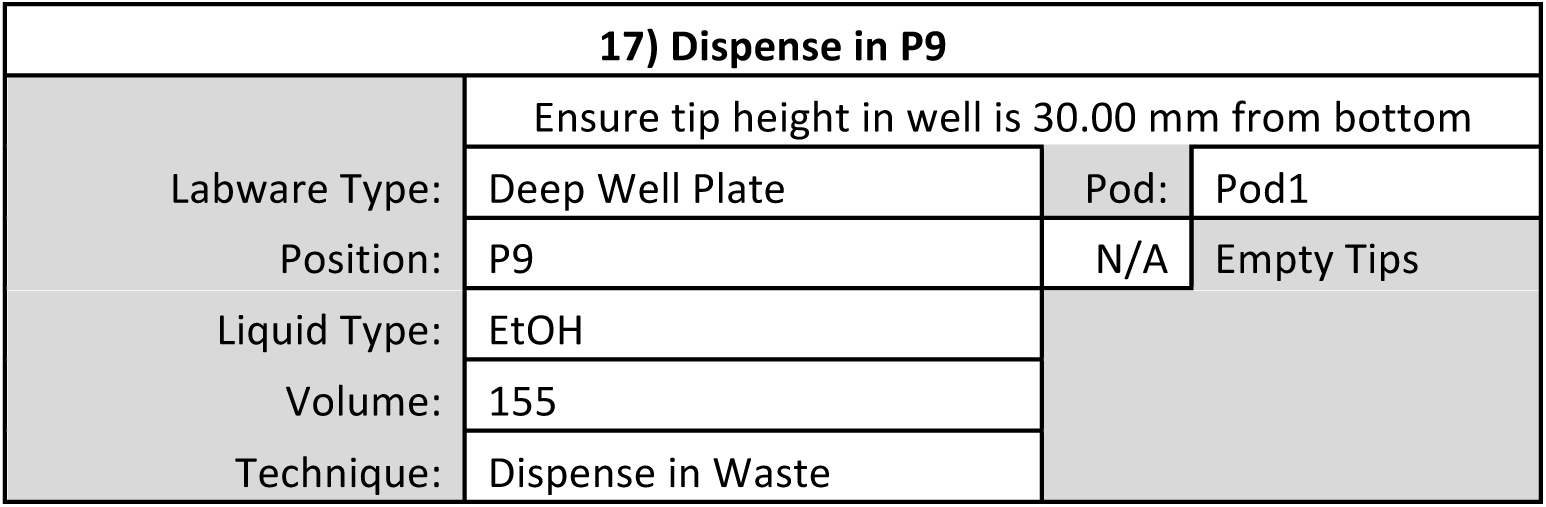

**18) Repeat Steps 13-17 to wash bead-bound DNA a second time with Ethanol**

| 19) Pause the whole system for 600 s |  |  |  |
| --- | --- | --- | --- |
| Check: Pause | the whole system | for | 600 s |

| 20) Unload tips to TL1 |  |
| --- | --- |
| Tips: | Unload |
| Location: | TL1 |
| Pod: | Pod1 |

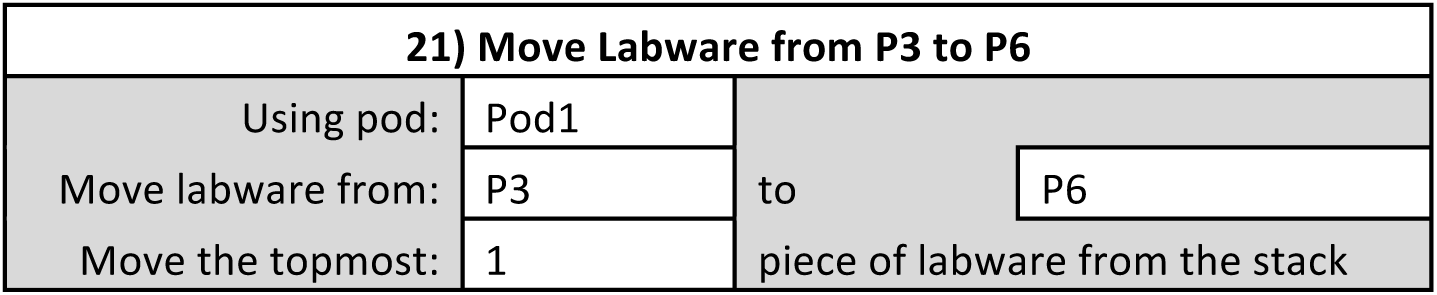

| 22) Load tips from TL1 |  |  |
| --- | --- | --- |
|  | Tips: | Load |
|  | Location: | TL1 |
|  | Pod: | Pod1 |

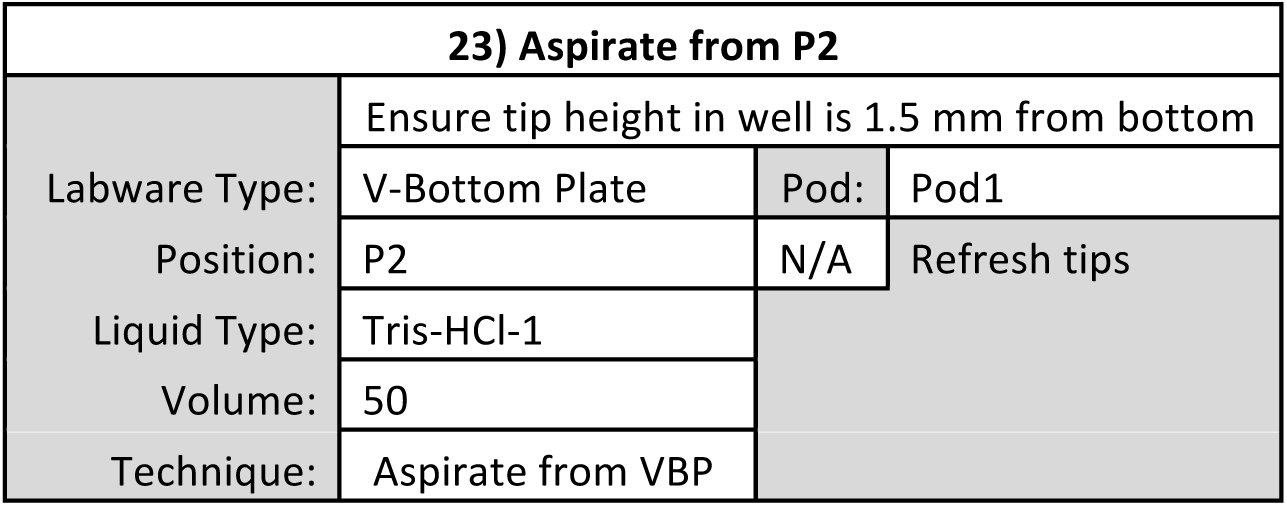

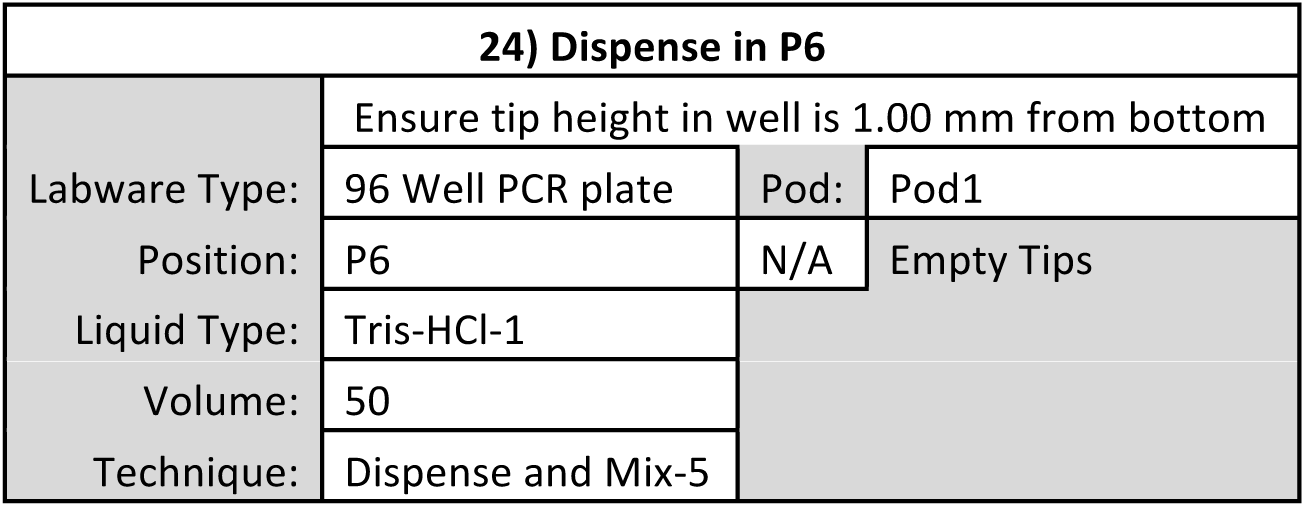

| 25) Pause the whole system for 300 s |  |  |  |
| --- | --- | --- | --- |
| Check: Pause | the whole system | for | 300 s |

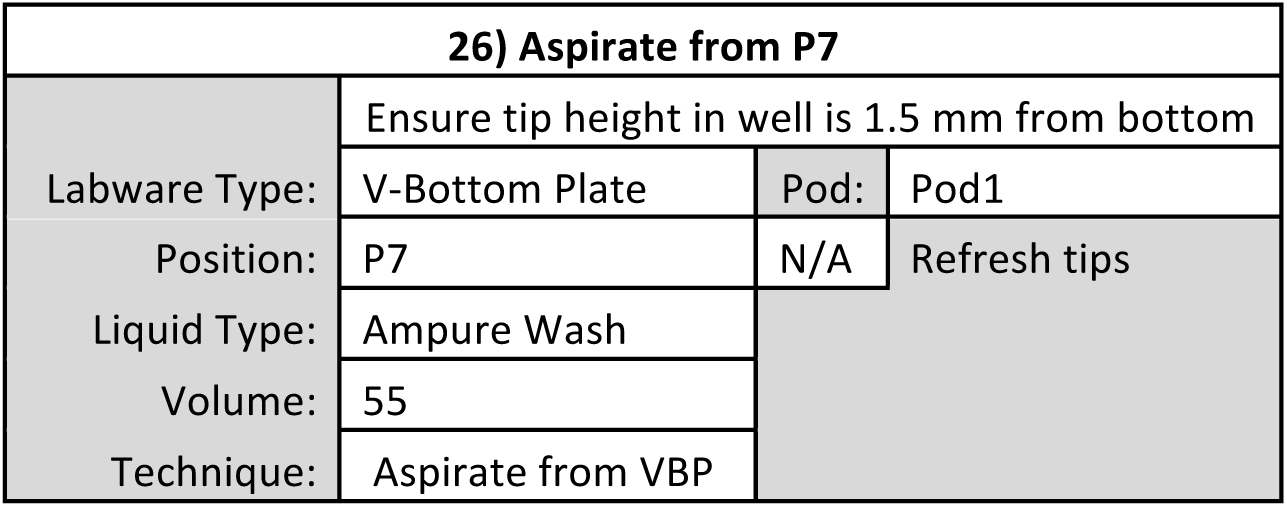

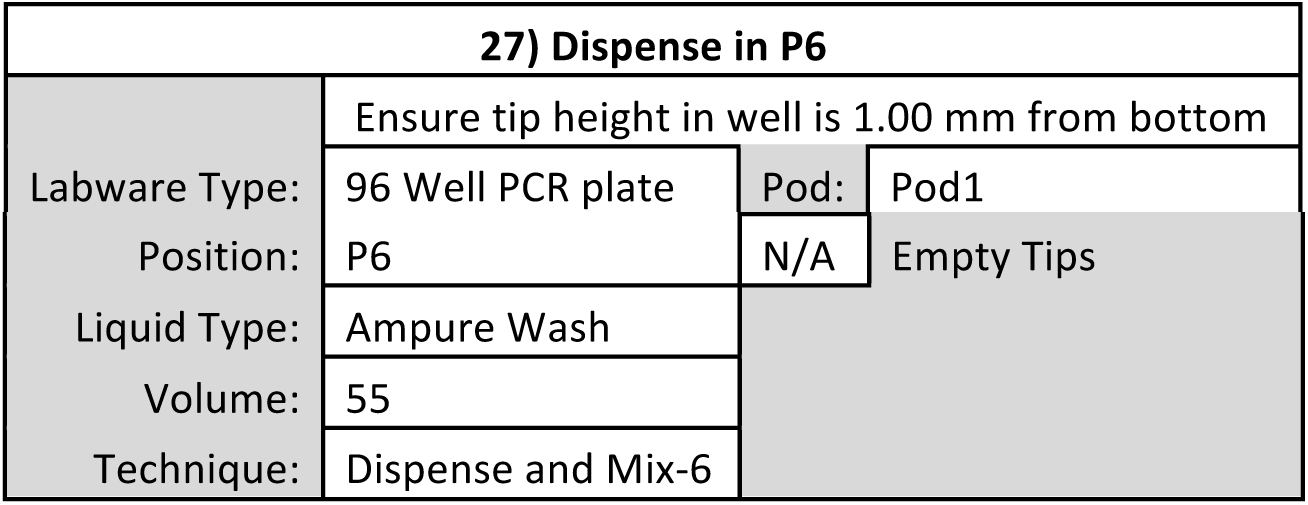

| 28) Unload tips to TL1 |  |
| --- | --- |
| Tips: | Unload |
| Location: | TL1 |
| Pod: | Pod1 |

| 29) Pause the whole system for 300 s |  |  |  |
| --- | --- | --- | --- |
| Check: Pause | the whole system | for | 300 s |

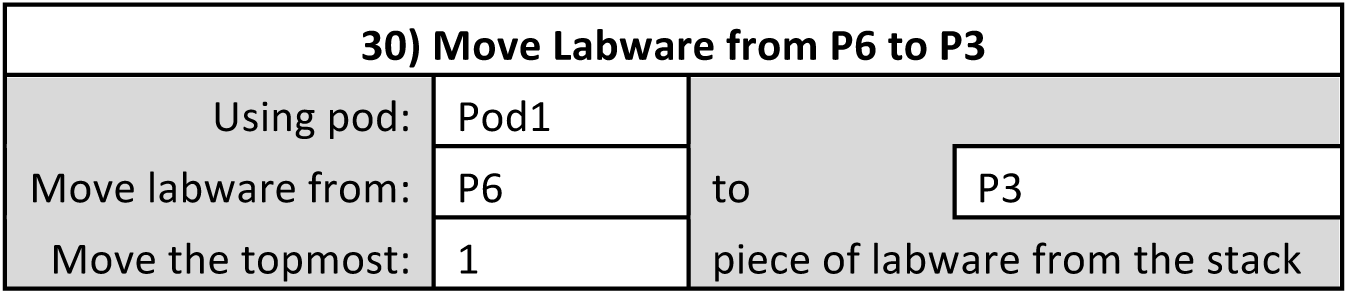

| 30) Move Labware from P6 to P3 |  |  |
| --- | --- | --- |
| Using pod: | Pod1 |  |
| Move labware from: | P6 | to P3 |
| Move the topmost: | 1 | piece of labware from the stack |

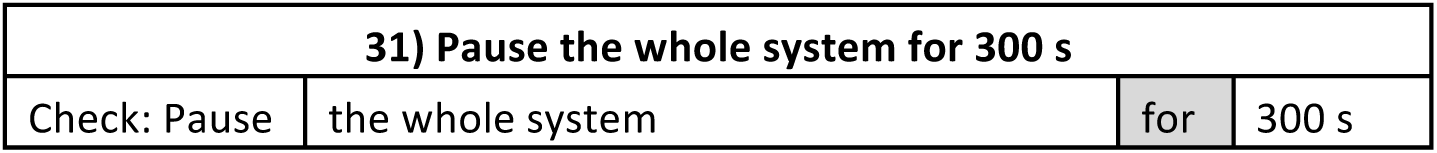

| 32) Load tips from TL1 |  |
| --- | --- |
| Tips: | Load |
| Location: | TL1 |
| Pod: | Pod1 |

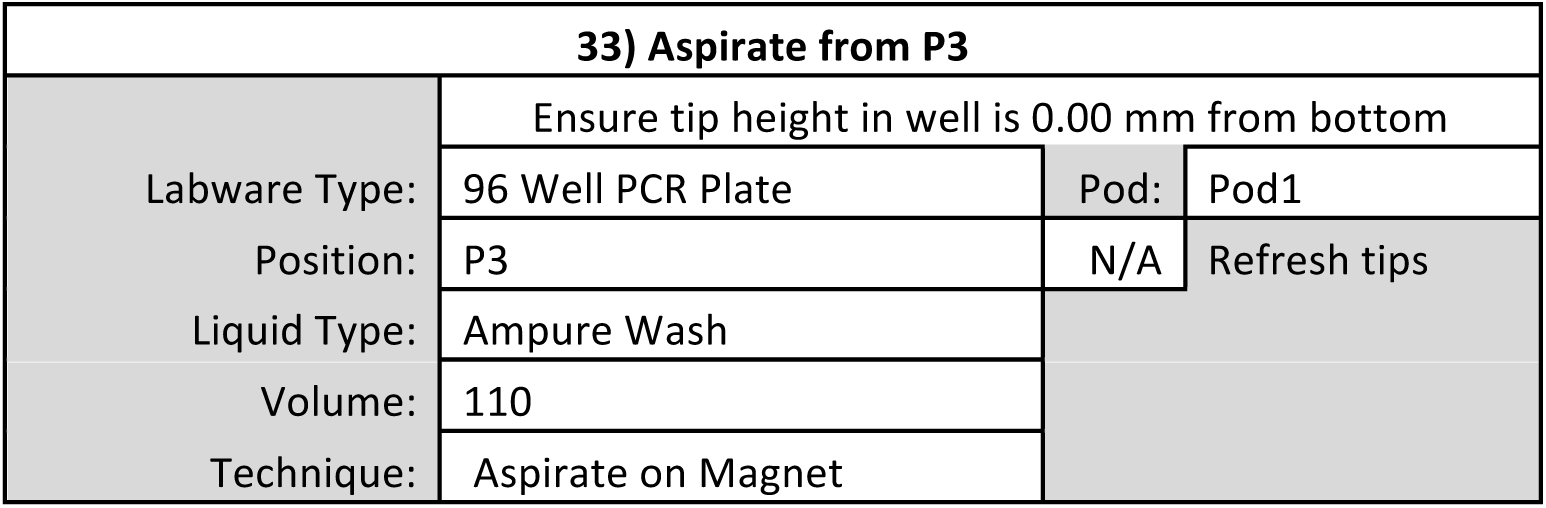

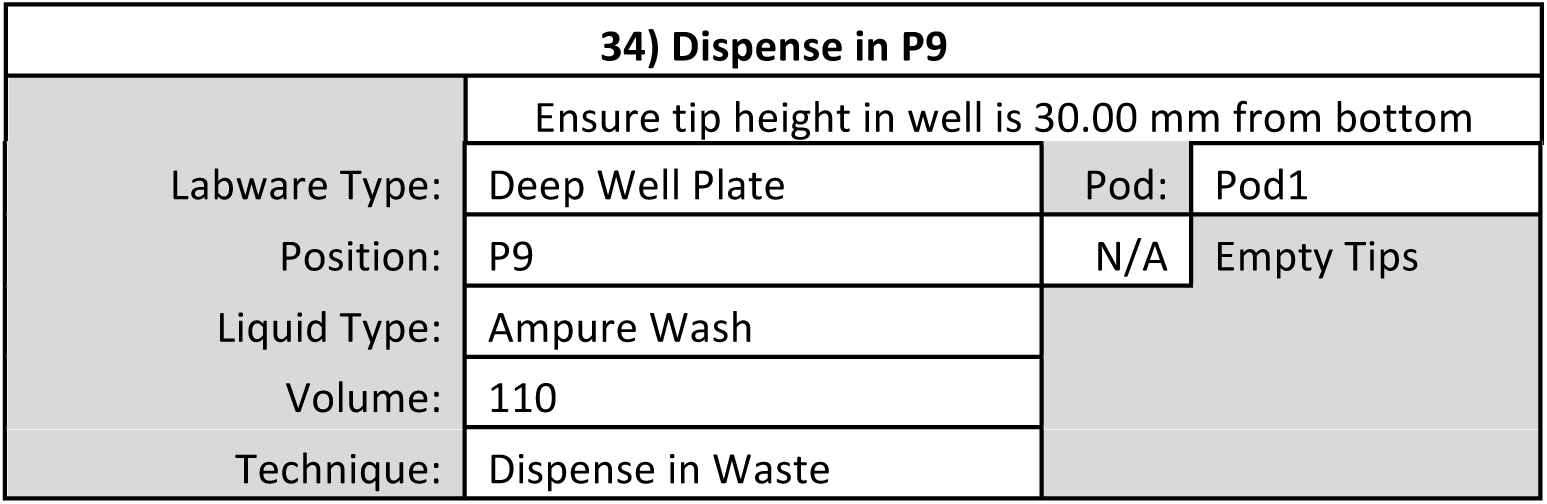

**35) Perform Steps 13-20 again to wash bead-bound DNA twice with Ethanol and then allow to air dry.**

| 36) Move Labware from P3 to P6 |  |  |  |
| --- | --- | --- | --- |
| Using pod: | Pod1 |  |  |
| Move labware from: | P3 | to | P6 |
| Move the topmost: | 1 | piece of labware from the stack |  |

| 37) Load tips from TL1 |  |
| --- | --- |
| Tips: | Load |
| Location: | TL1 |
| Pod: | Pod1 |

| 38) Aspirate from P2 |  |  |  |
| --- | --- | --- | --- |
|  | Ensure tip height in well is 1.5 mm from bottom |  |  |
| Labware Type: | V-Bottom Plate | Pod: | Pod1 |
| Position: | P2 | N/A | Refresh tips |
| Liquid Type: | Tris-HCl-1 |  |  |
| Volume: | 35 |  |  |
| Technique: | Aspirate from VBP |  |  |

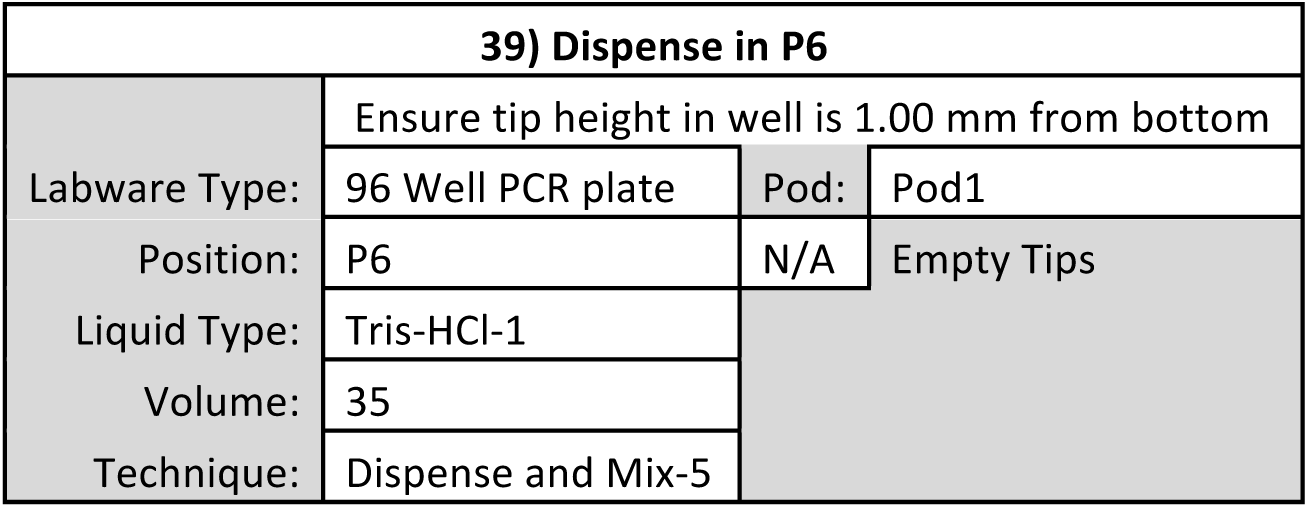

| 39) Dispense in P6 |  |  |  |
| --- | --- | --- | --- |
|  | Ensure tip height in well is 1.00 mm from bottom |  |  |
| Labware Type: | 96 Well PCR plate | Pod: | Pod1 |
| Position: | P6 | N/A | Empty Tips |
| Liquid Type: | Tris-HCl-1 |  |  |
| Volume: | 35 |  |  |
| Technique: | Dispense and Mix-5 |  |  |

| 40) Unload tips to TL1 |  |
| --- | --- |
| Tips: | Unload |
| Location: | TL1 |
| Pod: | Pod1 |

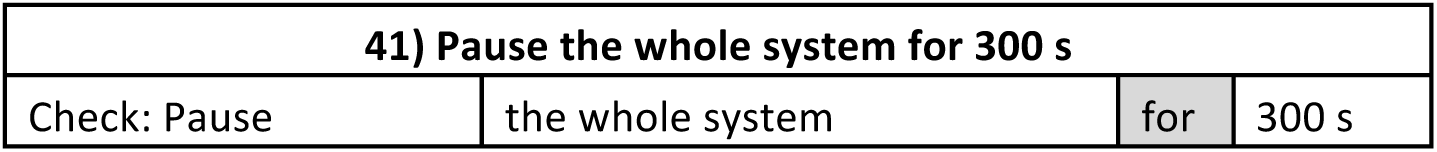

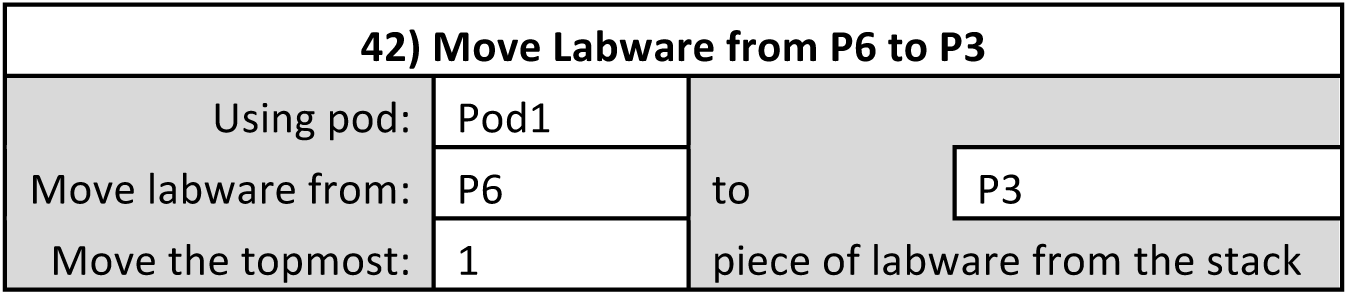

| 43) Pause the whole system for 300 s |  |  |  |
| --- | --- | --- | --- |
| Check: Pause | the whole system | for | 300 s |

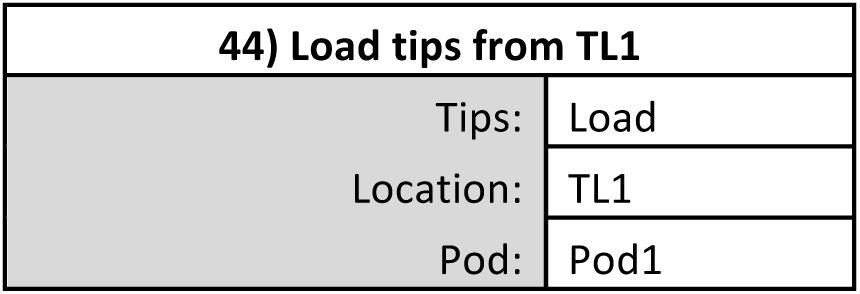

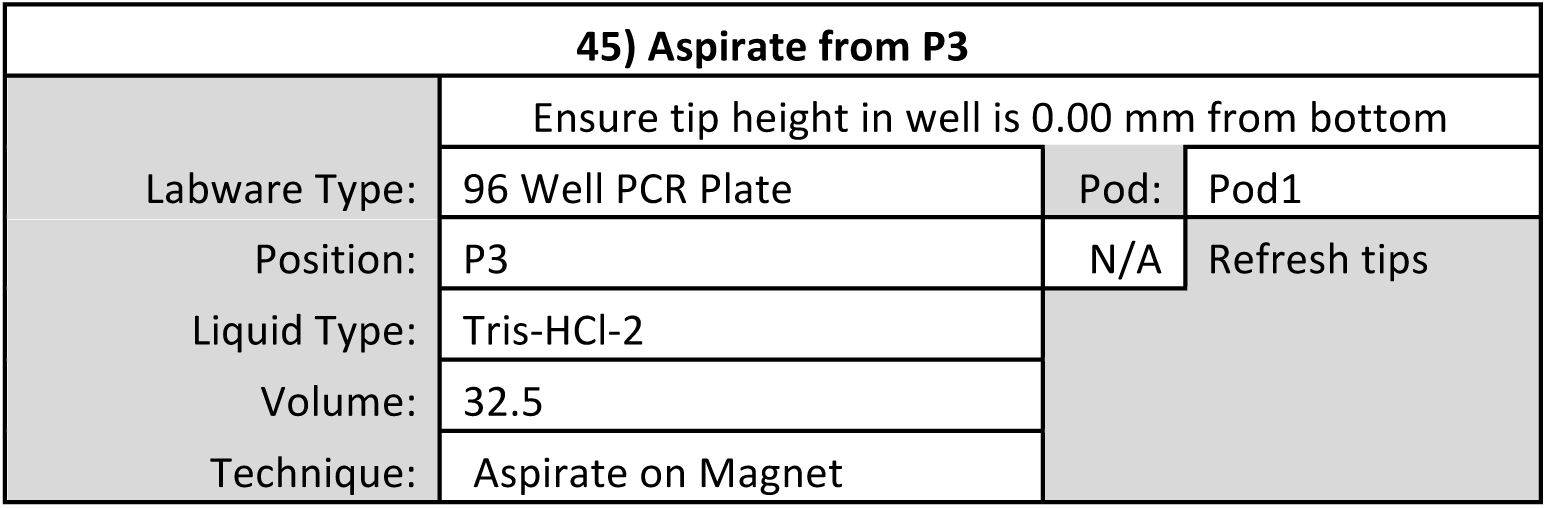

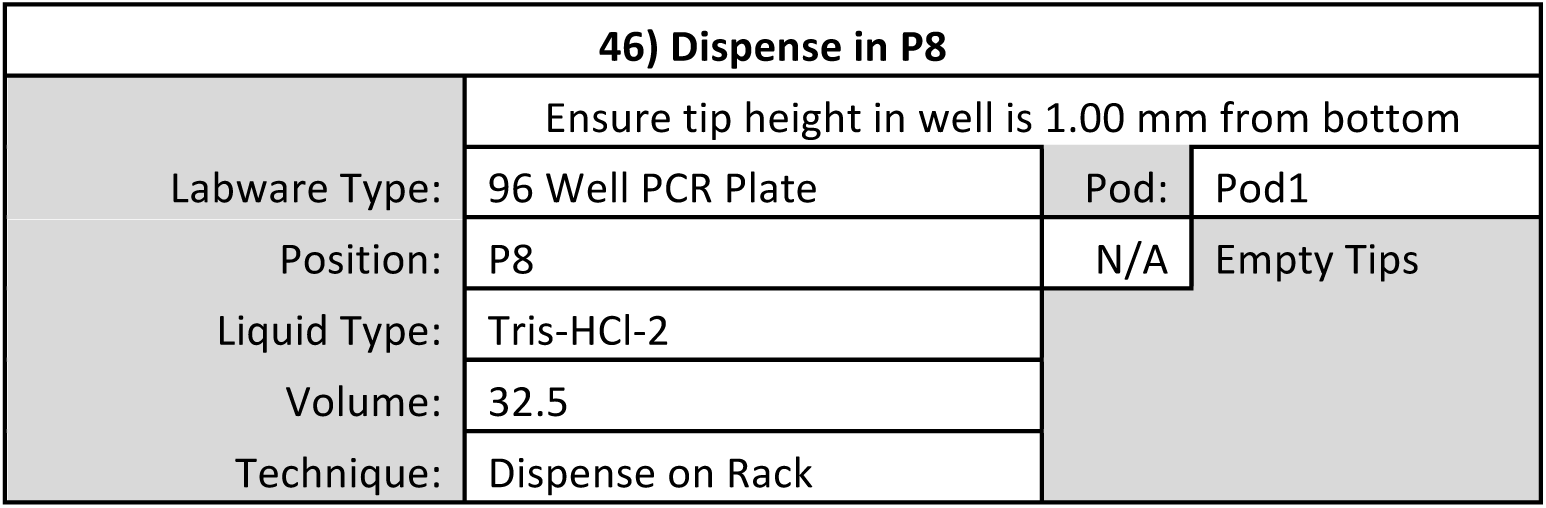

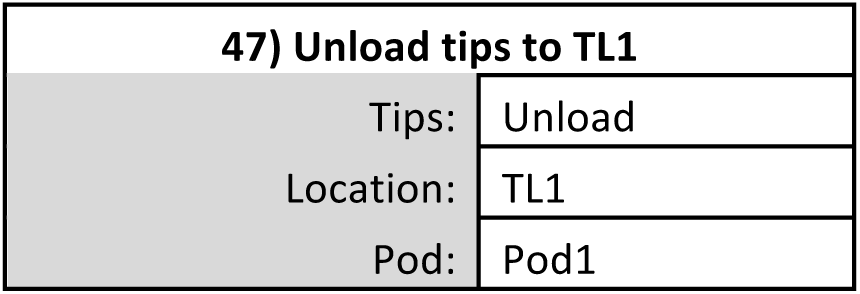

| 48) Finish |
| --- |
| Check: Clear current instrument setup of all labware after the method completes |
| Check: Clear current device setup of all labware after the method completes |
| Check: Unload disposable tips from all pods after the method completes |
| Check: Clear all global variables after the method completes |

### REAGENT SETUP

#### 5% Digitonin

1 mL of 5% digitonin is sufficient for up to 48 Auto CUT&RUN reactions, to perform up to 96 reactions prepare 2 mL of 5% digitonin. Weigh out the digitonin powder in a 2 mL microcentrifuge tube, boil water in a small beaker in a microwave oven. Pipette the hot water into the tube with the digitonin powder to make 5% (wt/vol), close the cap and quickly vortex on full. If digitonin is not completely dissolved seal with a tube-lock to prevent the cap from opening and place the tube in a 100°C heat block for 1-5 min. If saved and refrigerated, this stock can be used up to a week, but will need reheating as the digitonin slowly precipitates.

- **CAUTION:** Digitonin is toxic and PPE including a mask and gloves should be worn when weighing out the powder. A digitonin stock may be prepared by dissolving in dimethylsulfoxide (DMSO) but be aware that DMSO can absorb through the skin.

#### Binding buffer

Mix 20 mL of Binding Buffer in a 50 mL conical tube. Store the buffer at 4 °C for up to 6 months.

| Component | Amount | Final |
| --- | --- | --- |
| ddH<sub>2</sub>O | 19.36 mL | - |
| 1M HEPES-KOH pH 7.9 | 400 µL | 20 mM |
| 1M KCl | 200 µL | 10 mM |
| 1M CaCl<sub>2</sub> | 20 µL | 1 mM |
| 1M MnCl<sub>2</sub> | 20 µL | 1 mM |

#### Activate Concanavalin A-coated beads in Binding Buffer

Gently resuspend and withdraw enough of the bead suspension such that there will be 10 µL for each final sample. Transfer Concanavalin A-coated beads into 1.5 mL Binding Buffer in a 2 mL tube. For more than 48 reactions use a second 2 mL tube. Place tube(s) on a magnet stand to clear (30 s to 2 min). Withdraw the liquid and remove from the magnet stand. Add 1.5 mL Binding buffer, mix by inversion or gentle pipetting, remove liquid from the cap and side with a quick pulse on a micro-centrifuge. Place tube(s) on a magnet stand to clear (30 s to 2 min). Withdraw the liquid, then wash Concanavalin A-coated beads a second time with 1.5 mL of Binding buffer. After removing liquid from the second wash on a magnet stand, resuspend in a volume of Binding Buffer equal to the volume of bead suspension (10 µL per final sample).

#### Wash Buffer

50 mL of Wash Buffer is sufficient for up to 24 Auto CUT&RUN reactions. This buffer can be stored at 4 °C for up to 1 week, however, Roche Complete Protease Inhibitor tablet should be added fresh on the day of use.

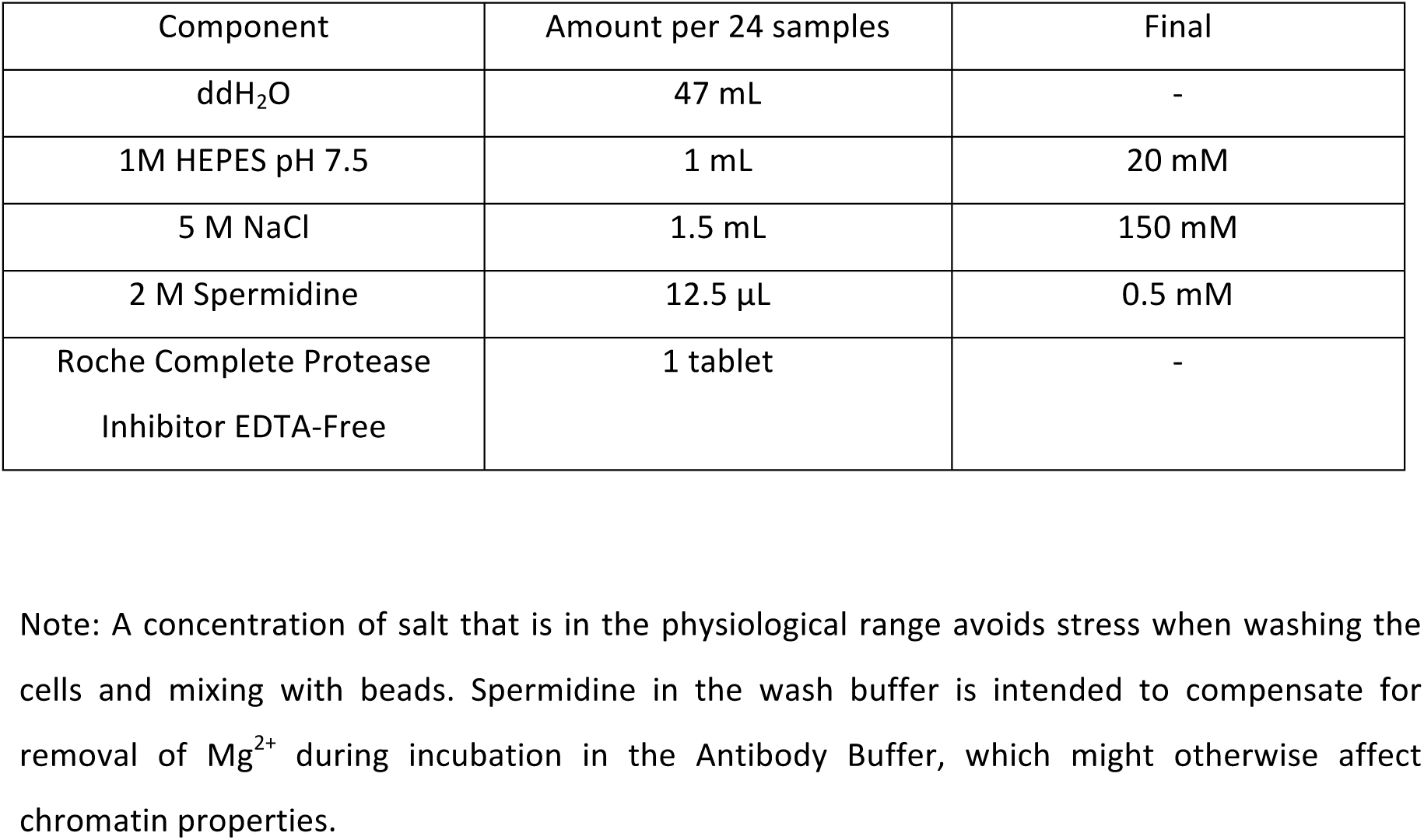

| Component | Amount per 24 samples | Final |
| --- | --- | --- |
| ddH<sub>2</sub>O | 47 mL | - |
| 1M HEPES pH 7.5 | 1 mL | 20 mM |
| 5 M NaCl | 1.5 mL | 150 mM |
| 2 M Spermidine | 12.5 μL | 0.5 mM |
| Roche Complete Protease Inhibitor EDTA-Free | 1 tablet | - |

Note: A concentration of salt that is in the physiological range avoids stress when washing the cells and mixing with beads. Spermidine in the wash buffer is intended to compensate for removal of Mg<sup>2+</sup> during incubation in the Antibody Buffer, which might otherwise affect chromatin properties.

#### Digitonin Buffer

For up to 24 Auto CUT&RUN reactions mix 150-600 µL 5% (wt/vol) digitonin with 30 mL Wash Buffer for a final concentration of digitonin between 0.025% and 0.1% (wt/vol). Store this buffer on ice or at 4 °C for up to 1 day, and vortex before use.

Note: The effectiveness of digitonin varies between batches, so testing for full permeability of Trypan blue is recommended to determine the concentration to use for a cell type. We have obtained excellent results for H1 and K562 cells with 0.05% digitonin (300 µL 5% (wt/vol) digitonin in 30 mL Wash Buffer). For simplicity, we use this same buffer for all steps starting from the incubation in primary antibody until the chromatin digestion.

#### Antibody Buffer

For up to 24 reactions (150 µL/reaction) mix 4 mL Digitonin Buffer with 16 µL 0.5 M EDTA and place on ice.

Note: The presence of EDTA during antibody treatment removes excess divalent cations used to activate the Concanavalin A-coated beads, as well as endogenous cations from the cells of interest. This serves to halt metabolic processes, stop endogenous DNAse activity, and prevent carry-over of Ca^2+^ from the Binding Buffer that might prematurely initiate strand cleavage after addition of pA-MNase. Washing out the EDTA before pA-MNase addition avoids inactivating the enzyme.

### PROCEDURE

- **CRITICAL STEP:** In order for the Biomek to perform Auto CUT&RUN as expected the labware must be properly calibrated and zeroed. The equipment specs provided in the Labware Type Editor Section should be empirically tested on your specific piece of equipment by setting up a short method in which the Biomek aspirates and dispenses liquid from each of the labware types at a height of 0 mm. The operator can then break the light screen to freeze the instrument and determine whether the pipette tips are indeed at the bottom of the well and then adjust the Labware Specs if necessary. The 96 Well PCR Plate should be zeroed while stacked on the PCR Plate Rack, ALPAQUA Magnet Stand and Cold Block, if necessary the Stack Offset Z parameter can be adjusted for each type of Rack independently. Properly zeroing the 96 Well PCR Plate on the ALPAQUA Magnet Stand should engage the spring without causing the stand to bottom out. Reducing the movement speed within the well during these initial calibration steps will minimize the possibility of the instrument becoming damaged if a crash occurs. In addition, prior to performing an Auto CUT&RUN reaction on biological samples it is recommended to test each method by pre-loading the labware with H_2_O containing a small amount of food coloring to improve visibility. The 96 well LoBind PCR plates, Semi-skirted (Eppendorf # 0030129504) are highly recommended because their dimensions are consistent enough from plate to plate to ensure the Biomek will also move and stack the plates in a consistent manner. To prevent the PCR Plate Racks from catching and being carried along with the 96 Well PCR Plate during movement steps, the PCR Plate Racks can be taped down to the stationary ALPs (this issue has not been observed for the ALPAQUA Magnet Stand or the Cold Block). Finally, the operator should remain present throughout each method for the first several runs to ensure the instrument performs as expected. In the event of a mishap the operator can then break the light curtain to immediately stop the procedure and intervene before the reaction is compromised.

### ? TROUBLESHOOTING

#### Binding cells to beads

- **TIMING 30 min**
- **CRITICAL STEP:** All steps prior to the addition of antibody are performed at room temperature (∼ 22 °C) to minimize stress on the cells. Because it is crucial that DNA breakage is minimized throughout the protocol, we recommend that cavitation during resuspension and vigorous vortexing be avoided.

1) Harvest fresh culture(s) at room temperature and count cells. The Auto CUT&RUN protocol is recommended for 50,000 to 1,000,000 mammalian cells per sample.

- **PAUSE POINT:** If necessary, cells can be cryopreserved in 10% (vol/vol) DMSO using a Mr. Frosty isopropyl alcohol chamber. We do not recommend flash freezing, as this can cause background DNA breakage that may impact final data quality.

2) Centrifuge 3 min 600 x g at room temperature and withdraw liquid.

3) Resuspend in 1.5 mL room temperature Wash Buffer by gently pipetting and transfer if necessary to a 2 mL tube.

4) Centrifuge 3 min 600 x g at room temperature and withdraw liquid.

5) Repeat steps 3 and 4 two more times.

- **CRITICAL STEP:** Thorough washing removes free sugars and other molecules that can compete for binding to the Concanavalin A coated-beads, ensuring efficient binding and recovery of the cells of interest.

6) Resuspend in 1 mL room temperature Wash Buffer by gently pipetting.

7) While gently vortexing the cells at room temperature, add the Concanavalin A coated-bead suspension.

8) Place on tube nutator at room temperature for 5-10 min.

#### Permeabilize cells and bind (primary) antibodies

- **TIMING 2 hrs–overnight**

9) Mix well by vigorous inversion to ensure the bead-bound cells are in a homogenous suspension and divide into aliquots in 0.6-mL low-bind tubes, one for each antibody to be used.

10) Place on the magnet stand to clear and pull off and discard the liquid.

11) Place each tube at a low angle on the vortex mixer set to low (∼1100 rpm) and squirt 150 µL of the Antibody Buffer per sample along the side while gently vortexing to allow the solution to dislodge most or all of the beads. Tap to dislodge the remaining beads.

- **CRITICAL STEP:** Permeabilizing the cells with digitonin and chelating divalent cations with EDTA serves to quickly halt metabolic processes and prevent endogenous DNAse activity. This helps to preserve the native chromatin state and reduce background noise in the final CUT&RUN libraries. Thus, it is recommended to work quickly to get cells into Antibody Buffer.

12) Mix in the primary antibody to a final concentration of 1:100 or to the manufacturer’s recommended concentration for immunofluorescence.

- **CRITICAL STEP:** To evaluate success of the procedure without requiring sequencing, include in parallel a positive control antibody (*e.g.* anti-H3K27me3) and a negative control antibody (*e.g.* anti-mouse IgG). Do not include a no-antibody control, as the lack of tethering may allow any unbound pA-MNase to act as a “time-bomb” and digest accessible DNA, resulting in a background of DNA-accessible sites.

13) Place on the tube nutator at room temperature for 2 hrs.

- **PAUSE POINT** Antibody incubation may proceed overnight at 4 °C.

### ? TROUBLESHOOTING

#### Bind secondary antibody (as required)

- **TIMING 1 hr-overnight**
- **CRITICAL STEP:** The binding efficiency of Protein A to the primary antibody depends on host species and IgG isotype. For example, Protein A binds well to rabbit and guinea pig IgG but poorly to mouse and goat IgG, and so for these latter antibodies a secondary antibody, such as rabbit anti-mouse is recommended.

14) Remove liquid from the cap and side with a quick pulse on a micro-centrifuge.

- **CRITICAL STEP:** After mixing, but before placing a tube on the magnet stand, a very quick spin on a micro-centrifuge (no more than 100 x g) will minimize carry-over of antibody and pA-MN that could result in overall background cleavages during the digestion step.

15) Place on the magnet stand to clear (∼30 s) and pull off all of the liquid.

16) Add 150 µL Digitonin Buffer to wash, mix by inversion, or by gentle pipetting using a 1 mL tip if clumps persist, and remove liquid from the cap and side with a quick pulse on a micro-centrifuge.

17) Repeat Digitonin Buffer wash steps 14-16.

18) Place on the magnet stand to clear and pull off all of the liquid.

19) Place each tube at a low angle on the vortex mixer set to low (∼ 1100 rpm) and squirt 150 µL of the Digitonin Buffer (per sample and/or digestion time point) along the side while gently vortexing to allow the solution to dislodge most or all of the beads. Tap to dislodge the remaining beads.

20) Mix in the secondary antibody to a final concentration of 1:100 or to the manufacturer’s recommended concentration.

21) Place on the tube nutator at 4 °C for ∼ 1 hr or overnight.

#### Bind Protein A-MNase fusion protein on Biomek

- **TIMING ∼ 1.5 hr**
- **CRITICAL STEP:** Performing the steps up until this point by hand increases the versatility of the platform, allowing individual users to harvest their cells or tissue of interest and bind any antibody of their choosing. Because the antibody incubation is not time sensitive, samples from multiple users can be synchronized at this step and arrayed on a single plate, allowing the remaining steps to be performed in unison on the Biomek by a single operator.

22) Set Cooling Unit (filled with antifreeze and routed to Heating/Cooling ALP with Aluminum Cooling Block at position P10) to 0 °C.

23) Prepare 4.5 mL of pA-MNase solution per 24 Auto CUT&RUN reactions by mixing the pA-MNase (supplied upon request) to a final concentration of ∼700 ng/mL in Digitonin Buffer. Transfer pA-MNase solution into a reservoir and dispense 175 µL into each of the active wells of a labeled V-Bottom Plate using a Multi-Channel Pipette. Place at position P4 on the Biomek deck.

- **CRITICAL STEP:** CUT&RUN is relatively insensitive to the concentration of pA-MNase as is evident from the titration test of two different batches, where increasing the concentration of pA-MNase above ∼100 ng/mL resulted in little additional release of H3K27me3-bound nucleosomes from 600,000 human cells after 30 min digestion in a 500 µL volume.

24) Using wide bore 200 µL tips, resuspend Concanavalin A bead-bound cells + Antibodies and array them in a 96 Well PCR Plate. Be sure to record the position of each sample in the plate and stack it on a PCR Plate Rack at position P6 on the Biomek deck.

25) Dispense 1 mL of Digitonin Buffer into each of the active wells of a labeled Deep Well Plate and place it at position P5 on the Biomek deck.

26) Place Fresh AP96 200 µL tips at position TL1, the ALPAQUA Magnet Plate at position P3, and an empty labeled Deep Well Plate for collecting liquid waste at position P9 on the Biomek deck.

27) Start the pA-MNase Binding Method. During the 1 hr mixing step continue on to prepare pA-MNase Reaction Mix and 4X Stop Buffer solutions and keep on Ice for the next step of the reaction.

*28)* ***Targeted chromatin digestion on the Biomek***

- **TIMING ∼1.5 hr**

29) Prepare pA-MNase Reaction Mix (50 µL/sample). Keep on Ice until use.

| Component | Amount per 24 samples (µL) | Added for Reservoir (µL) | Final volume per 24 samples (µL) | Final concentration |
| --- | --- | --- | --- | --- |
| Digitonin Buffer | 1176 | 441 | 1617 |  |
| 100mM CaCl<sub>2</sub><br>(diluted 1:10 from a 1 M stock) | 24 | 9 | 33 | 2mM |

30) Prepare 4X Stop Buffer (25 µL/sample). Keep on Ice until use.

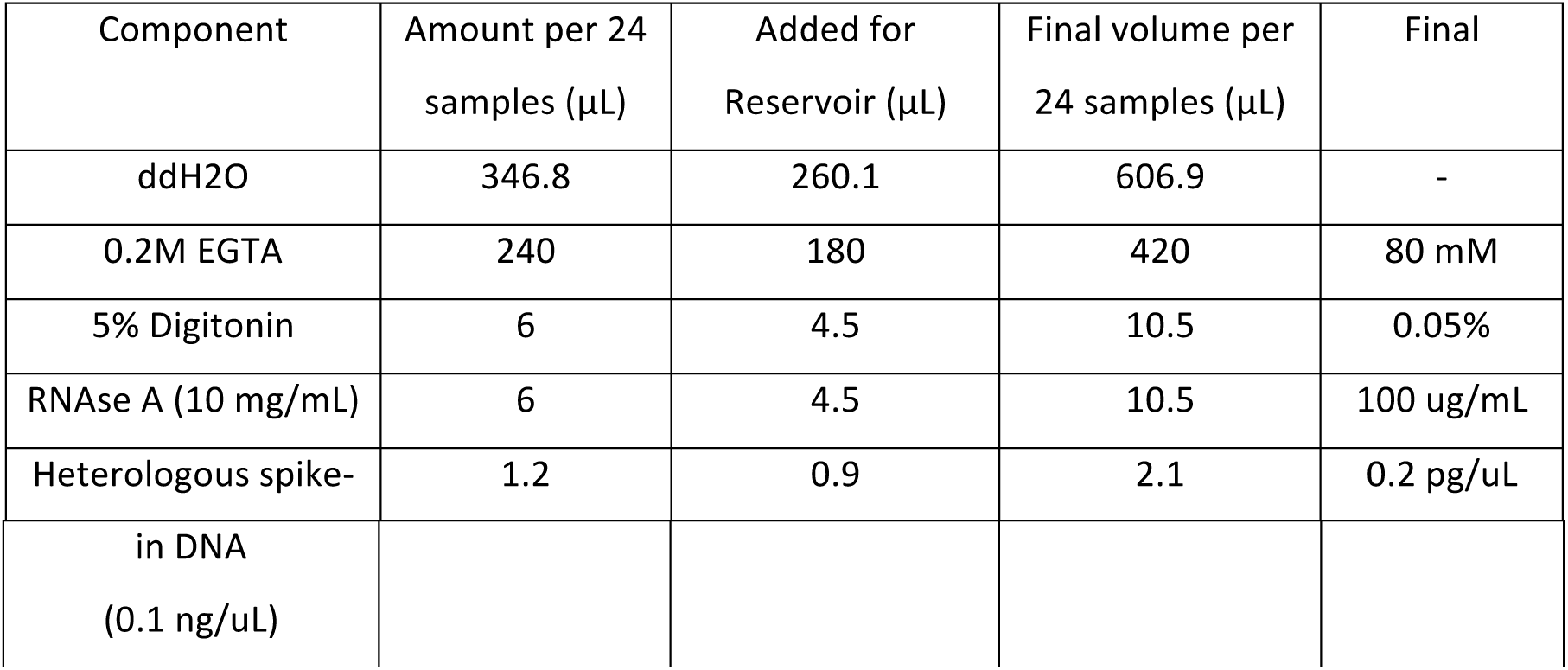

- **CRITICAL STEP:** Heterologous spike-in DNA is highly recommended for calibration, as there is too little background cleavage for normalization of samples. Spike-in DNA should be fragmented down to ∼200 bp mean size, for example an MNase-treated sample of mononucleosome-sized fragments. As we use the total number of mapped reads as a normalization factor only, very little spike-in DNA is needed. For example, addition of 1.5 pg results in 1,000-10,000 mapped spike-in reads for 1-10 million mapped experimental reads (in inverse proportion).

31) When the pA-MNase Binding Method is complete, remove the labeled V-Bottom plate containing residual pA-MNase solution and wash well with DI water (this plate can be reused in subsequent experiments). Then empty the liquid waste from the labeled Deep Well Plate and again place it at position P9 on the Biomek deck.

32) Using a Reservoir and Multi-Channel Pipette dispense 50 µL of pA-MNase Reaction Mix into the active wells of a labeled V-Bottom Plate and place it at position P4 on the Biomek deck. Dispense 25 µL of 4X STOP Buffer in the active wells of a labeled V-Bottom Bottom Plate and place it at position P7 on the Biomek deck. Gently shake the Deep Well Plate containing Digitonin Buffer to ensure digitonin remains suspended in solution and again place it at position P5 on the Biomek deck.

33) Place a fresh 96 Well PCR Plate in a PCR Plate Rack at position P8 for accepting digested chromatin. The AP96 200 µL Tips at TL1 can be used again for this method, the ALPAQUA Magnet Plate should remain at position P3, and a free PCR Plate Rack should remain at position P6. The 96 well PCR Plate containing Concanavalin A bead-bound cells + Antibodies + pA-MNase should remain on the Cooling Block set to 0°C at position P10 on the Biomek deck.

34) Ensure the Pause at Step 19 of the pA-MNase Digest Method (after addition of MNase Reaction Mix and before addition of 4X STOP Buffer) is set for the desired digestion time.

- **CRITICAL STEP:** Longer digestion times tend to increase the yield of digested chromatin from each reaction, and can be critical to obtain quality signal from low abundance epitopes (e.g. transcription factors). However, for some antibodies (e.g. anti-H3K27ac) longer digestion times have been observed to increase the amount of non-specific or “background” DNA that is recovered. Because the current platform adds 4X STOP Buffer to all the wells synchronously, setting individual digestion lengths for each reaction is currently not an option, therefore a digestion time of 9-30 min is recommended.

35) Start the pA-MNase Digest Method. During the chromatin release step, when the plate is mixing and incubating at room temperature for about half an hour, prepare 4X End Repair and A-Tailing Buffer and keep on Ice for the next step of the reaction.

#### Chromatin end repair and dA-tailing

- **TIMING ∼2.5 hr**

36) Prepare 4X End Repair and A-tailing (ERA) buffer (12.5 µL/sample). Keep on Ice until use.

| Component | Amount per 24 samples ($\mu\text{L}$) | Added for Reservoir ($\mu\text{L}$) | Final volume per 24 samples ($\mu\text{L}$) | Final concentration |
| --- | --- | --- | --- | --- |
| T4 DNA ligase Buffer (10X) | 120 | 180 | 300 | 4X |
| dNTPs (10mM each) | 60 | 90 | 150 | 2 mM each |
| ATP (10mM) | 30 | 45 | 75 | 1 mM |
| 40% PEG 4000 | 75 | 112.5 | 187.5 | 10% |
| T4 PNK (10 U/$\mu\text{L}$) | 15 | 22.5 | 37.5 | 0.5 U/$\mu\text{L}$ |
| T4 DNA Polymerase (5U/$\mu\text{L}$) | 3 | 4.5 | 7.5 | 0.05 U/$\mu\text{L}$ |
| Taq DNA Polymerase (5U/$\mu\text{L}$) | 3 | 4.5 | 7.5 | 0.05 U/$\mu\text{L}$ |

37) When the pA-MNase Digestion Method is complete, remove the labeled V-Bottom plates containing residual pA-MNase Reaction Mix and 4X STOP Buffer, as well as the labeled Deep Well Plates containing residual Digitonin Buffer and Liquid Waste from the Biomek deck. Empty the plates and wash well with DI water (these plates can be reused in subsequent experiments). Seal the 96 Well PCR Plate containing the digested Con-A bead-bound cells (position P3) with an adhesive cover and store at −20 °C for potential troubleshooting.

38) Remove the 96 Well PCR Plate containing digested chromatin (position P8) from the Biomek deck. Using a Reservoir and Multi-Channel Pipette add 12.5 µL 4X ERA buffer to each sample and mix by pipetting up and down 5 times.

39) Seal the 96 Well PCR Plate using an adhesive cover and place in a thermocycler that has been pre-cooled to 12°C and run the following program with heated lid for temp ≥20°C:

| Cycle number | End Polishing | 5' End phosphorylation | dA-Tailing | Final |
| --- | --- | --- | --- | --- |
| 1 | 12°C, 15 min |  |  |  |
| 2 |  | 37° C, 15 min |  |  |
| 3 |  |  | 58° C, 1.5 hour |  |
| 17 |  |  |  | 8° C, hold |

- **CRITICAL STEP:** The 58° C, 1.5 hour dA-Tailing step is optimized to prevent short fragments (30-120bp) from denaturing, while still heat inactivating the exonuclease activity of the T4 DNA Polymerase.

#### Adapter Ligation

- **TIMING 1 hr-overnight**

40) Remove pre-annealed 0.15 µM TruSeq adapters from freezer and allow to thaw on ice.

- **CRITICAL STEP:** To facilitate multiplexing during sequencing each adapter includes a six-nucleotide index or “barcode”. For subsequent data analysis make sure to keep track of which adapter is used for each sample. To allow for addition of adapters using a Multi-Channel Pipette 0.15 µM pre-annealed TruSeq adapters can be arrayed in a 96 Well PCR plate.

41) Prepare 2X Rapid Ligase solution (50 µL/ sample). Keep on Ice until use.

| Component | Amount per 24 samples (μL) | Added for Reservoir (μL) | Final volume per 24 samples (μL) | Final concentration |
| --- | --- | --- | --- | --- |
| Enzymatics 2X Rapid DNA ligase buffer | 1080 | 405 | 1485 | ~2 X |
| Enzymatics DNA Ligase | 120 | 45 | 165 |  |

- **Note:** Measure volumes carefully as the Enzymatics ligase buffer solution is very viscous. Prior to use pipette 2X Rapid Ligase solution up and down 10X to ensure adequate mixing - 42) Using a Multi-Channel Pipette, add 5 μL of 0.15 μM pre-annealed TruSeq adapters to each sample and mix by pipetting up and down 5 times.

42) Using a Multi-Channel Pipette, add 5 µL of 0.15 µM pre-annealed TruSeq adapters to each sample and mix by pipetting up and down 5 times.

43) Using a Reservoir and Multi-Channel Pipette, add 50 µL 2X Rapid Ligase solution to each sample and mix well by pipetting up and down 10 times.

44) Seal the 96 Well PCR Plate using an adhesive cover and place in a thermocycler that has been pre-cooled to 20°C and incubate at 20°C for 20 min. Following incubation proceed with Proteinase K digestion immediately.

45) Using a Reservoir and Multi-Channel Pipette, add 2 µL 10% SDS and 2 µL Proteinase K (20 mg/ml) to each sample. Seal the 96 Well PCR Plate using an adhesive cover and mix well by vortexing. Spin down solution at 2,000 rpm for 1 min and incubate in a thermocycler at 37°C for 30 min-overnight.

#### Pre-PCR DNA Cleanup on the Biomek

- **TIMING∼2 hrs**
- **CRITICAL STEP:** Two rounds of Ampure Bead Cleanup are performed prior to PCR amplification to remove un-ligated, and self-ligated adapter as well as unwanted protein, PEG, and salt.

46) Remove Ampure Bead Slurry from refrigerator, resuspend beads by vortexing and allow to equilibrate to room temperature before using.

47) Spin 96 Well PCR Plate containing Adapter Ligated DNA samples @2,000 rpm for 1 min. Remove seal and stack on a PCR Plate Rack positioned at P5 on the Biomeck deck.

48) Using a Reservoir and Multi-Channel Pipette distribute 90 µL of the Ampure Bead Slurry into the active wells of a 96 Well PCR Plate and stack on a PCR Plate Rack positioned at P6 on the Biomeck deck.

49) Dispense 1 mL of 80% Ethanol into each of the active wells of a labeled Deep Well Plate and place at position P4 on the Biomek deck.

50) For up to 24 Auto CUT&RUN reactions prepare 6 mL of HXP Mix (20% PEG 8000, 2.5M NaCl). Using a Reservoir and Multi-Channel Pipette distribute 100 µL of the HXP Mix into the active wells of a V-Bottom Plate and place at position P7 on the Biomek deck.

- **CRITICAL STEP:** HXP Mix is light sensitive and will degrade over time once mixed. Therefore, the HXP Mix should either be prepared fresh for each use or stored in the dark at −20°C.

51) Using a Reservoir and Multi-Channel Pipette distribute 100 µL of 10mM Tris-HCl pH 8 into the active wells of a labeled V-Bottom Plate and place at position P2 on the Biomek deck.

52) Place Fresh AP96 200 µL Tips at TL1, a labeled Deep Well Plate for accepting liquid waste at position P9, and a fresh 96 well PCR plate stacked on a PCR Plate Rack for accepting clean Adapter Ligated DNA in position P8 on the Biomek deck.

53) Start the Pre-PCR DNA Cleanup Method. After about 30 min continue on to prepare KAPA PCR Master Mix and keep on Ice for the next step of the reaction.

#### PCR Amplification of CUT&RUN Libraries

- **TIMING 1 hr-overnight**

54) Prepare KAPA PCR Master Mix.

| Component | Amount per 24 samples ($\mu\text{L}$) | Added for Reservoir ($\mu\text{L}$) | Final volume per 24 samples ($\mu\text{L}$) | Final Concentration |
| --- | --- | --- | --- | --- |
| 5X KAPA buffer | 240 | 225 | 465 | $\sim 2.5 \times$ |
| 10 mM dNTPs | 36 | 33.75 | 69.75 | 0.75 mM |
| 20 $\mu\text{M}$ P5 primer | 120 | 112.5 | 232.5 | 5 $\mu\text{M}$ |
| 20 $\mu\text{M}$ P5 primer | 120 | 112.5 | 232.5 | 5 $\mu\text{M}$ |
| KAPA HS HIFI polymerase | 24 | 22.5 | 46.5 |  |

55) When the Pre-PCR DNA Cleanup Method is complete, remove the labeled V-Bottom Plates containing residual 10mM Tris-HCl pH 8 and HXP Mix, as well as the labeled Deep Well Plates containing residual Ethanol and Liquid Waste from the Biomek deck. Empty the plates and wash well with DI water (these plates can be reused in subsequent experiments).

56) Remove 96 well PCR plate containing clean Adapter Ligated DNA (position P8) from the Biomek deck. Using a Reservoir and Multi-Channel Pipette add 20 µL KAPA PCR Master Mix to each sample and mix well by pipetting up and down 10 times.

57) Seal the PCR plate using an adhesive cover, place in thermocycler and run the following program with heated lid:

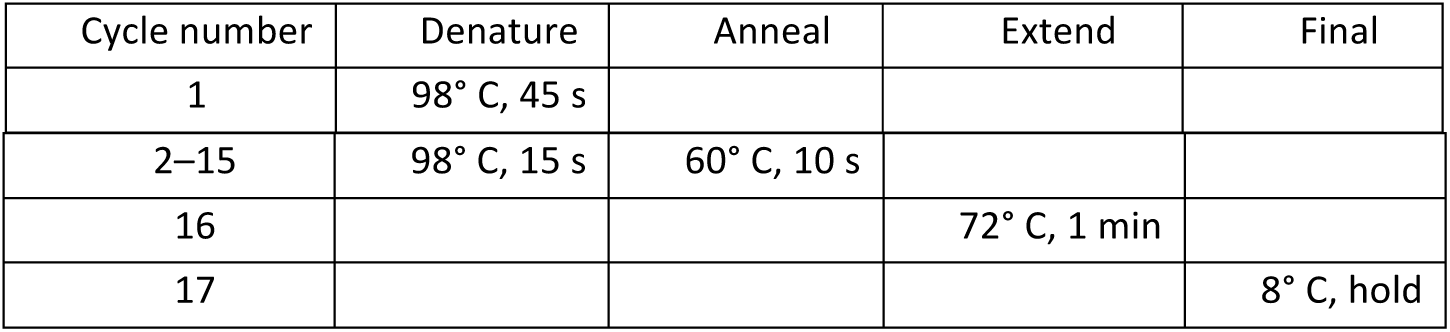

| Cycle number | Denature | Anneal | Extend | Final |
| --- | --- | --- | --- | --- |
| 1 | 98° C, 45 s |  |  |  |
| 2–15 | 98° C, 15 s | 60° C, 10 s |  |  |
| 16 |  |  | 72° C, 1 min |  |
| 17 |  |  |  | 8° C, hold |

- **CRITICAL STEP:** To minimize the contribution of large DNA fragments, PCR cycles should be at least 12-14 cycles, preferably with a 10 s 60°C combined annealing/extension step. Good results have been obtained with the Hyper-prep kit (KAPA Biosystems).

#### Post-PCR DNA Cleanup on the Biomek

- **TIMING ∼2 hr**
- **CRITICAL STEP:** Two rounds of Ampure Bead Cleanup are performed following PCR amplification to remove unused P5 and P7 primers, amplified self-ligated adapters and unwanted protein and salt. Once amplified, PCR products of a CUT&RUN library can serve as template for subsequent reactions using the P5 and P7 primers, making cross-contamination of Pre-PCR samples a serious concern. Therefore, all Post-PCR reagents should be stored separately from Pre-PCR reagents, and prepared in a separate area. If labware is to be reused, separate plates should be clearly labeled for use in Post-PCR reactions only.

58) Remove Ampure Bead Slurry from refrigerator, resuspend beads by vortexing and allow to equilibrate to room temperature before using.

59) Spin 96 Well PCR Plate containing PCR amplified CUT&RUN libraries @2,000 rpm for 1 min. Remove seal and stack on a PCR Plate Rack positioned at P5 on the Biomeck deck.

60) Using a Reservoir and Multi-Channel Pipette distribute 55 µL of the Ampure Bead Slurry into the active wells of a 96 Well PCR Plate and stack on a PCR Plate Rack positioned at P6 on the Biomeck deck.

61) Dispense 1 mL of 80% Ethanol into each of the active wells of a labeled Deep Well Plate and place at position P4 on the Biomek Deck.

62) Using a Reservoir and Multi-Channel Pipette distribute 100 µL of the remaining HXP Mix into the active wells of a V-Bottom Plate and place at position P7 on the Biomek deck.

63) Using a Reservoir and Multi-Channel Pipette distribute 100 µL of 10mM Tris-HCl pH 8 into the active wells of a V-Bottom Plate and place at position P2 on the Biomek deck.

64) Place Fresh AP96 200 µL Tips at TL1, a labeled Deep Well Plate for accepting liquid waste at position P9, and a fresh 96 well PCR plate stacked on a PCR Plate Rack for accepting clean CUT&RUN DNA Libraries at position P8 on the Biomek deck.

65) Start the Post-PCR DNA Cleanup Method. This Step should be complete in approximately 1.5 hr.

#### Sequencing

- **TIMING 1-2d**

66) When the Post-PCR DNA Cleanup Method is complete, remove the labeled V-Bottom plates containing residual 10mM Tris-HCl pH 8 and HXP Buffer, as well as the labeled Deep Well Plates containing residual Ethanol and Liquid Waste from the Biomek deck. Empty the plates and wash extremely well with DI water (these plates can be reused in subsequent experiments).

67) Remove 96 Well PCR Plate containing clean CUT&RUN Libraries (position P8) from the Biomek deck and seal with an adhesive cover for subsequent analysis.

68) Determine the size distribution of libraries by Agilent 4200 TapeStation analysis.

### ? TROUBLESHOOTING

69) Quantify library yield using dsDNA-specific assay, such as Qubit.

70) Pool indexed CUT&RUN libraries and perform paired-end Illumina sequencing following the manufacturer’s instructions.

- **CRITICAL STEP:** Because of the very low background with CUT&RUN, typically 5 million paired-end reads suffices for transcription factors or nucleosome modifications, even for the human genome. For maximum economy, we mix up to 24 barcoded samples per lane at equimolar concentration (proved a similar number of reads is desired for each sample) and perform paired-end 25−25 bp sequencing on a 2-lane flow cell. Single-end sequencing is not recommended for CUT&RUN, as it sacrifices resolution and discrimination between transcription factors and neighboring nucleosomes.

#### Data processing and analysis

- **TIMING1 d (variable)**

71) We align paired-end reads using Bowtie2 version 2.2.5 with options: --local --very-sensitive-local --no-unal --no-mixed --no-discordant --phred33 -I 10 -X 700. For mapping spike-in fragments, we also use the --no-overlap --no-dovetail options to avoid cross-mapping of the experimental genome to that of the spike-in DNA.

- **CRITICAL STEP:** Separation of sequenced fragments into ≤120 bp and ≥150 bp size classes provides mapping of the local vicinity of a DNA-binding protein, but this can vary depending on the steric access to the DNA by the tethered MNase.

**? TROUBLESHOOTING**

### TROUBLESHOOTING

**Table 2:** Troubleshooting table.

| Steps | Problem | Possible reasons | Solutions |
| --- | --- | --- | --- |
| Biomek Setup | Liquid volume is inconsistent between wells | The Biomek aspirate and or dispense function is imprecise from well to well potentially due to a clogged line or pump | Avoid using problematic wells. Contact Beckman Coulter technician, and possibly replace the P200 head. |
| Biomek Setup | Incorrect volume is being aspirated or dispensed in all wells | Aspiration is not occurring at the desired height within the well. The Biomek aspirate and or dispense function is not accurate. | Adjust aspiration height in the Method. Adjust volumes for desired result as determined empirically. Contact Beckman Coulter technician and possibly recalibrate the machine. |
| 13 | Beads clump and cannot be disaggregated | Cells lyse | Reduce the digitonin concentration. Use Non-stick tubes |
| 67 | No DNA is detected by Tape Station analysis | This indicates the reaction failed and could be due to (1) failed CUT&RUN reaction or (2) failed Library Prep. | Run a positive control sample for an abundant epitope, *e.g.* H3K27me3. Ensure Bioimek is pipetting accurately *e.g.* no Ethanol remains after 10 min Air Dry. |
| 67 | A prominent peak at ~120bp is detected by Tape station analysis in a large number of samples | This peak corresponds to self-ligated adapters and can persist after cleanup when relatively little digested chromatin is released (*i.e.* low cell numbers or low abundance epitopes). | Perform additional rounds of Ampure Cleanup. Reduce the ratio of Ampure Beads or HXP Mix to sample. Increase pA-MNase digestion time. |
| 67 | A small peak at | This is typical and indicates a small | Pool Libraries then |
|  | ~120bp is detected by Tapestation analysis in a few samples | amount of self-ligated adapter remains in these libraries. | perform one round of Ampure Cleanup before sequencing |
| 67 | No DNA <300bp is detected by Tapestation analysis | Sub-nucleosomal particles (often protected by transcription factors) are being denatured during the end repair and ligation or being removed during library cleanup. | Ensure the dA-Tailing step is at 58 degrees. Increase the concentration of Adapters. Increase the ratio of Ampure Beads or HXP Mix to sample. |
| 70 | Data quality from a sample of interest is indistinguishable from the IgG control | This indicates the reaction failed possibly due to an antibody failure or over digestion by pA-MNase. | Increase antibody concentration. Test antibody binding by adding a fluorescent secondary and imaging. Reduce digestion time. Replace antibody. |

### TIMING

**Days 1-3 Cells to Libraries**

Steps 1-8, binding cells to beads: 30 min

Steps 9-13, permeabilize cells and bind (primary) antibody: 2 hr-overnight

Steps 14-21 (Optional), bind secondary antibody: 1 hr-overnight

Steps 22-27, bind Protein A-MNase fusion protein on Biomek: ∼1.5 hr

Steps 28-34, targeted chromatin digestion on Biomek: ∼1.5 hr

Steps 35-38, chromatin end repair and dA-tailing: ∼2.5 hr

Steps 39-44, adapter ligation: 1 hr – overnight

Steps 45-52, Pre-PCR DNA cleanup on the Biomek: ∼2 hr

Steps 53-56, PCR amplification of CUT&RUN Libraries: 1 hr-overnight

Steps 57-64, Post-PCR DNA cleanup on the Biomek: ∼2 hr

**Days 4-6 Sequencing**

Step 65-69, sequencing: 1-2 days

**Day 7 (variable) Data processing and analysis**

Step 70, ≥1 day

**Supplementary Figure 1.**
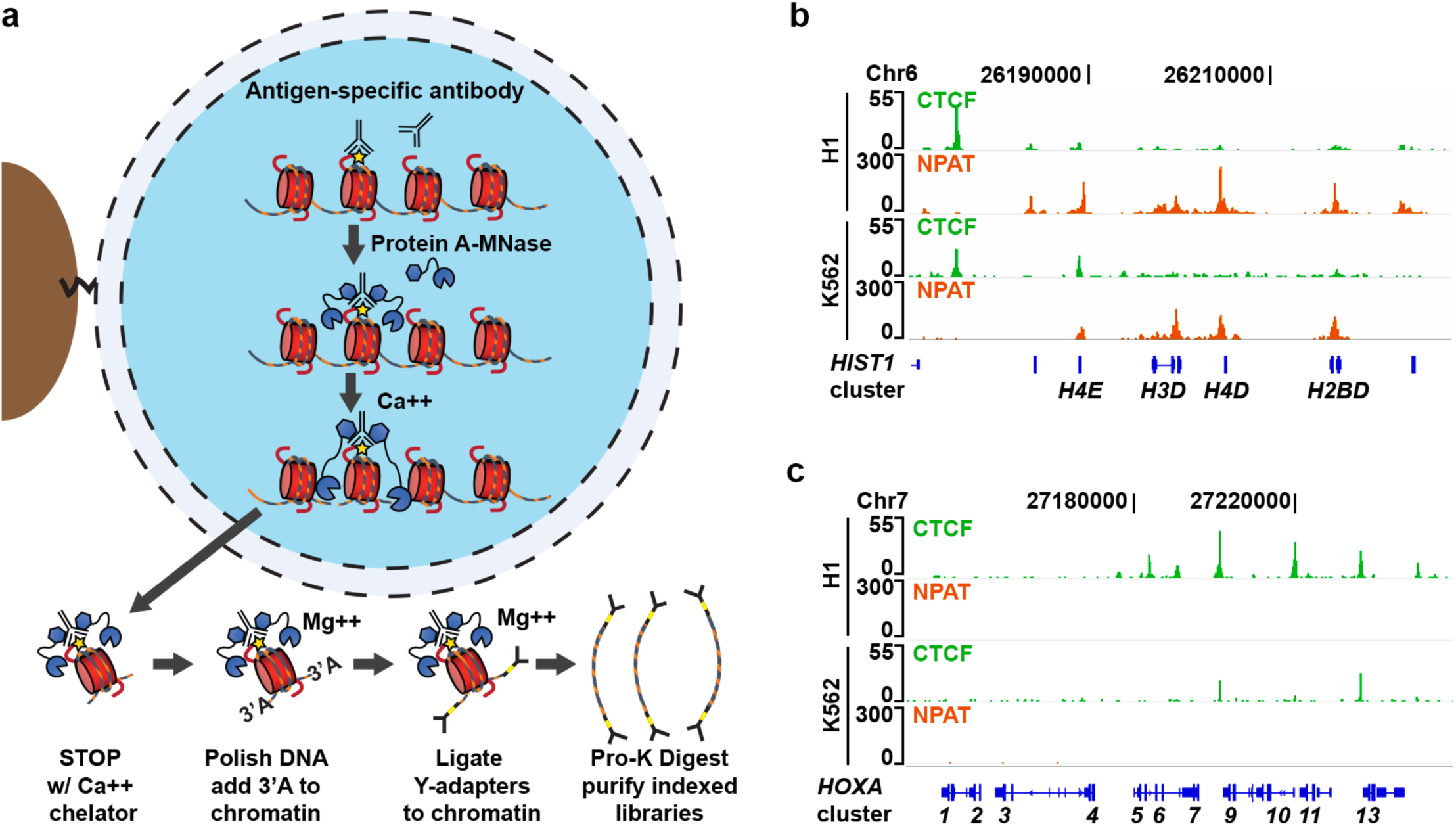
AutoCUT&RUN accurately maps NPAT and CTCF. (**a**) A modified CUT&RUN protocol allows for automation. ConA bead-bound samples are incubated with a chromatin protein-specific antibody, and arrayed on the Biomek for successive washes, tethering of a proteinA-MNase fusion protein, and cleavage of DNA by adding Ca2+. To avoid having to purify the digested DNA prior to library prep, the reaction is stopped with and EGTA only STOP buffer which specifically chelates Ca2+, but leaves adequate Mg2+ to allow End-polishing and Ligation of Illumina Y-adapters to the chromatin fragments. Chromatin protein is then digest with Proteinase-K and the indexed CUT&RUN libraries are purified on the Biomek using Ampure Magnetic Beads. (**b**) Genome browser tracks of NPAT and CTCF AutoCUT&RUN showing NPAT is enrichment at promoters of the *HIST1* gene cluster in both H1 and K562 cells. (**c**) Genome browser tracks confirming CTCF is bound to insulator regions in the *HOXA* locus.

**Supplementary Figure 2.**
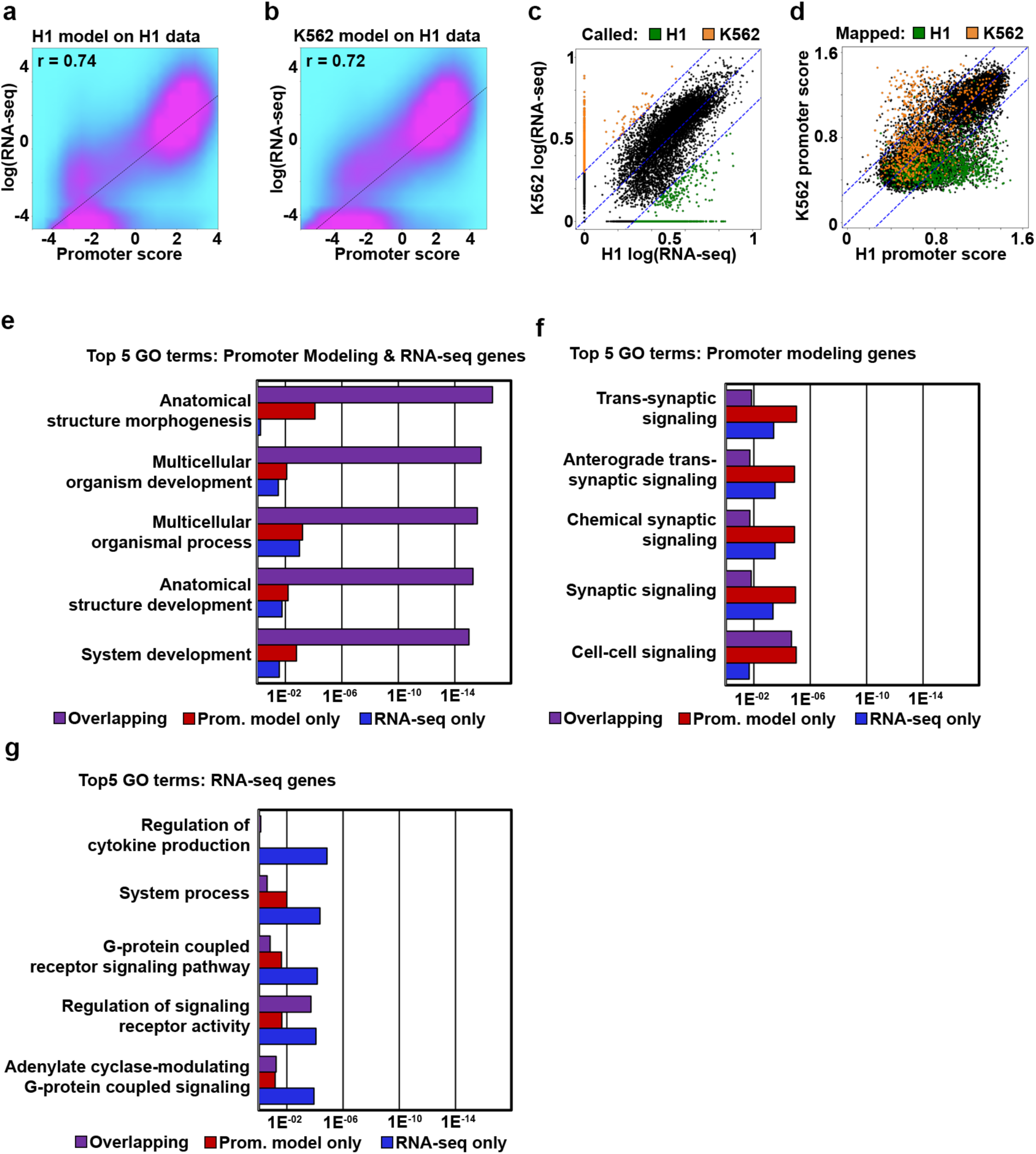
Developing a linear regression model to predict the activity of cis-regulatory elements. (**a**) Density scatterplot comparing H1 RNA-seq values for single-promoter genes to H1 promoter scores predicted by the model trained on H1 data. (**b**) Density scatterplot comparing H1 RNA-seq values for single-promoter genes to H1 promoter scores predicted by the model trained on K562 data. (**c**) Scatterplot of RNA-seq values for single-promoter genes in H1 and K562 cells. Colored dots indicate the RNA expression levels are ≥2-fold enriched in either H1 cells (green) or K562 cells (orange). (**d**) Scatterplot showing the distribution of genes with RNA-seq values that are ≥2-fold enriched in either H1 cells (green) or K562 cells (orange) mapped onto their corresponding promoter chromatin scores. Blue dotted lines indicated the 2-fold difference cut-off. (**e**) Bar graph showing the top five Gene Ontology (GO) terms overrepresented in the collection of cell-type specific genes identified by both promoter activity modeling as well as RNA-seq (purple), and the relative enrichment of these terms in the collections of genes uniquely identified as cell-type specific by promoter activity modeling (red) or RNA-seq (blue). (**f**) Bar graph showing the top five GO terms overrepresented in the collection of genes identified as cell-type specific according to promoter modeling scores only. (**g**) Bar graph showing the top five showing GO terms overrepresented in the collection of genes uniquely identified as cell-type specific according to RNA-seq.

**Supplementary Figure 3.**
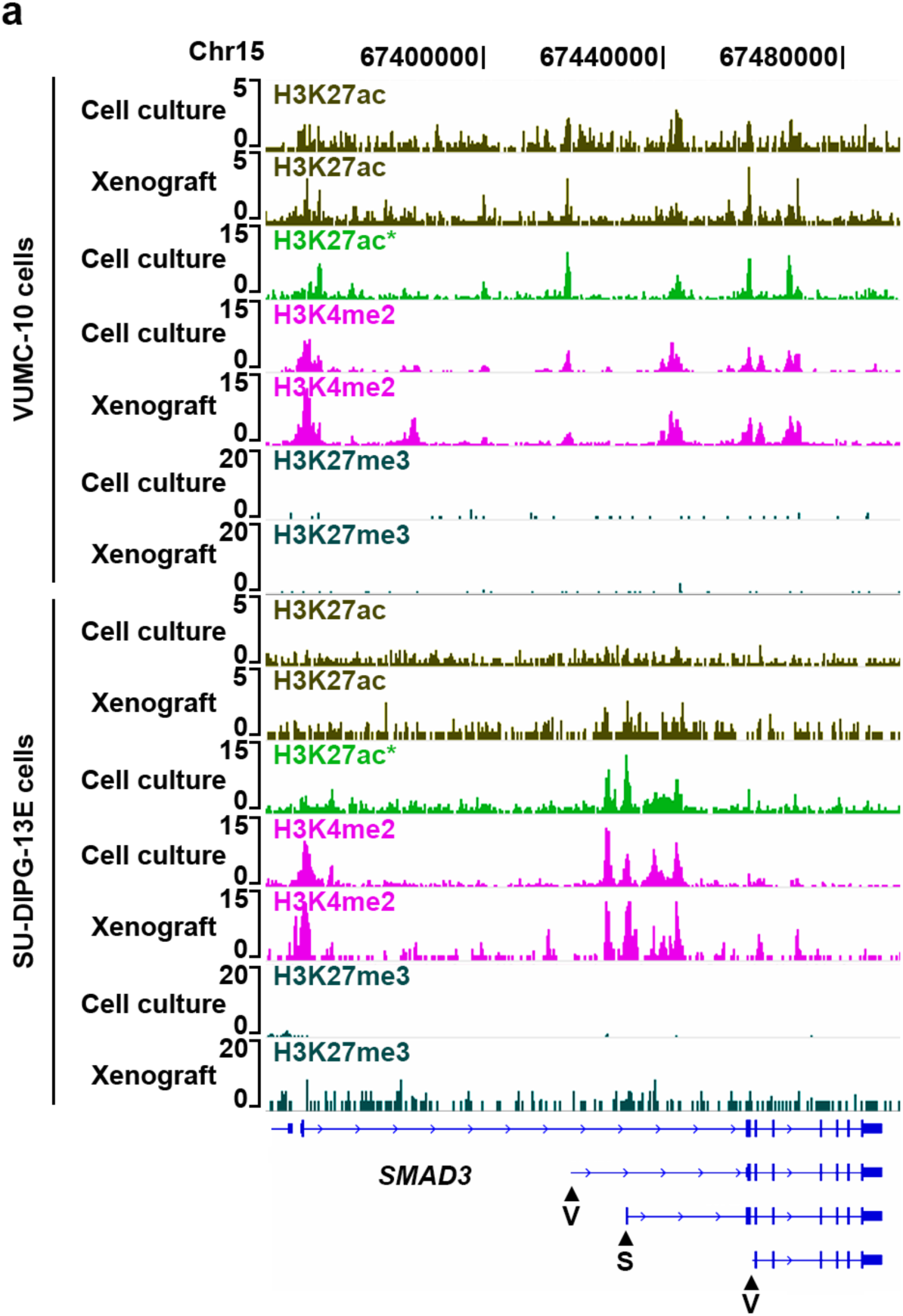
DMG subtype-specific SMAD3 promoter activities. (**a**) Genome browser tracks of histone marks profiled by AutoCUT&RUN in VUMC-10 and SU-DIPG-XIII cells at a representative locus (SMAD3) showing the concordance of profiles from cell culture and xenograft samples. The H3K27ac signal in SU-DIPG-XIII cells was noisy, but this issue is antibody specific. For comparison H3K27ac was also profiled manually using an alternative antibody (*). Arrowheads indicate promoters that are predicted to be specifically active in VUMC-10 (V) or SU-DIPG-XIII (S) cells.

**Supplementary Figure 4.**
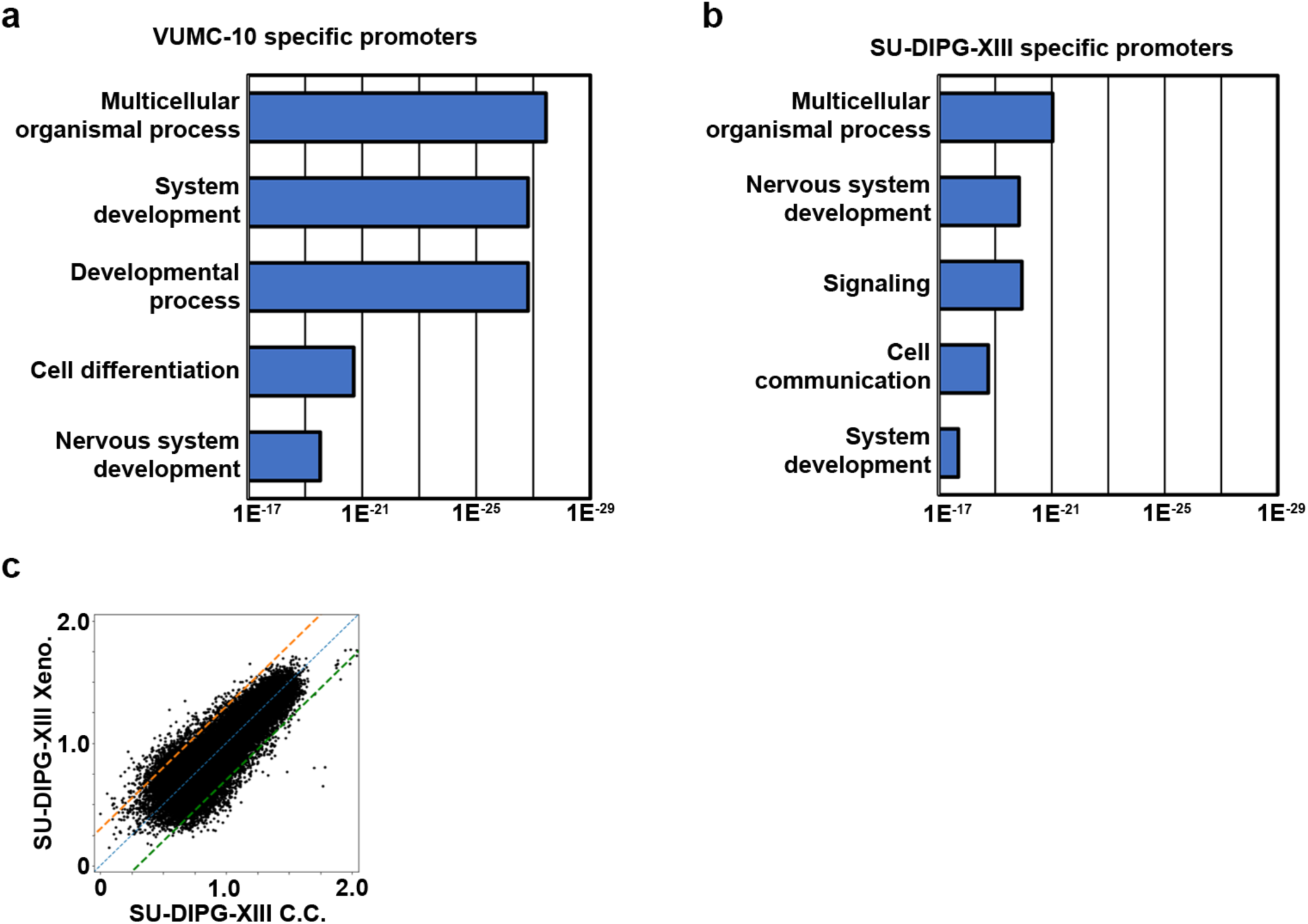
AutoCUT&RUN identifies DMG specific gene regulatory programs. (**a**) GO terms that are overrepresented in the collection of promoters that are ≥2-fold enriched in VUMC-10 cells according to CREAM analysis. (**b**) GO terms that are overrepresented in the collection of promoters that are ≥2-fold enriched in SU-DIPG-XIII cells according to CREAM analysis. (**b**) Scatterplot comparing the promoter CREAM scores of SU-DIPG-XIII cell culture (C.C.) and xenograft (Xeno.) samples. 1,619 promoters have a ≥2-fold difference in promoter chromatin scores between these samples.

**Supplementary Figure 5.**
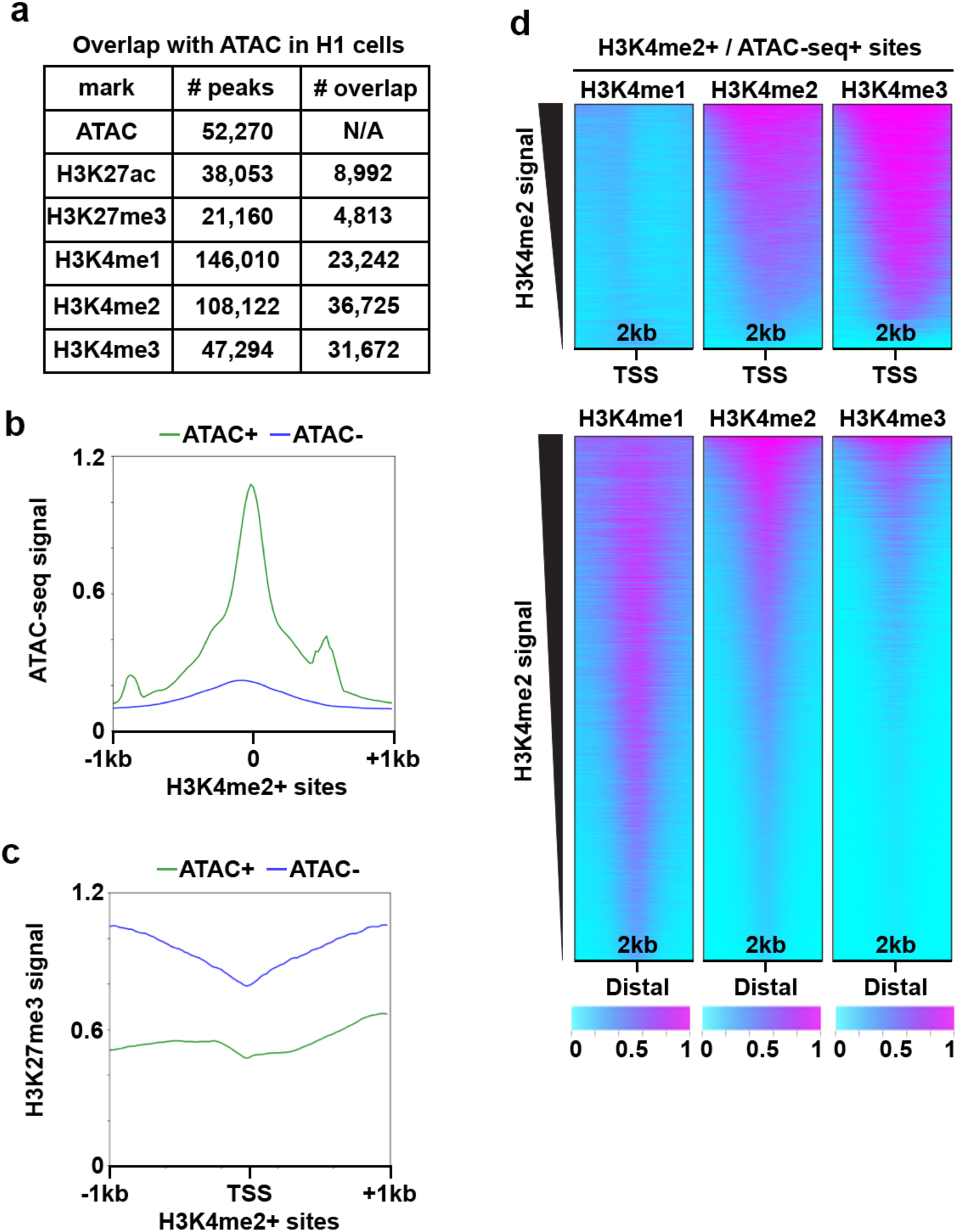
AutoCUT&RUN is a sensitive method to distinguish proximal and distal cis-regulatory elements. (**a**) Table of the overlap of accessible chromatin sites (ATAC-seq peaks) and peaks called on various AutoCUT&RUN profiles of histone marks in H1 cells. (**b**) Mean enrichment of ATAC signal at H3K4me2 peaks that were either called as ATAC+ (green) or ATAC- (blue). (**c**) Mean enrichment of H3K27me3 signal at H3K4me2+ TSSs that were either called as ATAC+ (green) or ATAC- (blue). (**c**) Heat maps showing the distribution of normalized H3K4me1, H3K4me2 and H3K4me3 profiles over all H3K4me2+/ATAC+ TSS and distal regulatory elements (DREs).

